# Rapid whole genome- and annotation-based multilocus sequence typing of the fungal pathogen *Histoplasma capsulatum*

**DOI:** 10.64898/2026.09.27.754713

**Authors:** Arnoud H.M. van Vliet

**Affiliations:** Discipline of Comparative Biomedical Sciences, Section of Infection and Immunity, School of Veterinary Medicine, Faculty of Health and Medical Sciences, University of Surrey, Guildford GU2 7AL, United Kingdom

**Keywords:** *Histoplasma*, multilocus sequence typing, MLST, genome assembly, population structure, fungal pathogen, genome annotation

## Abstract

**Introduction:** *Histoplasma capsulatum* is an important fungal pathogen of humans, especially in immunocompromised patients, and the World Health Organisation has listed *Histoplasma* as a High Priority fungal pathogen. The taxonomy of the genus *Histoplasma* is still under development, but its only species *H. capsulatum* has at least of seven lineages, and a range of sublineages.

**Gap Statement:** Allelic multilocus sequence typing (MLST) has been very successfully applied for typing and epidemiology of pathogenic bacteria, but its adoption for fungal pathogens is more limited. Genome sequence-based MLST approaches promise much a higher resolution for comparative phylogenetics, but there are only a small number of genome assemblies available for *H. capsulatum*.

**Aim:** To develop and validate genome-based MLST typing for *H. capsulatum*.

**Methodology:** A total of 406 genome assemblies were generated from publicly available Illumina short-read sequencing samples. Selected genome assemblies were subsequently used to generate MLST schemes with chewBBACA, based on whole genome sequences (wgMLST), NCBI-obtained annotations (annMLST) and annotations obtained with Funannotate (fanMLST), and used for lineage identification, and comparative analyses of population structure.

**Results:** Genome assemblies were successfully generated for 406 *H. capsulatum* samples, with BUSCO completeness scores of over 98%. A subset of 60 genome assemblies was used to generate and validate MLST schemes based on the 60 genome assemblies, 5 genome annotations obtained from Genbank, and 13 genome annotations generated using Funannotate, resulting in schemes with > 8,000 markers. Each of the MLST schemes allowed reproducible separation of all 7 lineages and 6 *H. capsulatum* Suramericanum sublineages, without the need for high-performance clusters or powerful computing equipment. Comparison of the different schemes using tanglegrams and scatter plots showed good reproducibility (r^2^ values of 0.8-1.0). Comparison of total sizes of genome assemblies showed significant lineage-dependent size differences, with the Suramericanum, LAmB, India, Capsulatum and Mississippiense lineages averaging a total genome size of 30-33 million bp, and the Africa and Ohiense lineages averaging at 39 million bp.

**Conclusion:** Genome sequence- and especially genome annotation-based MLST analyses are a promising approach that can be adapted as a rapid, high-throughput method for genomic clustering and epidemiological tracing of pathogenic fungi.

## INTRODUCTION

Fungal infections are causing increasing public health and economic problems worldwide through increased dissemination and their severity, especially in vulnerable groups [1]. One of these fungal pathogens of concern is *Histoplasma*, which is an ascomycete fungus that causes histoplasmosis in humans and animals. *Histoplasma* are dimorphic fungi, with an environmental mold stage that produces infectious conidia, which upon entry into the lungs convert to a yeast stage which can disseminate from the lungs to a systemic infection [2]. *Histoplasma* is commonly found in soil, but has also been found in wild animals such as rodents and bats. It has a worldwide distribution, but is endemic in the Americas and Africa. *Histoplasma* can cause severe disease in immunocompromised hosts, such as patients with HIV/AIDS, on chemotherapy or those on immunosuppressant drugs [3]. In 2022, the World Health Organisation included *Histoplasma* on its fungal priority pathogens list in the High Priority group, together with the *Mucorales* and several *Candida* species [4].

The taxonomy of the genus *Histoplasma* is under continuous development. Initially, three *Histoplasma capsulatum* major lineages were recognised: *H. capsulatum* var. *duboisii* (*H. duboisii*) causing African histoplasmosis with cutaneous and subcutaneous granulomatous lesions, the “New World” *H. capsulatum* var. *capsulatum* (*H. capsulatum*) causing pulmonary and disseminated histoplasmosis, and the horse pathogen *H. capsulatum* var. *farciminosum* (*H. farciminosum*) [5]. Subsequent molecular investigations based on four protein-coding genes concluded that *H. farciminosum* was not a monophyletic species, but more importantly, showed that *H. capsulatum* could be divided into multiple genetic lineages [6]. Later studies based on genome sequencing studies have supported this subdivision of genetic lineages, with major lineages being from Latin America (LAmA and LAmB), Africa (Africa/duboisii), North America (NAm1 and NAm2), and worldwide (H81/Panama). Some of these have subsequently been proposed as separate *Histoplasma* species (*H. mississippiense*, *H. ohiense*, *H. suramericanum*) [7, 8], but *H. ohiense* and *H. mississippiense* are currently labeled as invalid names in the NCBI Taxonomy database. The current status of *Histoplasma* taxonomy is summarised in Table S1; in this study, *H. capsulatum* lineage and sublineage names will be used without use of disputed species names.

The use of genome sequencing has enabled the study of *Histoplasma* genetic diversity to a much higher resolution. The reported genome size of *H. capsulatum* is between 30-38 million basepairs (Mbp) based on 12 genome assemblies in Genbank, although only a few genomes have been assembled at chromosome level [9]. Genome-level epidemiology studies have so far focused on mapping to reference genomes followed by either use of single nucleotide polymorphisms (SNPs) or concatenated genome-wide alignments [10–14]. A weakness of such approaches is the need for a reference genome scaffold and mapping.

Multilocus sequence typing (MLST) has been used extensively with pathogenic bacteria, initially with seven genes [15], but has progressed to core genome and whole genome-based schemes available via websites such as PubMLST and databases such as BigsDB [16]. In MLST, gene sequences are converted into allele numbers with sequence variations resulting in new allele numbers. Each genome is separately analysed and converted in a collection of allele numbers, representing a genomic barcode. Freely available software packages such as chewBBACA now allow for rapid development and analysis of genome-based MLST schemes [17, 18], but their use so far has been limited to prokaryotic organisms.

One of the new 2030 Global Targets for Histoplasmosis is “understanding pathogen diversity and hist interactions” [3], and availability of whole genome MLST schemes would be beneficial to achieving this target. In this study, *de novo* genome assemblies have been generated for 406 previously described *Histoplasma* samples, and the chewBBACA software has been used to generate multiple genome-wide MLST schemes based on either whole genome or annotated features. After validation by comparison with previously used typing schemes, these MLST schemes have subsequently been used for a rapid large-scale comparative analysis of *Histoplasma* genomic diversity.

## MATERIALS AND METHODS

### Data acquisition

Information on available genome assemblies and genome sequencing reads were obtained from the National Center for Biotechnology Information (NCBI), using the NCBI Genome (https://www.ncbi.nlm.nih.gov/datasets/genome/) and NCBI SRA (https://www.ncbi.nlm.nih.gov/sra/) websites. Search terms used were “Histoplasma” and “Histoplasma capsulatum”. A total of 12 *H. capsulatum* genome assemblies were downloaded with ncbi-genome-download version 0.3.3 (https://github.com/kblin/ncbi-genome-download), while the FASTQ files for 403 *H. capsulatum* and 3 *H. capsulatum* var. *farciminosum* (subsequently referred to as *H. farcimosum*) samples were downloaded using the prefetch and fasterq-dump utilities of the SRA toolkit version 3.2.1 (https://github.com/ncbi/sra-tools). One samples each of *Blastomyces dermatitidis*, *Blastomyces parvus*, *Emergomyces* (*Emmonsia*) *pasteuriana* and *Paracoccidioides brasiliensis* was included to serve as outgroups for phylogenetic tree rooting [11]. All accession numbers are provided in Table S3. Genome annotations of 5 selected *H. capsulatum* genome assemblies were downloaded from NCBI Genome (Table S4). Metadata were obtained from NCBI databases as well as from publications that described the isolates.

### Genome assembly, assembly metrics and genome annotation

Genome assemblies were generated *de novo* using Shovill version 1.1.0 (https://github.com/tseemann/shovill) using the SPAdes assembler version 3.15.5 [19]. Reads were subsampled to a maximum depth of 100×, polished using Lighter version 1.1.3 [20], overlapping reads were stitched using FLASH version 1.2.11 [21], assembled with SPAdes with a minimum contig size of 500 nt and a minimum contig coverage of 2.0×, with post-assembly correction disabled. FASTA headers were made specific for each assembly using a combination of the linux command sed and the Seqfu program version 1.25 [22] (Table S6). The assembly metrics for all genome assemblies (those downloaded and those generated for this study) were determined using Quast version 5.0.2 [23] and are included in Table S2. Genome completeness was estimated using BUSCO 6.1.0 [24], with the 5,439 markers of the ajellomycetaceae_odb12.2 lineage database, searched with the Miniprot version 0.18-r281 gene predictor [25]. Three genome assemblies (SRR29905408, SRR31893259 and SRR31893306) were excluded from further analysis, as they were likely to represent contaminated/mixed samples with genome sizes of >65 Mbp, far exceeding the genome sizes observed in the other samples (maximum 49 Mbp, average genome size 35.4 ± 4.1 Mbp, Figure S1). Another 98 genome assemblies and 4 downloaded assemblies were either mutants derived from isolate G217B or its *ura5*^-^ derivative WU15 [9, 26] or duplicate assemblies of isolates G217B, H88, WU24, G184AR, G186A or G186AR. For the duplicates, the best quality assembly based on Quast metrics was selected for full analyses, and the others used for assessing the reproducibility of the wgMLST approaches.

### Generation of multilocus sequence typing (MLST) schemes based on whole genome sequences and annotated features

The chewBBACA software suite version 3.5.4 [18] was used for all MLST analyses in this study (https://chewbbaca.readthedocs.io/en/latest/index.html). A training file was generated with Pyrodigal version 3.6.3 [27, 28] using the complete *H. capsulatum* H88 genome assembly (Africa lineage, accession number GCA_017310615.1) with a minimum open reading frame (ORF) length of 300 and 500 nt, respectively. Three types of MLST schemes were generated, each with a minimum ORF length of 300 or 500 nt: whole genome MLST schemes (wgMLST), Genbank annotation-based gene MLST schemes (annMLST) and Funannotate-based gene MLST schemes (fanMLST). The wgMLST schemes were generated using a subset of 60 *H. capsulatum* genome assemblies representing six recognised lineages and seven sublineages: Africa, Capsulatum (*sensu stricto*), India, LamB, Mississippiense, Ohiense, and the Suramericanum sublineages Amazon-1, Amazon-2, Mz5-like, NEB, RJ, Sura, and 27-14 [10, 11], with the list of genomes used indicated in Tables S2 and S3. The Funannotate schemes were based on gene calling using the Funannotate version 1.8.17 Docker installation [29] without Genemark installed, and using the pretrained Augustus dataset for *H. capsulatum*. One genome each for each of the 13 lineages/sublineages/ungrouped isolate was annotated (Table S4), and the genes (exons and introns combined, from startcodon to stopcodon) extracted using AGAT version 1.6.1 (https://github.com/NBISweden/AGAT). Finally, 5 *H. capsulatum* annotations were downloaded from NCBI Genome representing two capsulatum *sensu stricto* genomes, and one each of the Africa, Ohiense and Mississippiense lineages, with the genes (exons and introns combined) extracted (Table S4). MLST schemes were generated using the CreateScheme command for each minimum ORF length with translation table 1 selected, tested against the 60 genomes representing the wgMLST scheme, against the 4 outgroup genomes, and against the 98 duplicate/derivative genomes and 11 comparator genomes to assess robustness of the assignment and schemes. Subsequently, the full unique dataset of 310 *H. capsulatum* and 3 *H. farciminosum* genomes was tested against the three schemes and two minimum gene length settings using the AlleleCall command of chewBBACA. Schemes thus generated were stored as tar.gz archives and are available via Figshare (doi: 10.6084/m9.figshare.33946246) and Zenodo (doi: 10.5281/zenodo.22865196), which also contains the Funannotate annotations of the 13 genomes.

### Generation and comparison of MLST and core genome SNP-based phylogenetic trees

Allele calls of MLST schemes as produced by chewBBACA were used to build phylogenetic trees using GrapeTree version 2.2 [30] using the RapidNJ algorithm. The four outgroup genomes were initially used to confirm that the India sublineage of *H. capsulatum* can be used for rooting trees [10, 11] and were not included in further comparisons. For comparison purposes, trees were also generated for the 60 wgMLST scheme genomes based on core genome single nucleotide polymorphisms (SNPs), as determined using ParSNP version 1.2 [31] using a k-mer setting of 31 (-a 31) and the PhiPack setting activated (+x) [32]. SNPs were based on a cluster coverage of 37.5% of the respective genomes included (range 29.2-43.3%). Phylogenetic trees thus generated were rooted based on the India lineage, visualised using Figtree version 1.4.2 (https://github.com/rambaut/figtree/) and tree orders were converted into numerical values using TreeGraph v2.0 [33] and are provided in Table S5. Trees were compared using tanglegram approaches, with the SNP-based tree used as the gold standard, plotted as scatter plots with the r^2^ value determined to assess the correlation between two trees. Metadata (year, isolation source and country) were obtained from publications [6, 8, 10–13] and from the metadata associated with the NCBI Biosample code of the samples, and are provided in Table S3. It needs to be noted that year of isolation did not always match between the information from NCBI Biosample and publications, and hence may be less reliable.

### Data description

All Linux commands used for training file generation, MLST scheme generation, allele calls and phylogenetic tree generation are provided in Table S6. Genome assemblies have been made available via the European Nucleotide Archive (https://www.ebi.ac.uk/ena) in Bioproject number PRJEB125984, with the individual accession numbers provided in Table S3. The genome assembly-and genome annotation-based MLST schemes have been made available via Zenodo (doi: 10.5281/zenodo.22865196) and Figshare (doi: 10.6084/m9.figshare.33946246), together with Excel-files of Tables S2-S5 and a flat text version of Table S6.

## RESULTS

### De novo assembly of Histoplasma capsulatum genomes

A search of the NCBI SRA and NCBI Genome databases highlighted a contrast between the number of *Histoplasma* samples with Illumina sequencing reads available (406) and the limited number of genome assemblies available (12). To address this paucity of *H. capsulatum* genome assemblies, the 406 *Histoplasma* samples were used for genome assembly. A minimum contig size of 500 nt was chosen to avoid contamination with small contigs which are difficult to assess for contamination. Figure S1 shows the genome sizes of the individual assemblies compared to the genome assemblies sourced from NCBI Genome, with detailed information per genome assembly provided in Table S2. The average size of the 12 *Histoplasma* genome assemblies from NCBI Genome was 35.88 ± 5.12 Mbp, with genome sizes ranging between 30.48 - 46.12 Mbp, whereas the 406 new genome assemblies had an average genome size of 35.66 ± 5.06 Mbp, with genome sizes ranging between 26.48 - 74.96 Mbp. Subsequent analysis of assembly metrics (size, number of contigs, N50) led to the exclusion of the three largest genome assemblies (Figure S1) as these were >20 Mbp larger than the reference genome assemblies from NCBI Genome. BUSCO was used to assess completeness of the genome assemblies, with the NCBI Genome assemblies scoring an average of 98.5% (range 93.5-99.7%), the new assemblies scoring an average of 99.0% (range 69.0-99.7%), with 2 and 5 assemblies scoring below 95%, respectively; these were mostly more fragmented genomes with lower assembly metrics. Analysis of sample metadata highlighted that 17 of the *de novo* assembled genomes were independent duplicates of already sequenced reference isolates, whereas 81 were derivatives of *H. capsulatum* G217B or WU15 (G217B *ura5*^-^) [9, 26]. Separation of these exclusion, derivatives and duplicates left 305 *de novo* genome assemblies with an average genome size of 34.79 ± 4.32 Mbp, with genome sizes ranging between 26.48 - 49.19 Mbp (Figure S1). The G217B/WU15 derivatives showed two different assembly sizes: 36-38 Mbp and 31-33 Mbp, consistent with the reported chromosomal alterations in a subset of these derivatives [9, 26]. Other duplicates/derivatives showed approximate matching of genome assembly sizes.

### Generation of multilocus sequence typing schemes for *Histoplasma*

An initial selection of 60 genomes previously assigned was made representing all 13 lineages and sublineages of *H. capsulatum* [10, 11]. A phylogenetic tree was generated from core genome SNPs, which showed the predicted grouping and subgrouping, ordering the Suramerican sublineages (e.g. Sura, Mz5-like) together, as well as North-American lineages (e.g. Ohiense, Mississippiense), global lineages (e.g. Africa, Capsulatum), with the India lineage as the most remote group (Figure 2A, Figure S2). The chewBBACA software suite [18] was used to generate whole genome MLST (wgMLST) schemes, with the minimum ORF length fixed at 300 and 500 nt as setting for the prokaryotic gene prediction module Pyrodigal, resulting in 52,068 and 17,117 alleles, respectively. Analysis of the annotations of five NCBI Genome-obtained genome assemblies showed that only 32.3-42.4% of their genomes represented coding genes (Table S4). To assess whether the non-coding sequences affected the wgMLST schemes, two additional sets of MLST schemes with 300 and 500 nt minimum ORF length were generated, one based on the annotated NCBI Genome-obtained assemblies (annMLST) and one based on a representative genome of each of the 13 lineages and sublineages, annotated with Funannotate (fanMLST). Predicted genes (exons and introns combined) covered 33.8-45.5% of the total genome sequences, similar to the NCBI Genome-annotated assemblies (Table S4), and contained 14,963 and 7,958 alles (annMLST), and 15,849 and 8,979 alleles (fanMLST), respectively. The phylogenetic trees based on the MLST allele assignments were rooted with outgroup fungi closely related to *Histoplasma* (*Blastomyces*, *Emergomyces* and *Paracoccidioides* [10, 11]), which consistently showed the India lineage as most closely to these outgroups, and this lineage was used to root the subsequent trees (Figure S2).

**Figure 1.**
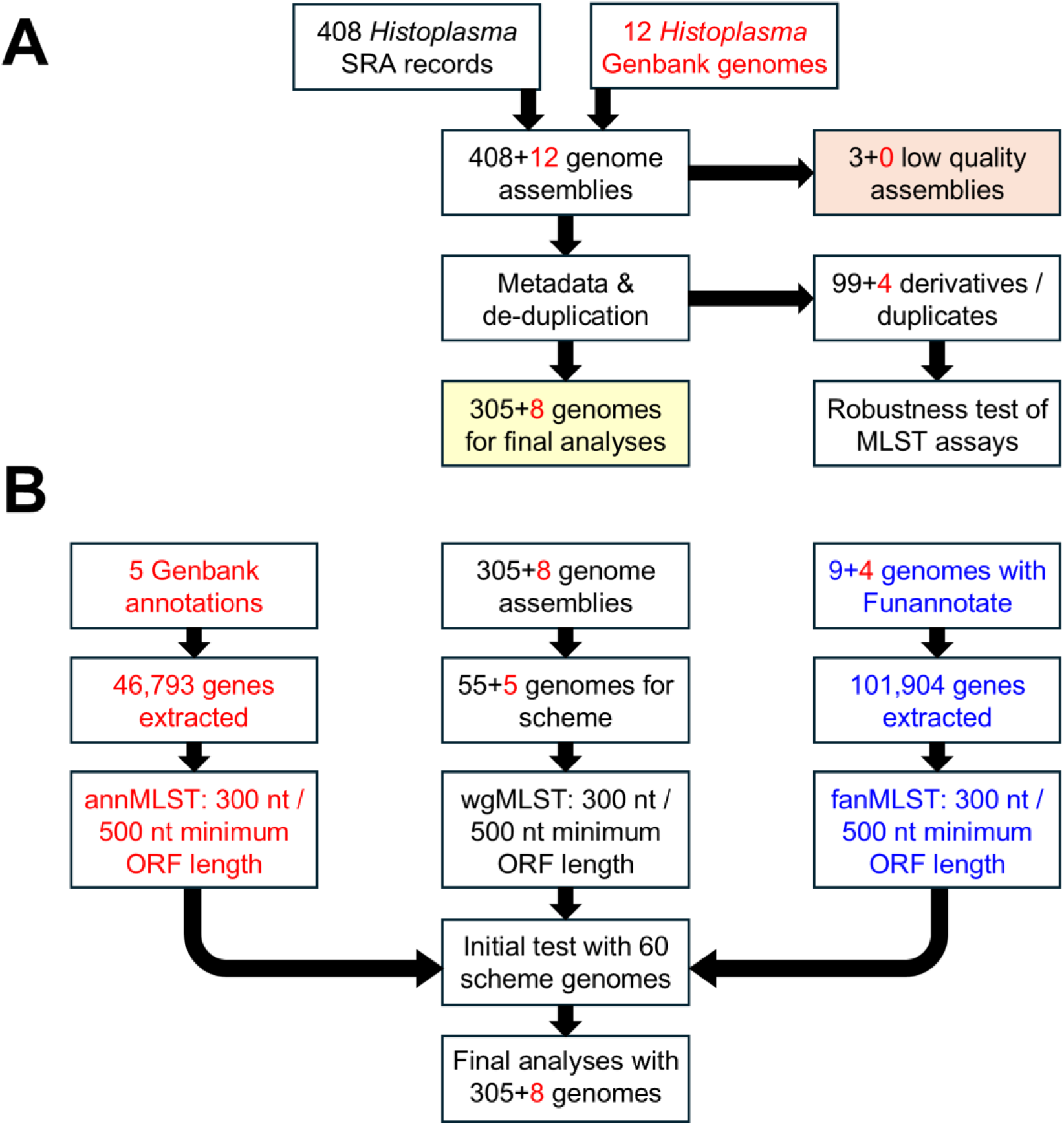
Schematic overview of the setup of this study, with the number of genomes used in each step indicated. Numbers in red indicate previously assembled *H. capsulatum* genomes available from NCBI Genome (Genbank). Part A shows the division of samples, whereas part B shows the generation of genome- and annotation-based MLST schemes and subsequent analyses.

**Figure 2.**
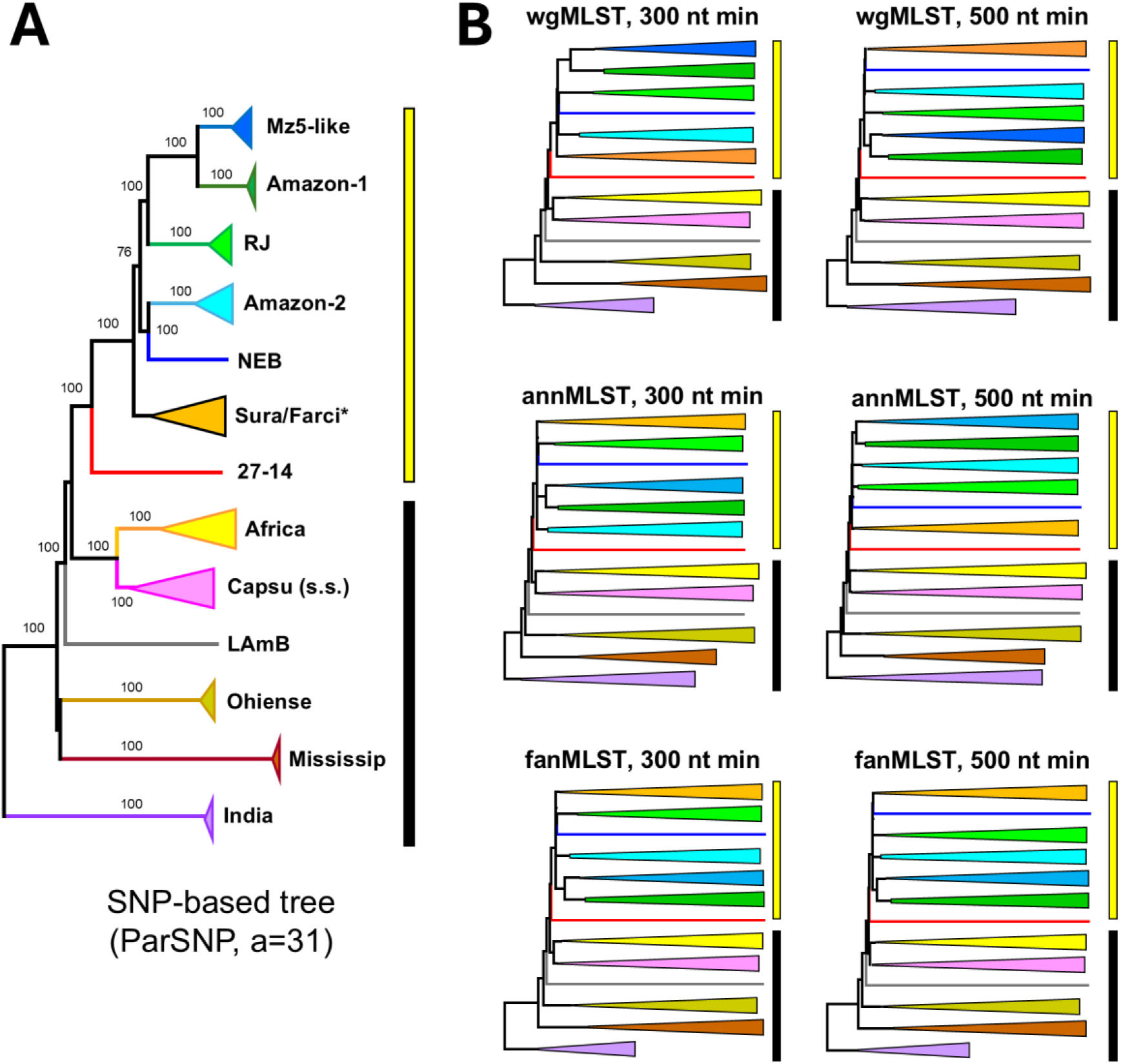
Schematic trees comparing *H. capsulatum* genome population structure by core genome SNPs (ParSNP) versus MLST schemes based on whole genome sequences (wgMLST) and schemes based on annotated features (annMLST and fanMLST). The 300/500 nt min indicates the minium ORF length set for the Pyrodigal part of the chewBBACA pipeline. The lineages and sublineages are indicated by colour and text in the ParSNP tree, with the yellow vertical bar marking the *H. capsulatum* var. *suramericanum* sublineages, and the black bar all the other lineages for easy visual marking. The numbers on the ParSNP tree are bootstrap values as reported by ParSNP. All trees are rooted on the India lineage of *H. capsulatum*. Tanglegram trees comparing the ParSNP tree with each of the MLST trees are provided in Figure S3, with the order of specific samples provided in Table S5.

### Validation of the six MLST schemes for *H. capsulatum*

The six MLST schemes were subsequently tested using the 60 selected reference genomes (Figure 2) and compared to the phylogenetic tree based on core genome SNPs. The wgMLST, annMLST and fanMLST schemes all correctly clustered and separated the 13 lineages and sublineages, and also showed similar order of lineages and sublineages. All Suramerican sublineages clustered together, as did the global and North-American lineages and India lineage [10, 11]. There was variation in the order of sublineages and individual genomes within the major lineages, such as the relative positions of the Amazon-1 and Amazon-2 minor sublineages compared to the Sura, RJ and Mz5-like lineages, but tanglegram comparisons of the core genome SNP tree and the three MLST schemes showed strong conversation of overall tree architecture based on r^2^ values of 0.79-0.99 in scatter plots (Figure S3). All six schemes gave similar performance and resolution with the 60 reference genomes.

The genomes derived from G217B/WU15 and duplicate assemblies were used to further test the MLST schemes and the reproducibility of the genome assembly process. The genomes were from 4 different lineages (Ohiense, Mississippiense, Africa and Capsulatum), and all genomes clustered with the comparator genome, with only minor differences (Figure S4), demonstrating the reproducibility of the analysis methods used, as well as the comparable quality of the newly assembled genomes compared to the ones available from NCBI Genome/ENA. There were no clear differences between the three types of whole genome MLST approaches or the two minimum ORF lengths.

### Analysis of *H. capsulatum* population structure using genome sequence-based MLST schemes

The full dataset of 313 *Histoplasma* genomes was analysed using the three genome sequence-based MLST schemes, and as with the initial dataset, displayed similar tree topologies with the six schemes (Figures S5, S6, S7). The tree obtained with wgMLST/300 nt minimal ORF length was subsequently populated with the metadata obtained for these samples, which was complete for the isolation source and lacked information for a single sample with regard to the country/continent of isolation (Figure 3). The majority of samples were obtained from humans (285/313, 91%), with the other samples being from pets (17), horses (3), soil (5) and rodents/marsupial (3), likely a consequence of sampling bias due to healthcare requirements. The non-human samples were primarily found in the Mississippiense lineage (pets), the Ohiense lineage (pets and rodents/marsupial) and the RJ sublineage (soil, rodent and pet). The horse samples were all *H. farciminosum*, which clustered with the Sura sublineage.

**Figure 3.**
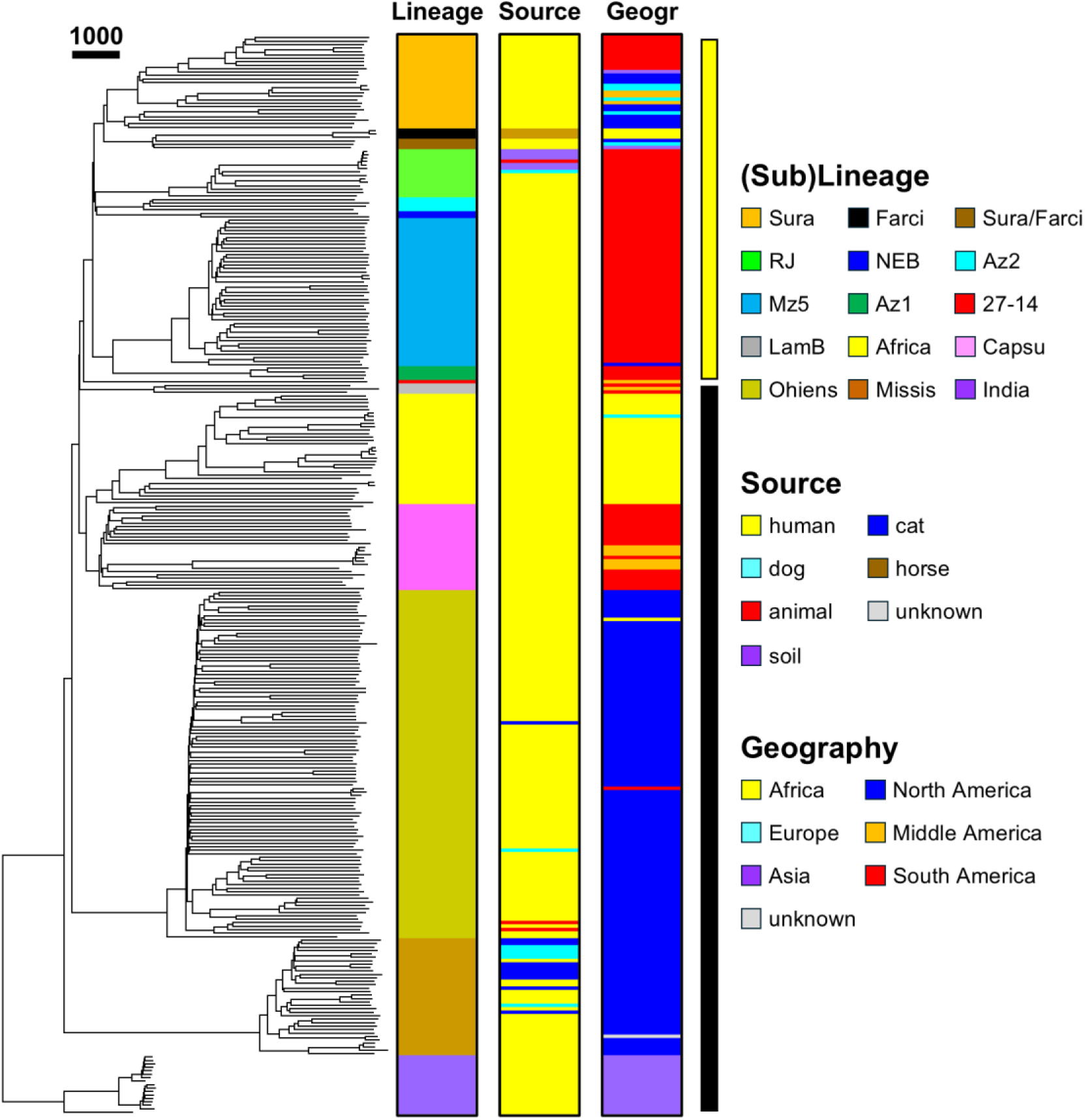
Analysis of population structure of *H. capsulatum* using wgMLST. The tree of 313 *H. capsulatum* genomes was generated using the 300 nt minimum ORF length wgMLST scheme, with the distance bar representing the number of allele differences for a given branch. The bars show the different lineages, isolation source (human/animal/soil) and the geographic sourse of the samples grouped per continent. The yellow vertical bar marks the Suramericanum sublineages, the black bar the other lineages for easy visual marking. The tree is rooted on the India lineage of *H. capsulatum*, with the order of specific samples provided in Table S5.

The geography data were categorised per continent, with Middle America treated as a separate continent for resolution. Most lineages showed the strong geographic associations described before, such as Ohiense (99/101 North-America), Mississippiense (33/34 North-America), India (16/16 Asia) and Africa (31/32 Africa). The Capsulatum lineage was a mix of South-America (19/25) and Middle-America (6/25). Within the Suramericanun lineage, most sublineages showed a similar strong association with continents, with Mz5-like 42/43 from South-America, RJ being 14/14 from South-America and the two Amazon sublineages both 4/4 from South-America. The only sublineage with significant geographic variation was Sura, which had a South-American cluster (10) and a mixed cluster of samples from North-America (10), Middle-America (3), Europe (5) and Asia (2). This latter cluster also contained the *H. farciminosum* samples from Africa (3).

### Total genome size varies between *H. capsulatum* (sub)lineages

With all the samples assigned to a lineage or sublineage, the total genome assembly sizes were plotted accordingly (Figure 4). There was significant variation in total genome size between the (sub)lineages, which was not related to the order in the phylogenetic tree. The Africa and Ohiense lineages had an average genome size of ∼39 Mbp with a range between 36-44 Mbp (Africa) and 36-49 Mbp (Ohiense), while most other (sub)lineages had average genome sizes between 29-33 Mbp, with Mz5-like ranging between 31-40 Mbp. Comparison of *H. farciminosum* genomes (44-49 Mbp) and the three Sura genomes clustering with *H. farciminosum* (Figure 3) suggested that the former may also be *H. farciminosum*, as their genome sizes are much larger (40-46 Mpb) than the other Sura sublineage genomes (30-35 Mbp). These have been labeled “Sura/Farci” in Figures 3 and 4 and Tables S2 and S3.

**Figure 4.**
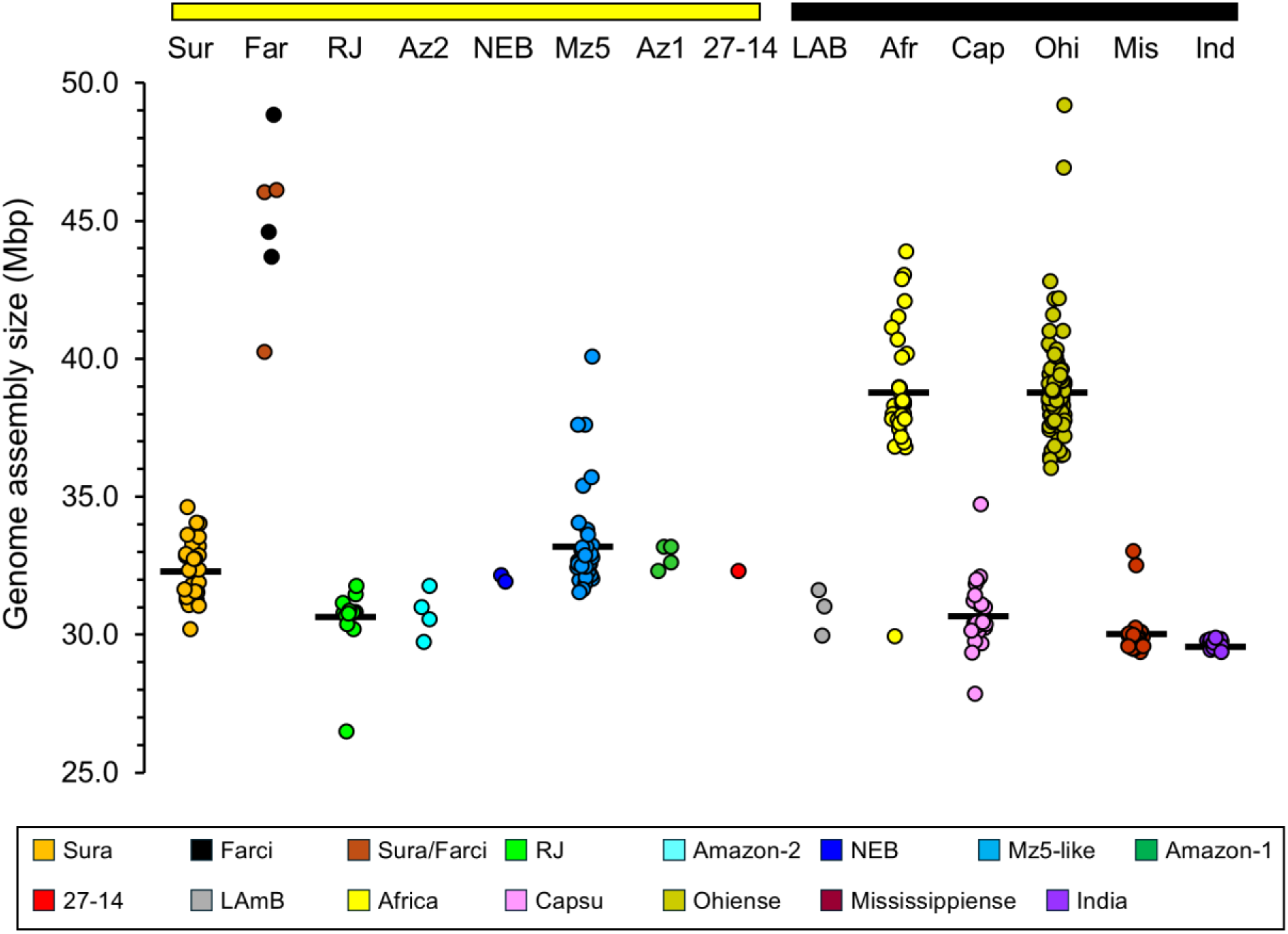
Total sizes of *H. capsulatum* genome assemblies expressed as millions of basepairs (Mbp), divided per (sub)lineage. The yellow horizontal bar at the top marks the *H. capsulatum* var. *suramericanum* sublineages, the black bar the other lineages for easy visual marking. Abbreviated names for lineages are shown at the top. The average assembly size is indicated by a horizontal bar for (sub)lineages with >10 assemblies. Far/Farci indicates Farciminosum, Sura/Farci the putative Farciminosum samples.

## DISCUSSION

At the start of this study, only 12 genome assemblies were available for *Histoplasma capsulatum*. In this study, an additional 403 genome assemblies have been generated from publicly available Illumina sequencing datasets, and after deduplication using metadata, these have been used for generating whole genome/genome annotation-based MLST schemes, followed by validation of these schemes by comparison with prior population structure analyses and lineage distribution. This study is the first to describe use of genome-based MLST schemes for fungal genomes. Gene-based MLST has been used for *Histoplasma* and other pathogenic fungi, but use a limited number of genes (e.g. between three to six for *H. capsulatum*, e.g. [6, 34]) which give limited resolution and information. Other molecular typing methods previously used are ITS and rRNA-based [35], while genome-based methods have focused on single nucleotide polymorphisms (SNPs) and concatenated genome-wide alignments [10–14]. These utilised the Illumina reads rather than genome assemblies, which does require more sophisticated analysis methods and availability of computing power, wich can be an issue for more resource-limited areas, such as those where *H. capsulatum* is endemic. The Shovill genome assembly software and the chewBBACA wgMLST program do not require particularly high-powered computing facilities, although they will run much faster on more powerful systems. For example, running chewBBACA with the 60 *H. capsulatum* genomes used for Figure 2 on a Ubuntu virtual computer with 4 cores and 48 GB of RAM took 48 minutes for the FanMLST_0300 scheme (15,849 alleles), and 20 minutes for the fanMLST_0500 scheme (8,979 alleles). Conversely, use of SNPs does not only require large amounts of memory once increasing the number of genomes, but also requires agreement of which positions to use for comparative purposes, and as such may not be easy to standardise internationally, and will potentially be inflexible with regard to new lineages.

To date, genome-based MLST analysis has been primarily applied to the haploid, small genomes of bacteria. In bacterial genomes, there are no introns and hence no splicing, making identification of open reading frames relatively straightforward. This is achieved by the Prodigal/Pyrodigal program [27, 28], which uses a training file to identify the most likely start and stop signals, and these are used for generation of MLST schemes and their subsequent application. In addition, bacterial genomes have a very high density of coding sequences (up to 90%), which is not necessarily the case for eukaryotic genomes. As shown in Table S4, the *H. capsulatum* annotation covers about 40% of the genome, and this includes the introns, with the rest of the genome containing non-coding sequences such as transposable elements [36]. This is likely to lead to Prodigal/Pyrodigal overestimating the presence of putative coding regions. This is a potential weakness of the approach used here, and future studies could look into the application of eukaryotic gene prediction tools such as those used in software such as BUSCO [24], which can utilise Augustus [37], MetaEuk [38] and Miniprot [25]. Here this potential problem has been in part addressed by also generating MLST-schemes based on annotated features of *H. capsulatum* (exons and introns combined), either those available from NCBI Genome and those *de novo* generated by using Funannotate [29]. While Funannotate is less accurate without transcriptome/proteome data, it does allow selection of which genomes to annotate, as the annotated *H. capsulatum* genomes in NCBI did not cover all the lineages previously identified. Both the annotation and Funannotate-based schemes contained a much lower number of alleles than the whole genome-based ones, about one-third for the 300 nt minimum ORF length, and about half of the 500 nt minimum ORF length setting, without reducing resolution (Figures 2, S3-S7). Hence the Funannotate-based approach may be the most promising for future applications in either *H. capsulatum* or other pathogenic fungi, as it is not dependent on the prior availability of genome annotations, and will give more consistent input material for building MLST schemes.

The genome assemblies generated for this study were based on Illumina short-read sequencing data only, without use of long-read sequencing data, which was not available for most samples. Use of Illumina reads only will invariably lead to fragmented genome assemblies, but this had no discernable consequences on the use of the whole genome/annotation-based MLST schemes, as is shown in Figure S4. All assemblies tightly clustered with the comparator genomes, and while the quality and resolution may not be sufficient for outbreak analyses such as those used for pathogenic bacteria (e.g. listeriosis in [39]), the different lineages and sublineages of *H. capsulatum* separated with sufficient resolution. Like with total genome size, there were differences in the average number of contigs per lineage, with the India lineage showing much higher average number of contigs (3,514 contigs) while having a smaller genome of 30 Mbp (Figure 4), while the Suramerican, Africa and Capsu lineages having a much lower number of average contigs with larger average genomes (952 contigs, 33 Mbp; 1,199 contigs, 39 Mbp; and 735 contigs, 31 Mbp respectively), with the Mississipiense and Ohiense genomes inbetween (1,795 contigs, 33 Mbp and 2,132 contigs, 39 Mbp). Almost all assemblies had a BUSCO completeness score of >98% based on 5,439 markers (Table S2), and as such assembly with Shovill/SPAdes seems to be an acceptable approach for *H. capsulatum*, despite Shovill originally being developed for bacteria and other small microbes, not for fungi. The commands and settings provided in Table S6 should be helpful for future sequencing projects on *H. capsulatum* and related fungal pathogens. The genome assemblies produced for this study will also support future studies that can be aimed at generating new molecular detection assays, and studies on development and detection of antifungal resistance [40]. The latter is still underdeveloped for *H. capsulatum*, with the FungAMR database only containing a Cyp51/Erg11 Y136F mutation for *H. capsulatum* associated with fluconazole/voriconazole resistance, which has low confidence scores [41]. The ChroQueTas tool [41] was used to test the 313 genome assemblies, and only found 3 positive samples, all Mississippiense samples from cat (1) and dogs (2) (not shown). Studying the new genome assemblies in combination with *in vitro* comparative experimental work [8] could allow for the identification of novel antifungal targets and improvement of screening mechanisms.

The samples included in this study represented seven previously described *H. capsulatum* lineages, of which the Suramericanum lineage was further subdivided into seven sublineages (Table S1). The 313 genome assemblies analysed for Figures 3 and S5-S7 represented all those lineages and sublineages, and were delineated by the whole genome and both annotation-based MLST schemes. This allowed visulation of patterns as done in Figure 3, showing the geographic differences per continent and lineage. Most lineages showed strong linkage to a specific continent/subcontinent, such as Mississippiense and Ohiense with North-America, Africa with Africa, India with India, and most Suramericanum sublineages with South-America. The only lineage showing more variation was Capsulatum which combined samples from Middle- and South-America, and the Sura sublineage, which had South-American, North-American, European and African samples, and may actually consist of two or three separate sublineages. Conversely, there was little variation in isolation source, as most samples were derived from human cases, but this is likely to represent a sampling bias. Exceptions were in the Mississippiense lineage where several animal samples (cat, dog, wild rat) were included, and the RJ sublineage where soil samples were included from the early eighties. A more varied source sampling in future studies may well expand the population structure of *H. capsulatum*, and reveal new lineages whose existence has been predicted in earlier work [6].

This work was aimed at demonstrating the feasibility of genome-based MLST analysis for fungal genomes. It should be interpreted as a proof-of-principle study, using *H. capsulatum* as the model fungal pathogen. As indicated, there are several options for making the chewBBACA MLST approach [17, 18] more specific for fungal pathogens and other eukaryotes, but even with its current imperfections it does show good potential to be expanded to a diverse set of fungal pathogens. In conclusion, this study has shown the feasibility of generating and using rapid whole genome and annotation-based MLST schemes for a fungal pathogen. Future work can focus on generating pipelines for the software to portals like Galaxy [42], the usage of more specialised eukaryotic gene prediction algorithms, and the usage of the genome assemblies for further in-depth genetic analysis, such as lineage-specific detection assays and identification of antifungal resistance markers.

## Data statement

All supporting data, software commands and protocols have been provided within the article or through supplementary data files and/or via Zenodo. Seven supplementary figures and six supplementary tables are available with the online version of this article.

## Data summary

All genome assemblies are available from the European Nucleotide Archive in Bioproject number PRJEB125984. The information on the individual *H. capsulatum* genome assemblies has been included in the Supplementary Information (Tables S2 and S3). The genome assembly- and genome annotation-based MLST schemes have been made available via Zenodo (doi: 10.5281/zenodo.22865196) and Figshare (doi: 10.6084/m9.figshare.33946246), together with Excel-files of Tables S2-S5 and a flat text version of Table S6 for convenient re-use.

## Abbreviations

Mbp: millions of basepairs
Bp: basepairs; nt, nucleotides
MLST: multilocus sequence typing
Wg: whole genome
ORF: open rading frame
SRA: Sequence Read Archive
ENA: European Nucleotide Archive.

## Acknowledgements

The author gratefully acknowledges the makers of the invaluable, freely available bioinformatic software programs that made this research possible, as well as the many uploaders of the sequence data and accompanying metadata.

## Author contributions

A.H.M.v.V. designed the study, acquired data, performed all bioinformatic analyses, interpreted the data, and drafted the manuscript.

