## Supplementary Information (Tables S1-S6, Figures S1-S7) for "Rapid whole genome- and annotation-based multilocus sequence typing of the fungal pathogen *Histoplasma capsulatum*"

This Supplementary Information contains 7 Supplementary Figures (S1-S7) and 6 Supplementary Tables (S1-S6). The genome assemblies created for this study have been uploaded into ENA with accession numbers available in Table S3, and the 6 MLST schemes (wgMLST, annMLST, fanMLST, both 300 and 500 nt minimum ORF length) have been uploaded to Zenodo (doi: 10.5281/zenodo.22865196) and Figshare (doi: 10.6084/m9.figshare.33946246). Excel spreadsheet versions of Tables S2-S5 and a text version of Table S6 are available on Figshare (doi: 10.6084/m9.figshare.33946246) and Zenodo (doi: 10.5281/zenodo.22865196) for easier access/re-use.

### LEGENDS TO SUPPLEMENTARY INFORMATION

**Figure S1.** Total sizes of *Histoplasma capsulatum* genome assemblies expressed as millions of basepairs (Mbp). The first column shows the 305 assemblies included in the main study and the 3 assemblies excluded due to their assembly size too large. The second column shows the 98 duplicate assemblies of WU24, H88, G184/G186 and G217B and the G217B/WU15 derivatives. The WU15_R genome assemblies are specifically indicated in grey as these are smaller the other G217B/WU15 assemblies. The 12 assemblies downloaded from Genbank are shown for comparitive purposes.

**Figure S2.** Rooting of MLST-based phylogenetic trees using outgroup genome sequences from *Blastomyces*, *Emergomyces* and *Paracoccidioides* spp. Only major lineages are shown, and only the

MLST schemes with a minimum ORF length of 300 nt. wg = based on whole genome sequences; ann = based on annotated features obtained from Genbank; fan = based on annotated features identified using Funannotate. Note that in each of the three phylogenetic trees, the India lineage is the closest to the outgroups, and was used to root subsequent trees.

**Figure S3.** Comparison of a core genome SNP-based reference phylogenetic tree with a whole genome- and two annotation-based MLST schemes for *Histoplasma capsulatum*, using the 60 genome dataset from Figure 2. parSNP = core genome SNPs; wg = based on whole genome sequences; ann = based on annotated features obtained from Genbank; fan = based on annotated features identified using Funannotate. The minimum ORF length setting of chewBBACA is indicated as 0300 or 0500. The positioning in the tree is visualised using a tanglegram, and quantified using a scatter plot based on the positions in the respective trees from top to bottom. Colour codes for lineages and sublineages are shown below.

**Figure S4.** Testing of the reproducibility of genome assembly and MLST schemes, using duplicate samples of *H. capsulatum* H88, G184/G186 and WU24, and assemblies of derivatives of *H. capsulatum* G217B/WU15. wg = based on whole genome sequences; ann = based on annotated features obtained from Genbank; fan = based on annotated features identified using Funannotate; parSNP = core genome SNPs. The minimum ORF length setting of chewBBACA is indicated as 300 or 500 nt. The phylogenetic trees were rooted using an lineage India genome. The distance bar in the parSNP tree (0.1) represents the number of nucleotide substitutions per site for a given branch, based on the number of SNPs found in the core genome. The distance bar in the MLST trees represents the number of allele differences for a given branch.

**Figure S5.** Comparison of the effects of minimum ORF length on genome-based MLST schemes for *Histoplasma capsulatum*, using the complete 313 genome dataset from Figure 3. wg = based on whole genome sequences; ann = based on annotated features obtained from Genbank; fan = based on annotated features identified using Funannotate. The positioning in the tree is visualised using a tanglegram, and quantified using a scatter plot based on the positions in the respective trees from top to bottom.

**Figure S6.** Comparison of genome-based MLST schemes for *Histoplasma capsulatum* with a minimum ORF length of 300 nt, using the complete 313 genome dataset from Figure 3. wg = based on whole genome sequences; ann = based on annotated features obtained from Genbank; fan =

based on annotated features identified using Funannotate. The positioning in the tree is visualised using a tanglegram, and quantified using a scatter plot based on the positions in the respective trees from top to bottom.

**Figure S7.** Comparison of genome-based MLST schemes for *Histoplasma capsulatum* with a minimum ORF length of 500 nt, using the complete 313 genome dataset from Figure 3. wg = based on whole genome sequences; ann = based on annotated features obtained from Genbank; fan = based on annotated features identified using Funannotate. The positioning in the tree is visualised using a tanglegram, and quantified using a scatter plot based on the positions in the respective trees from top to bottom.

**Table S1.** Overview of historical and current names of *Histoplasma capsulatum* variants/lineages.

**Table S2.** Quast assembly metrics and BUSCO completeness scores of *Histoplasma capsulatum* genome assemblies used in this study.

**Table S3.** Metadata of *Histoplasma capsulatum* genome assemblies used in this study, including order in the ParSNP and wgMLST trees, accession and biosample numbers, isolation source, year of isolation, country/continent of isolation, and indication of status (duplicate/derivative).

**Table S4.** Information on the number of annotated protein coding genes for *Histoplasma capsulatum* genomes annotated with Funannotate, and annotated *H. capsulatum* genomes downloaded from NCBI Genome.

**Table S5.** Order of samples in the phylogenetic trees shown in Figures 2, 3, S2, S4, S5-S7.

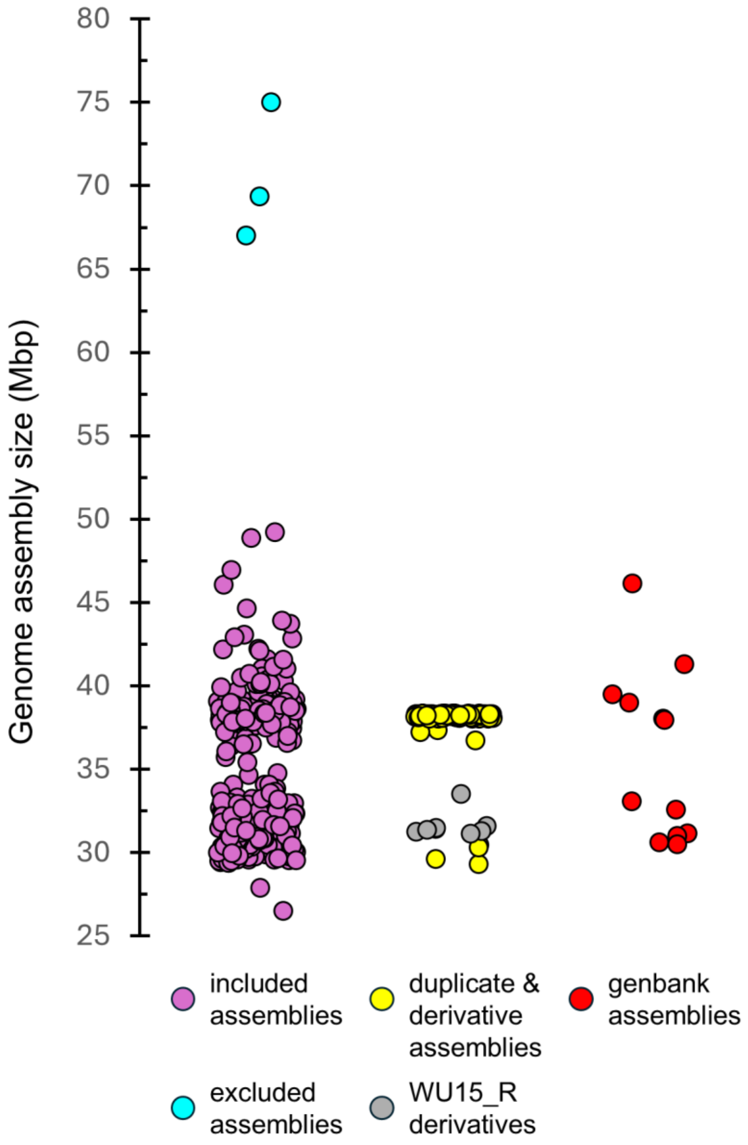


**Figure S1.** Total sizes of *Histoplasma capsulatum* genome assemblies expressed as millions of basepairs (Mbp). The first column shows the 305 assemblies included in the main study and the 3 assemblies excluded due to their assembly size too large. The second column shows the duplicate assemblies of WU24, H88, G184/G186 and G217B, and the G217B/WU15 derivatives. The WU15_R genome assemblies are specifically indicated as these are smaller than their G217B/WU15 versions, confirmed in Heater et al, 2026a, 2026b. The 12 assemblies downloaded from Genbank are shown for comparitive purposes.

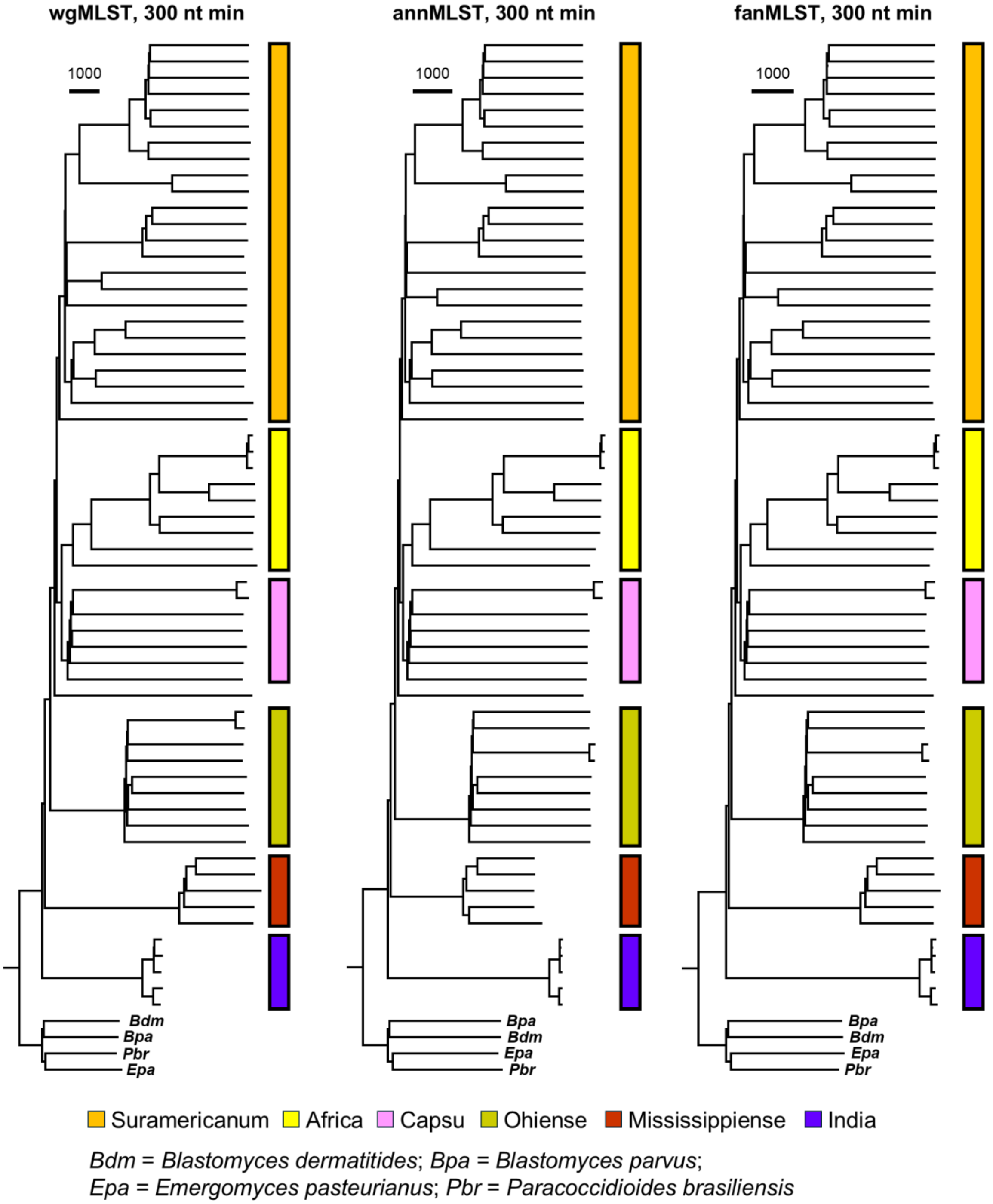


**Figure S2.** Rooting of MLST-based phylogenetic trees using outgroup genome sequences from *Blastomyces*, *Emergomyces* and *Paracoccidioides* spp. Only major lineages are shown, and only the MLST schemes with a minimum ORF length of 300 nt. wg = based on whole genome sequences; ann = based on annotated features obtained from Genbank; fan = based on annotated features identified using Funannotate. Note that in each of the three phylogenetic trees, the India lineage is the closest to the outgroups, and was used to root subsequent trees.

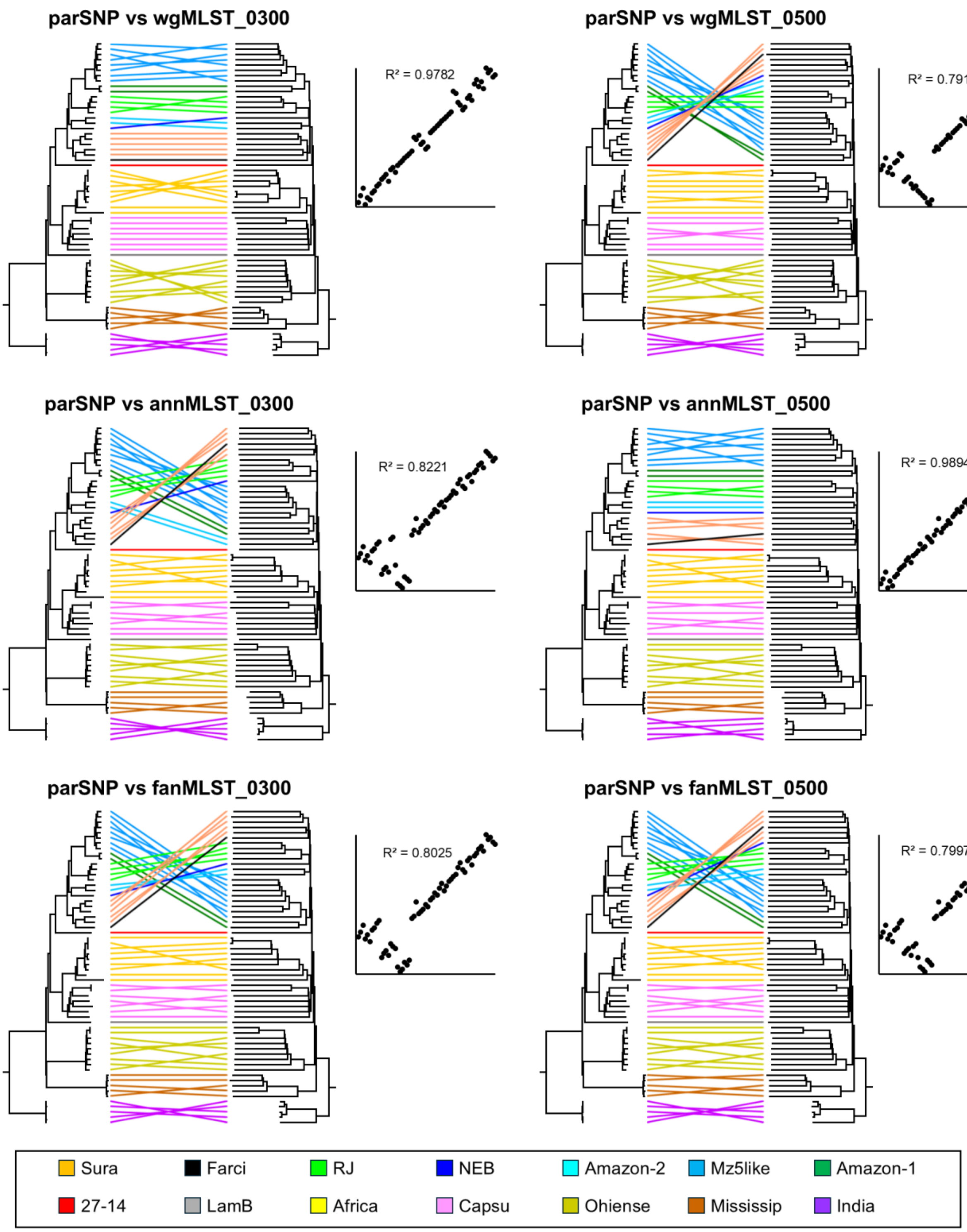


**Figure S3.** Comparison of a core genome SNP-based reference phylogenetic tree with a whole genome- and two annotation-based MLST schemes for *Histoplasma capsulatum*, using the 60 genome dataset from Figure 2. parSNP = core genome SNPs; wg = based on whole genome sequences; ann = based on annotated features obtained from Genbank; fan = based on annotated features identified using Funannotate. The minimum ORF length setting of chewBBACA is indicated as 0300 or 0500. The positioning in the tree is visualised using a tanglegram, and quantified using a scatter plot based on the positions in the respective trees from top to bottom. Colour codes for lineages and sublineages are shown below.

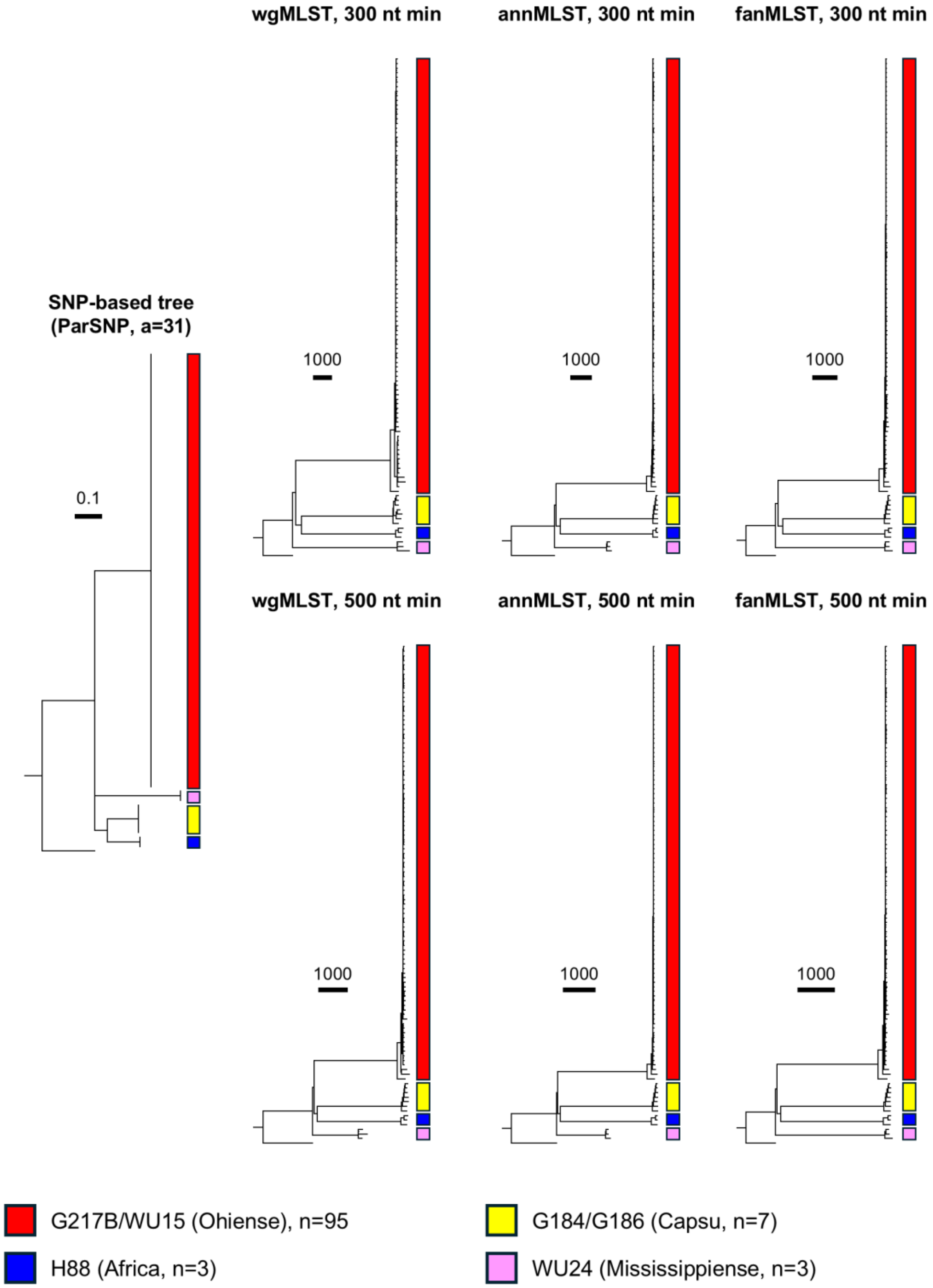


**Figure S4.** Testing of the reproducibility of genome assembly and MLST schemes, using duplicate samples of *H. capsulatum* H88, G184/G186 and WU24, and assemblies of derivatives of *H. capsulatum* G217B/WU15. wg = based on whole genome sequences; ann = based on annotated features obtained from Genbank; fan = based on annotated features identified using Funannotate; parSNP = core genome SNPs. The minimum ORF length setting of chewBBACA is indicated as 300 or 500 nt. The phylogenetic trees were rooted using an lineage India genome. The distance bar in the parSNP tree (0.1) represents the number of nucleotide substitutions per site for a given branch, based on the number of SNPs found in the core genome. The distance bar in the MLST trees represents the number of allele differences for a given branch.

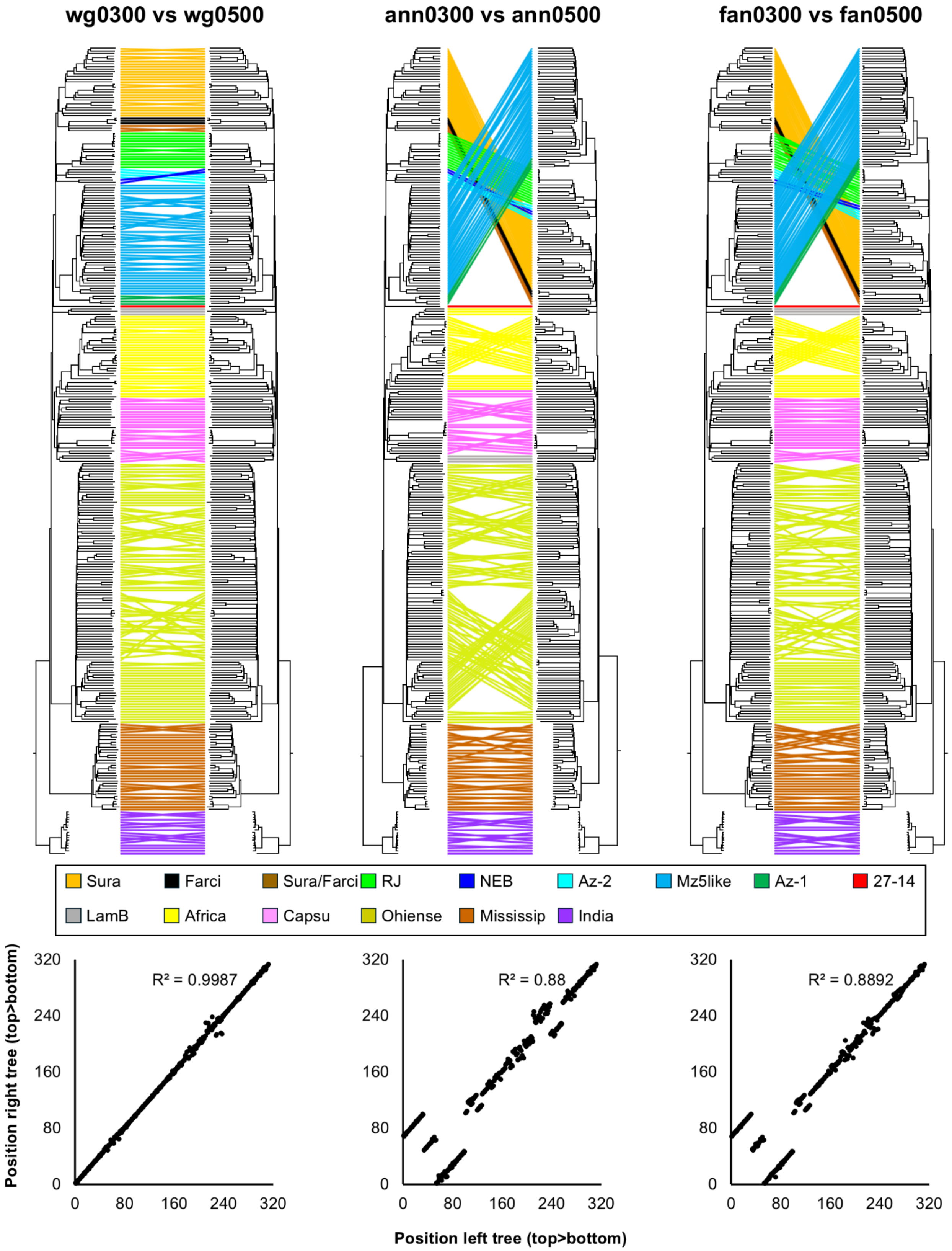


**Figure S5.** Comparison of the effects of minimum ORF length on genome-based MLST schemes for *Histoplasma capsulatum*, using the complete 313 genome dataset from Figure 3. wg = based on whole genome sequences; ann = based on annotated features obtained from Genbank; fan = based on annotated features identified using Funannotate. The positioning in the tree is visualised using a tanglegram, and quantified using a scatter plot based on the positions in the respective trees from top to bottom.

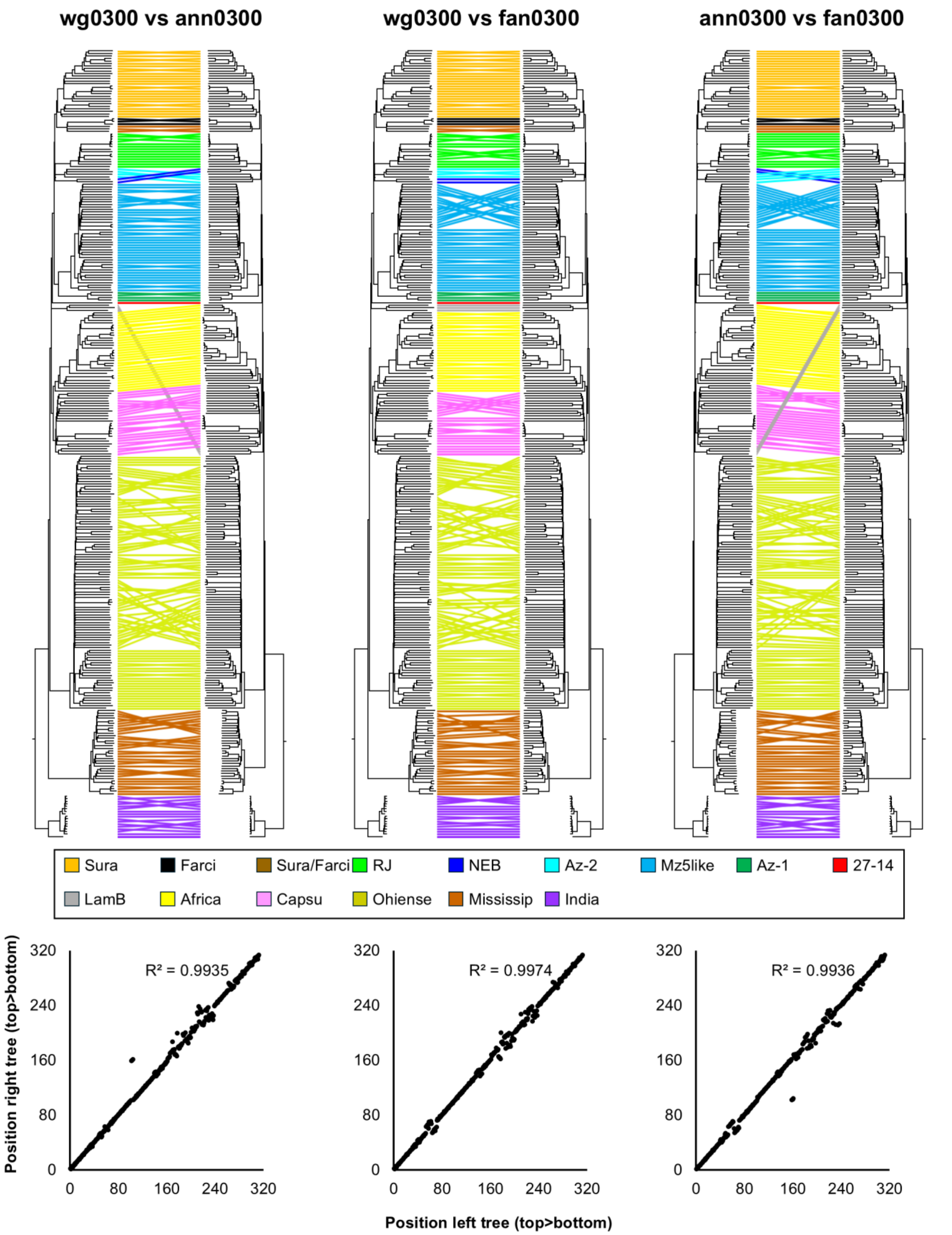


**Figure S6.** Comparison of genome-based MLST schemes for *Histoplasma capsulatum* with a minimum ORF length of 300 nt, using the complete 313 genome dataset from Figure 3. wg = based on whole genome sequences; ann = based on annotated features obtained from Genbank; fan = based on annotated features identified using Funannotate. The positioning in the tree is visualised using a tanglegram, and quantified using a scatter plot based on the positions in the respective trees from top to bottom.

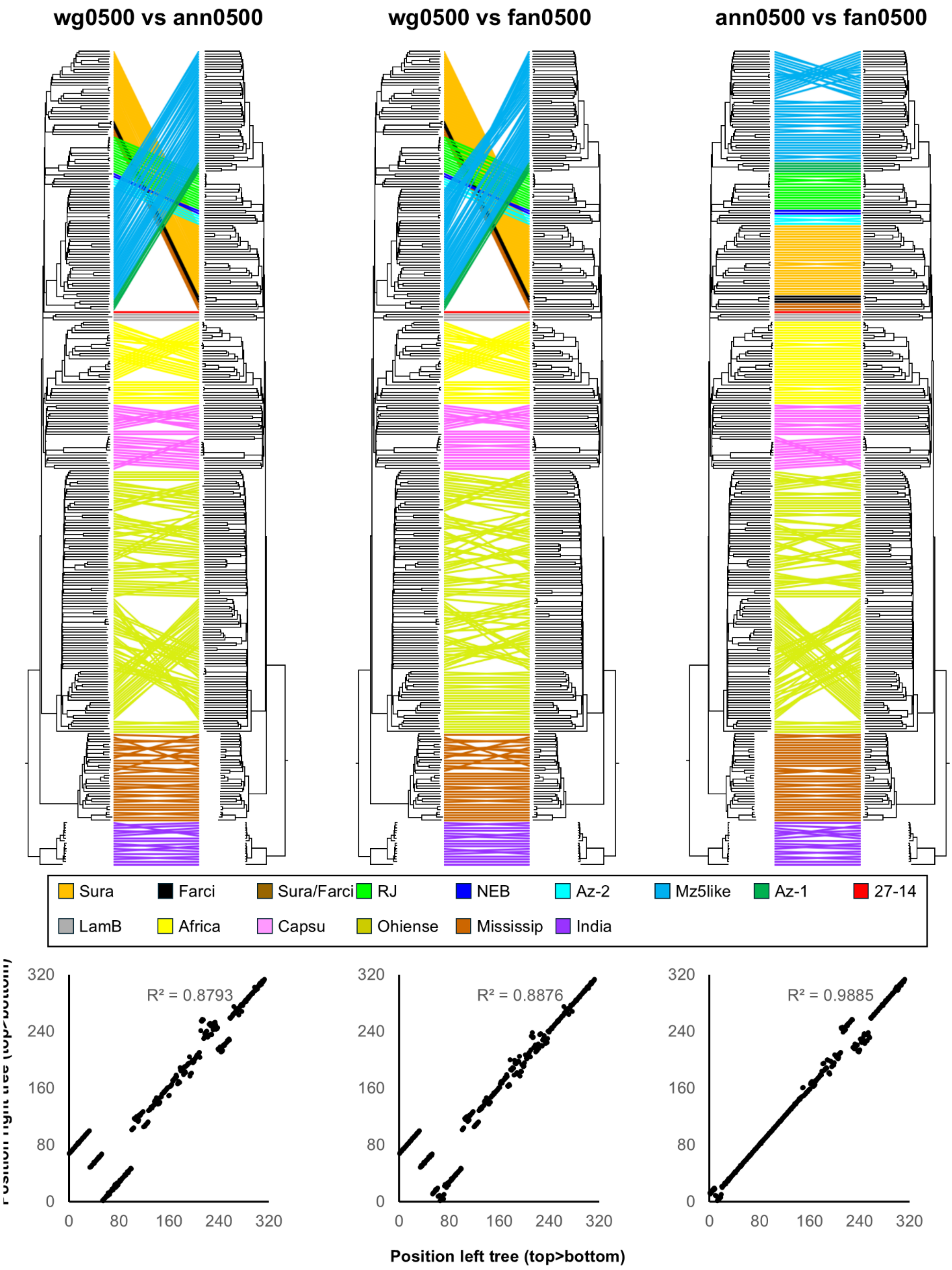


**Figure S7.** Comparison of genome-based MLST schemes for *Histoplasma capsulatum* with a minimum ORF length of 500 nt, using the complete 313 genome dataset from Figure 3. wg = based on whole genome sequences; ann = based on annotated features obtained from Genbank; fan = based on annotated features identified using Funannotate. The positioning in the tree is visualised using a tanglegram, and quantified using a scatter plot based on the positions in the respective trees from top to bottom.

**Table S1**. Overview of *Histoplasma capsulatum* lineages and previously proposed species names.

| Original lineage[1] | Proposed species[2] | Species name[3] | (sub)lineages[4] |
|---|---|---|---|
| NAm1 | *Histoplasma mississippiense* | *Histoplasma capsulatum* | Mississippiense |
| NAm2 | *Histoplasma ohiense* | *Histoplasma capsulatum* | Ohiense |
| Africa/duboisii | *Histoplasma duboisii* | *Histoplasma capsulatum var. duboisii* | Africa |
| LAmA | *Histoplasma suramericanum* | *Histoplasma capsulatum* | Suramericanum (Sura, Mz5-like, RJ, Amazon-1, Amazon-2, 27-14, NEB) |
| LAmB | *Histoplasma capsulatum* | *Histoplasma capsulatum* | LAmB |
| Panama/H81 | *Histoplasma capsulatum* sensu stricto | *Histoplasma capsulatum var. capsulatum* | Capsu |
| India | *Histoplasma capsulatum* | *Histoplasma capsulatum* | India |
| Farciminosum | *Histoplasma farciminosum* | *Histoplasma capsulatum var. farciminosum* | Farciminosum |

1. Lineages as described in (Kasuga *et al.*, 2003) and other pre-genomics studies.
2. Species proposed in (Sepulveda *et al.*, 2017; Sepulveda *et al.*, 2024) and other genomic studies.
3. Taxonomy as presented in NCBI Taxonomy (https://www.ncbi.nlm.nih.gov/Taxonomy) in August of 2026
4. Lineage and sublineage names as used in this study.

**Table S2.** Assembly metrics and BUSCO completeness scores of *Histoplasma capsulatum* samples/genome assemblies used in this study.

**Genomes used in main study**

| Order[1] | Assembly[2] | (Sub)Lineage | # contigs | Largest contig | Total length | N50 | L50 | #N's/100 kbp | BUSCO %[3] |
|---|---|---|---|---|---|---|---|---|---|
| 1 | Hcaps00164_SRR31893252 | Sura | 407 | 585936 | 31652328 | 155097 | 66 | 0 | 99.50% |
| 2 | Hcaps00166_SRR31893254 | Sura | 540 | 600693 | 31798684 | 117274 | 78 | 0 | 99.60% |
| 3 | Hcaps00170_SRR31893258 | Sura | 2206 | 223041 | 31130534 | 42851 | 218 | 0 | 99.40% |
| 4 | Hcaps00290_SRR8347492 | Sura | 625 | 383187 | 31263837 | 107945 | 91 | 0 | 99.60% |
| 5 | Hcaps00172_SRR31893260 | Sura | 637 | 338180 | 31269385 | 115753 | 86 | 0 | 99.60% |
| 6 | Hcaps00173_SRR31893261 | Sura | 457 | 443722 | 31066180 | 124782 | 79 | 0 | 99.50% |
| 7 | Hcaps00163_SRR31893251 | Sura | 472 | 476117 | 31042841 | 145660 | 72 | 0 | 99.50% |
| 8 | Hcaps00168_SRR31893256 | Sura | 672 | 487572 | 31890555 | 104389 | 85 | 0 | 99.50% |
| 9 | Hcaps00165_SRR31893253 | Sura | 332 | 779175 | 31359010 | 190007 | 49 | 0 | 99.60% |
| 10 | Hcaps00167_SRR31893255 | Sura | 569 | 614769 | 31538891 | 118496 | 84 | 0 | 99.50% |
| 11 | Hcaps00120_SRR25580058 | Sura | 533 | 630824 | 32354867 | 110331 | 83 | 0 | 99.40% |
| 12 | Hcaps00020_SRR21059728 | Sura | 326 | 538289 | 30206282 | 166585 | 57 | 0 | 99.40% |
| 13 | Hcaps00135_SRR25636145 | Sura | 1281 | 868799 | 32333008 | 191207 | 48 | 0 | 98.70% |
| 14 | Hcaps00014_SRR17307793 | Sura | 530 | 691253 | 31572530 | 153132 | 58 | 0 | 99.50% |
| 15 | Hcaps00177_SRR31893265 | Sura | 596 | 384914 | 33220457 | 106269 | 101 | 0 | 99.50% |
| 16 | Hcaps00176_SRR31893264 | Sura | 557 | 1111414 | 33280898 | 124814 | 82 | 0 | 99.50% |
| 17 | Hcaps00254_SRR6243645 | Sura | 364 | 653281 | 32882345 | 157726 | 68 | 0 | 99.50% |
| 18 | Hcaps00178_SRR31893266 | Sura | 406 | 506664 | 32861151 | 151099 | 69 | 0 | 99.40% |
| 19 | Hcaps00175_SRR31893263 | Sura | 937 | 349502 | 32747348 | 66717 | 144 | 0 | 99.10% |
| 20 | Hcaps00252_SRR6243643 | Sura | 643 | 427433 | 33541631 | 102361 | 92 | 0 | 99.50% |
| 21 | Hcaps00110_SRR25580048 | Sura | 884 | 271115 | 31638061 | 66030 | 157 | 0 | 99.30% |
| 22 | Hcaps00007_SRR17307786 | Sura | 533 | 558197 | 32938430 | 111446 | 96 | 0 | 99.50% |
| 23 | Hcaps00174_SRR31893262 | Sura | 692 | 409335 | 32749529 | 85373 | 119 | 0 | 99.40% |
| 24 | Hcaps00015_SRR17307794 | Sura | 823 | 336491 | 34043146 | 80031 | 129 | 0 | 99.30% |
| 25 | Hcaps00026_SRR21059734 | Sura | 530 | 568096 | 33622533 | 127441 | 85 | 0 | 99.50% |
| 26 | Hcaps00013_SRR17307792 | Sura | 893 | 484216 | 34621631 | 89900 | 102 | 0 | 99.40% |
| 27 | Hcaps00008_SRR17307787 | Sura | 589 | 497438 | 34055290 | 121465 | 90 | 0 | 99.50% |
| 28 | Hcaps00406_SRR29905426 | Farciminosum | 2112 | 291956 | 44615505 | 60459 | 208 | 0 | 99.30% |
| 29 | Hcaps00395_SRR29905415 | Farciminosum | 2263 | 306992 | 43704421 | 60660 | 178 | 0 | 99.30% |
| 30 | Hcaps00382_SRR29905402 | Farciminosum | 3655 | 368285 | 48851205 | 28412 | 499 | 0 | 98.00% |
| 31 | Hcaps00105_SRR25580043 | Sura/Farci | 1289 | 343577 | 46046595 | 68152 | 196 | 0 | 99.30% |

| | | | | | | | | | |
|---|---|---|---|---|---|---|---|---|---|
| 32 | Hcaps00001_ERR7414368 | Sura/Farci | 3749 | 212214 | 40263619 | 27805 | 366 | 0 | 99.20% |
| 33 | Hcapsulatum_GCA000313325 | Sura/Farci | 10513 | 90691 | 46124030 | 9657 | 1146 | 0 | 98.50% |
| 34 | Hcaps00275_SRR8084707 | RJ | 749 | 252243 | 30813866 | 72557 | 135 | 0 | 99.40% |
| 35 | Hcaps00276_SRR8084708 | RJ | 713 | 313428 | 30819194 | 78872 | 129 | 0 | 99.20% |
| 36 | Hcaps00277_SRR8084709 | RJ | 686 | 251413 | 30791266 | 79749 | 126 | 0 | 99.40% |
| 37 | Hcaps00273_SRR8084705 | RJ | 529 | 344899 | 30787728 | 104757 | 98 | 0 | 99.50% |
| 38 | Hcaps00274_SRR8084706 | RJ | 752 | 411043 | 30705243 | 90821 | 107 | 0 | 99.50% |
| 39 | Hcaps00272_SRR8084704 | RJ | 685 | 443275 | 30817848 | 75657 | 120 | 0 | 99.40% |
| 40 | Hcaps00282_SRR8084714 | RJ | 893 | 236513 | 30205054 | 67779 | 135 | 0 | 99.50% |
| 41 | Hcaps00284_SRR8084716 | RJ | 635 | 429591 | 30375770 | 81998 | 115 | 0 | 99.30% |
| 42 | Hcaps00285_SRR8084717 | RJ | 1132 | 339657 | 31162747 | 66241 | 144 | 0 | 99.50% |
| 43 | Hcaps00289_SRR8084721 | RJ | 464 | 371720 | 31460371 | 121627 | 84 | 0 | 99.50% |
| 44 | Hcaps00283_SRR8084715 | RJ | 1327 | 228488 | 26484994 | 45111 | 172 | 0 | 98.40% |
| 45 | Hcaps00288_SRR8084720 | RJ | 906 | 324698 | 30879075 | 66896 | 144 | 0 | 99.30% |
| 46 | Hcaps00287_SRR8084719 | RJ | 758 | 273347 | 30764699 | 79691 | 129 | 0 | 99.50% |
| 47 | Hcaps00278_SRR8084710 | RJ | 616 | 356313 | 31780432 | 98415 | 99 | 0 | 99.50% |
| 48 | Hcaps00193_SRR31893282 | Amazon-2 | 461 | 427687 | 31764482 | 125034 | 77 | 0 | 99.50% |
| 49 | Hcaps00192_SRR31893281 | Amazon-2 | 421 | 531639 | 30990018 | 127063 | 69 | 0 | 99.50% |
| 50 | Hcaps00189_SRR31893278 | Amazon-2 | 399 | 458557 | 29738317 | 137725 | 67 | 0 | 99.40% |
| 51 | Hcaps00191_SRR31893280 | Amazon-2 | 794 | 349648 | 30553049 | 74581 | 132 | 0 | 99.40% |
| 52 | Hcaps00280_SRR8084712 | NEB | 1032 | 396930 | 32170068 | 84521 | 117 | 0 | 99.40% |
| 53 | Hcaps00281_SRR8084713 | NEB | 483 | 561914 | 31935425 | 115820 | 80 | 0 | 99.60% |
| 54 | Hcaps00210_SRR31893300 | Mz5-like | 707 | 445184 | 31988547 | 88331 | 107 | 0 | 99.40% |
| 55 | Hcaps00196_SRR31893286 | Mz5-like | 1017 | 252287 | 32628527 | 58959 | 178 | 0 | 99.30% |
| 56 | Hcaps00180_SRR31893268 | Mz5-like | 1122 | 231392 | 32666681 | 56580 | 188 | 0 | 99.30% |
| 57 | Hcaps00199_SRR31893289 | Mz5-like | 1183 | 232485 | 32585368 | 56388 | 182 | 0 | 99.30% |
| 58 | Hcaps00169_SRR31893257 | Mz5-like | 1012 | 248656 | 32853988 | 62589 | 164 | 0 | 99.30% |
| 59 | Hcaps00203_SRR31893293 | Mz5-like | 1015 | 264988 | 32855917 | 61248 | 166 | 0 | 99.20% |
| 60 | Hcaps00198_SRR31893288 | Mz5-like | 1157 | 214472 | 32513731 | 53766 | 184 | 0 | 99.40% |
| 61 | Hcaps00201_SRR31893291 | Mz5-like | 1260 | 198599 | 32358877 | 49637 | 208 | 0 | 98.50% |
| 62 | Hcaps00222_SRR31893312 | Mz5-like | 885 | 312148 | 32038552 | 68907 | 154 | 0 | 99.40% |
| 63 | Hcaps00212_SRR31893302 | Mz5-like | 2034 | 219476 | 32412142 | 29168 | 323 | 0 | 98.30% |
| 64 | Hcaps00218_SRR31893308 | Mz5-like | 764 | 408991 | 31851069 | 73227 | 132 | 0 | 99.40% |
| 65 | Hcaps00208_SRR31893298 | Mz5-like | 1026 | 283231 | 32721045 | 60168 | 165 | 0 | 99.40% |
| 66 | Hcaps00217_SRR31893307 | Mz5-like | 1042 | 444191 | 32374800 | 65123 | 153 | 0 | 99.40% |

| | | | | | | | | | |
|---|---|---|---|---|---|---|---|---|---|
| 67 | Hcaps00223_SRR31893313 | Mz5-like | 656 | 384745 | 32169386 | 99381 | 95 | 0 | 99.40% |
| 68 | Hcaps00204_SRR31893294 | Mz5-like | 1051 | 300544 | 32315074 | 63203 | 157 | 0 | 99.40% |
| 69 | Hcaps00161_SRR31893249 | Mz5-like | 947 | 253881 | 32575390 | 69359 | 143 | 0 | 99.40% |
| 70 | Hcaps00207_SRR31893297 | Mz5-like | 1115 | 294379 | 32922702 | 54022 | 184 | 0 | 99.30% |
| 71 | Hcaps00200_SRR31893290 | Mz5-like | 966 | 320073 | 32929830 | 64556 | 157 | 0 | 99.30% |
| 72 | Hcaps00190_SRR31893279 | Mz5-like | 2071 | 192558 | 33238130 | 41300 | 247 | 0 | 99.40% |
| 73 | Hcaps00219_SRR31893309 | Mz5-like | 1497 | 307979 | 35399589 | 75733 | 139 | 0 | 99.50% |
| 74 | Hcaps00211_SRR31893301 | Mz5-like | 1506 | 396432 | 35701311 | 67592 | 144 | 0 | 99.30% |
| 75 | Hcaps00221_SRR31893311 | Mz5-like | 983 | 398473 | 33059098 | 65363 | 157 | 0 | 99.50% |
| 76 | Hcaps00215_SRR31893305 | Mz5-like | 1057 | 246852 | 32443599 | 55196 | 172 | 0 | 99.30% |
| 77 | Hcaps00213_SRR31893303 | Mz5-like | 604 | 354504 | 32558717 | 101776 | 105 | 0 | 99.50% |
| 78 | Hcaps00202_SRR31893292 | Mz5-like | 780 | 303857 | 32565667 | 70956 | 141 | 0 | 99.30% |
| 79 | Hcaps00209_SRR31893299 | Mz5-like | 656 | 304246 | 32507497 | 93674 | 110 | 0 | 99.40% |
| 80 | Hcaps00160_SRR31893248 | Mz5-like | 905 | 256168 | 32094156 | 61989 | 156 | 0 | 99.40% |
| 81 | Hcaps00206_SRR31893296 | Mz5-like | 1050 | 336082 | 32792249 | 59276 | 177 | 0 | 99.40% |
| 82 | Hcaps00214_SRR31893304 | Mz5-like | 691 | 373706 | 32960842 | 85194 | 117 | 0 | 99.50% |
| 83 | Hcaps00159_SRR31893247 | Mz5-like | 633 | 460634 | 31633303 | 93534 | 104 | 0 | 99.40% |
| 84 | Hcaps00228_SRR32121389 | Mz5-like | 547 | 637495 | 32928890 | 130429 | 76 | 0 | 99.50% |
| 85 | Hcaps00227_SRR32121388 | Mz5-like | 549 | 500286 | 33807233 | 134657 | 77 | 0 | 99.60% |
| 86 | Hcaps00241_SRR32121402 | Mz5-like | 657 | 542027 | 37612689 | 132648 | 91 | 0 | 99.60% |
| 87 | Hcaps00240_SRR32121401 | Mz5-like | 789 | 571122 | 37612534 | 106759 | 107 | 0 | 99.50% |
| 88 | Hcaps00226_SRR32121387 | Mz5-like | 584 | 463711 | 33197191 | 124379 | 77 | 0 | 99.40% |
| 89 | Hcaps00230_SRR32121391 | Mz5-like | 650 | 559834 | 33616831 | 114291 | 86 | 0 | 99.50% |
| 90 | Hcaps00220_SRR31893310 | Mz5-like | 1023 | 307049 | 32472188 | 61075 | 155 | 0 | 99.30% |
| 91 | Hcaps00231_SRR32121392 | Mz5-like | 593 | 730580 | 32488301 | 120934 | 77 | 0 | 99.50% |
| 92 | Hcaps00205_SRR31893295 | Mz5-like | 3075 | 252010 | 40098870 | 44676 | 256 | 0 | 99.30% |
| 93 | Hcaps00229_SRR32121390 | Mz5-like | 611 | 584573 | 34057862 | 120396 | 83 | 0 | 99.50% |
| 94 | Hcaps00258_SRR6243649 | Mz5-like | 1099 | 722301 | 33172518 | 116407 | 83 | 0 | 99.40% |
| 95 | Hcaps00374_SRR33700783 | Mz5-like | 875 | 323987 | 31534574 | 70519 | 146 | 0 | 99.30% |
| 96 | Hcaps00042_SRR21059750 | Mz5-like | 623 | 473678 | 32885925 | 111576 | 92 | 0 | 99.50% |
| 97 | Hcaps00195_SRR31893285 | Amazon-1 | 1237 | 280865 | 33182976 | 63630 | 165 | 0 | 99.40% |
| 98 | Hcaps00194_SRR31893284 | Amazon-1 | 1461 | 234120 | 33184251 | 46414 | 206 | 0 | 99.30% |
| 99 | Hcaps00294_SRR31893283 | Amazon-1 | 1629 | 240399 | 32611780 | 45730 | 219 | 0 | 99.20% |
| 100 | Hcaps00197_SRR31893287 | Amazon-1 | 1274 | 250067 | 32311184 | 57050 | 172 | 0 | 99.50% |
| 101 | Hcaps00255_SRR6243646 | 27-14 | 602 | 472656 | 32305380 | 111687 | 88 | 0 | 99.50% |

| | | | | | | | | | |
|---|---|---|---|---|---|---|---|---|---|
| 102 | Hcaps00279_SRR8084711 | LAmB | 549 | 289121 | 29977273 | 102415 | 95 | 0 | 99.50% |
| 103 | Hcaps00162_SRR31893250 | LAmB | 1381 | 406976 | 31021971 | 73667 | 127 | 0 | 99.50% |
| 104 | Hcaps00286_SRR8084718 | LAmB | 1511 | 286387 | 31630153 | 74323 | 127 | 0 | 99.40% |
| 105 | Hcaps00400_SRR29905420 | Africa | 1291 | 410894 | 41530322 | 70657 | 188 | 0 | 99.50% |
| 106 | Hcaps00402_SRR29905422 | Africa | 1145 | 393534 | 42088653 | 84182 | 145 | 0 | 99.70% |
| 107 | Hcaps00391_SRR29905411 | Africa | 856 | 693978 | 38988175 | 152197 | 74 | 0 | 99.70% |
| 108 | Hcaps00385_SRR29905405 | Africa | 899 | 616345 | 41136836 | 122900 | 91 | 0 | 99.70% |
| 109 | Hcaps00392_SRR29905412 | Africa | 803 | 531084 | 38311988 | 147522 | 73 | 0 | 99.70% |
| 110 | Hcaps00253_SRR6243644 | Africa | 906 | 448019 | 36802196 | 110447 | 99 | 0 | 99.70% |
| 111 | Hcapsulatum_GCA017310615 | Africa | 7 | 9359328 | 37996987 | 7000705 | 3 | 0 | 99.70% |
| 112 | Hcaps00248_SRR6243639 | Africa | 976 | 504603 | 36823087 | 93415 | 109 | 0 | 99.60% |
| 113 | Hcaps00396_SRR29905416 | Africa | 1026 | 298112 | 38448895 | 80409 | 150 | 0 | 99.60% |
| 114 | Hcaps00393_SRR29905413 | Africa | 701 | 1609975 | 37830355 | 198922 | 52 | 0 | 99.60% |
| 115 | Hcaps00375_SRR29905395 | Africa | 721 | 521466 | 37865214 | 124281 | 90 | 0 | 99.60% |
| 116 | Hcaps00379_SRR29905399 | Africa | 755 | 752746 | 37471985 | 183361 | 61 | 0 | 99.70% |
| 117 | Hcaps00387_SRR29905407 | Africa | 715 | 831256 | 37803783 | 191079 | 59 | 0 | 99.70% |
| 118 | Hcaps00390_SRR29905410 | Africa | 587 | 1575874 | 36980428 | 217898 | 44 | 0 | 99.70% |
| 119 | Hcaps00397_SRR29905417 | Africa | 1735 | 230497 | 29934054 | 59370 | 150 | 0 | 99.40% |
| 120 | Hcaps00404_SRR29905424 | Africa | 806 | 430309 | 38369572 | 117134 | 98 | 0 | 99.50% |
| 121 | Hcaps00405_SRR29905425 | Africa | 736 | 758420 | 38010844 | 222522 | 57 | 0 | 99.60% |
| 122 | Hcaps00381_SRR29905401 | Africa | 784 | 430135 | 40695162 | 117228 | 107 | 0 | 99.50% |
| 123 | Hcaps00384_SRR29905404 | Africa | 937 | 430110 | 38440091 | 110369 | 111 | 0 | 99.50% |
| 124 | Hcaps00377_SRR29905397 | Africa | 1029 | 564976 | 37184636 | 102212 | 111 | 0 | 99.50% |
| 125 | Hcaps00378_SRR29905398 | Africa | 599 | 905115 | 37938293 | 145843 | 78 | 0 | 99.60% |
| 126 | Hcaps00398_SRR29905418 | Africa | 1106 | 439505 | 37672297 | 72187 | 152 | 0 | 99.50% |
| 127 | Hcapsulatum_GCA000151035 | Africa | 49 | 6121860 | 38961716 | 1740271 | 7 | 7155.11 | 94.20% |
| 128 | Hcaps00389_SRR29905409 | Africa | 1277 | 741652 | 43039917 | 134004 | 85 | 0 | 99.60% |
| 129 | Hcaps00383_SRR29905403 | Africa | 7434 | 86959 | 37829933 | 8658 | 1202 | 0 | 69.00% |
| 130 | Hcaps00403_SRR29905423 | Africa | 963 | 581505 | 40184414 | 117858 | 99 | 0 | 99.60% |
| 131 | Hcaps00376_SRR29905396 | Africa | 816 | 1588833 | 40069503 | 180386 | 59 | 0 | 99.70% |
| 132 | Hcaps00394_SRR29905414 | Africa | 1448 | 320215 | 42880552 | 69668 | 187 | 0 | 99.30% |
| 133 | Hcaps00399_SRR29905419 | Africa | 1205 | 359916 | 38817973 | 72848 | 161 | 0 | 99.50% |
| 134 | Hcaps00401_SRR29905421 | Africa | 1809 | 498519 | 43889345 | 68027 | 170 | 0 | 99.50% |
| 135 | Hcaps00380_SRR29905400 | Africa | 1065 | 1452042 | 38955321 | 395131 | 31 | 0 | 99.60% |
| 136 | Hcaps00386_SRR29905406 | Africa | 845 | 1015043 | 38499857 | 163104 | 62 | 0 | 99.40% |

| | | | | | | | | | |
|---|---|---|---|---|---|---|---|---|---|
| 137 | Hcaps00182_SRR31893271 | Capsu | 344 | 693951 | 29692122 | 161762 | 59 | 0 | 99.50% |
| 138 | Hcaps00183_SRR31893272 | Capsu | 606 | 325117 | 31085791 | 109174 | 93 | 0 | 99.50% |
| 139 | Hcaps00185_SRR31893274 | Capsu | 643 | 345760 | 30175063 | 80676 | 115 | 0 | 99.50% |
| 140 | Hcaps00179_SRR31893267 | Capsu | 455 | 437319 | 31228880 | 142484 | 68 | 0 | 99.60% |
| 141 | Hcaps00232_SRR32121393 | Capsu | 680 | 398208 | 30360524 | 83137 | 114 | 0 | 99.50% |
| 142 | Hcaps00181_SRR31893270 | Capsu | 320 | 588461 | 30302368 | 174157 | 54 | 0 | 99.50% |
| 143 | Hcaps00186_SRR31893275 | Capsu | 573 | 340074 | 30264186 | 99823 | 99 | 0 | 99.50% |
| 144 | Hcaps00187_SRR31893276 | Capsu | 582 | 435316 | 30137361 | 96090 | 93 | 0 | 99.60% |
| 145 | Hcaps00188_SRR31893277 | Capsu | 598 | 410962 | 30303367 | 94887 | 100 | 0 | 99.50% |
| 146 | Hcaps00293_SRR31893269 | Capsu | 1130 | 306557 | 29778142 | 80087 | 116 | 0 | 99.60% |
| 147 | Hcaps00184_SRR31893273 | Capsu | 426 | 771420 | 31858067 | 135770 | 70 | 0 | 99.60% |
| 148 | Hcaps00261_SRR6243652 | Capsu | 215 | 784907 | 32067491 | 307800 | 37 | 0 | 99.60% |
| 149 | Hcaps00259_SRR6243650 | Capsu | 511 | 512959 | 30400550 | 120619 | 80 | 0 | 99.60% |
| 150 | Hcaps00256_SRR6243647 | Capsu | 656 | 331092 | 30387254 | 90328 | 107 | 0 | 99.40% |
| 151 | Hcapsulatum_GCA017607465 | Capsu | 12 | 9773074 | 30991520 | 4063218 | 3 | 0 | 99.70% |
| 152 | Hcaps00235_SRR32121396 | Capsu | 740 | 360794 | 30431522 | 77929 | 117 | 0 | 99.40% |
| 153 | Hcaps00234_SRR32121395 | Capsu | 443 | 458127 | 30453995 | 125847 | 75 | 0 | 99.50% |
| 154 | Hcapsulatum_GCA017355575 | Capsu | 7 | 7332157 | 31111494 | 5585295 | 3 | 0 | 99.70% |
| 155 | Hcaps00233_SRR32121394 | Capsu | 2495 | 113322 | 27859423 | 17094 | 502 | 0 | 84.30% |
| 156 | Hcaps00237_SRR32121398 | Capsu | 555 | 638703 | 32116608 | 141745 | 70 | 0 | 99.60% |
| 157 | Hcaps00236_SRR32121397 | Capsu | 342 | 732971 | 30140011 | 165476 | 56 | 0 | 99.70% |
| 158 | Hcaps00238_SRR32121399 | Capsu | 292 | 608210 | 29358000 | 202867 | 46 | 0 | 99.60% |
| 159 | Hcaps00225_SRR32121386 | Capsu | 3440 | 747284 | 34736327 | 16724 | 516 | 0 | 96.30% |
| 160 | Hcaps00224_SRR32121385 | Capsu | 456 | 436825 | 31432304 | 131016 | 72 | 0 | 99.60% |
| 161 | Hcaps00239_SRR32121400 | Capsu | 407 | 566064 | 32000535 | 150700 | 67 | 0 | 99.40% |
| 162 | Hcaps00046_SRR21059754 | Ohiense | 1251 | 308952 | 37913006 | 67939 | 171 | 0 | 99.20% |
| 163 | Hcaps00057_SRR21059765 | Ohiense | 1327 | 265916 | 38396880 | 63202 | 186 | 0 | 99.20% |
| 164 | Hcaps00056_SRR21059764 | Ohiense | 1359 | 239932 | 38566544 | 59456 | 199 | 0 | 99.10% |
| 165 | Hcaps00062_SRR21059770 | Ohiense | 1240 | 256136 | 38933421 | 71619 | 166 | 0 | 99.20% |
| 166 | Hcaps00054_SRR21059762 | Ohiense | 1366 | 242890 | 38448551 | 61539 | 196 | 0 | 99.20% |
| 167 | Hcaps00040_SRR21059748 | Ohiense | 1140 | 329683 | 38480064 | 68963 | 172 | 0 | 99.20% |
| 168 | Hcaps00055_SRR21059763 | Ohiense | 1318 | 282162 | 38637043 | 66431 | 179 | 0 | 99.20% |
| 169 | Hcaps00031_SRR21059739 | Ohiense | 1299 | 293849 | 38532582 | 59112 | 196 | 0 | 99.30% |
| 170 | Hcaps00407_SRR29905427 | Ohiense | 1596 | 190085 | 38687518 | 49490 | 231 | 0 | 99.30% |
| 171 | Hcaps00019_SRR21059727 | Ohiense | 1144 | 276509 | 38638059 | 68761 | 174 | 0 | 99.10% |

| | | | | | | | | | |
|---|---|---|---|---|---|---|---|---|---|
| 172 | Hcaps00018_SRR21059726 | Ohiense | 1258 | 403121 | 38763638 | 63589 | 191 | 0 | 99.30% |
| 173 | Hcaps00045_SRR21059753 | Ohiense | 1311 | 365612 | 39169358 | 65231 | 188 | 0 | 99.30% |
| 174 | Hcaps00063_SRR21059771 | Ohiense | 1942 | 387783 | 40447048 | 64397 | 180 | 0 | 99.30% |
| 175 | Hcaps00048_SRR21059756 | Ohiense | 1298 | 512898 | 39027790 | 70756 | 165 | 0 | 99.30% |
| 176 | Hcaps00059_SRR21059767 | Ohiense | 1278 | 269616 | 38832297 | 65975 | 185 | 0 | 99.10% |
| 177 | Hcaps00052_SRR21059760 | Ohiense | 7090 | 252050 | 42804321 | 31573 | 330 | 0 | 94.20% |
| 178 | Hcaps00021_SRR21059729 | Ohiense | 1321 | 309421 | 39089267 | 68041 | 181 | 0 | 99.30% |
| 179 | Hcaps00053_SRR21059761 | Ohiense | 1181 | 413122 | 38572333 | 70328 | 163 | 0 | 99.30% |
| 180 | Hcaps00049_SRR21059757 | Ohiense | 1223 | 375355 | 38384130 | 71475 | 164 | 0 | 99.30% |
| 181 | Hcaps00111_SRR25580049 | Ohiense | 1892 | 231272 | 38562521 | 42719 | 273 | 0 | 99.10% |
| 182 | Hcaps00093_SRR25580031 | Ohiense | 2719 | 161300 | 39125736 | 29624 | 412 | 0 | 98.80% |
| 183 | Hcaps00043_SRR21059751 | Ohiense | 1504 | 227026 | 38511922 | 56459 | 206 | 0 | 99.10% |
| 184 | Hcaps00047_SRR21059755 | Ohiense | 1186 | 381541 | 38323036 | 72078 | 166 | 0 | 99.10% |
| 185 | Hcaps00039_SRR21059747 | Ohiense | 1118 | 423232 | 38595693 | 75800 | 157 | 0 | 99.20% |
| 186 | Hcaps00061_SRR21059769 | Ohiense | 1197 | 305500 | 38997137 | 72376 | 167 | 0 | 99.20% |
| 187 | Hcaps00123_SRR25580061 | Ohiense | 2406 | 164084 | 38534281 | 31188 | 371 | 0 | 99.00% |
| 188 | Hcaps00132_SRR25580070 | Ohiense | 2696 | 151179 | 38593581 | 29092 | 402 | 0 | 98.80% |
| 189 | Hcaps00099_SRR25580037 | Ohiense | 2300 | 152590 | 38141801 | 34585 | 345 | 0 | 99.00% |
| 190 | Hcaps00072_SRR25580010 | Ohiense | 4023 | 96605 | 37321098 | 17487 | 618 | 0 | 98.20% |
| 191 | Hcaps00065_SRR21059773 | Ohiense | 1092 | 287670 | 37424684 | 81287 | 146 | 0 | 99.30% |
| 192 | Hcaps00106_SRR25580044 | Ohiense | 2269 | 161653 | 37861795 | 32560 | 354 | 0 | 98.70% |
| 193 | Hcaps00121_SRR25580059 | Ohiense | 2026 | 198231 | 38329711 | 37635 | 311 | 0 | 99.10% |
| 194 | Hcaps00094_SRR25580032 | Ohiense | 2729 | 231875 | 38735100 | 29122 | 412 | 0 | 98.80% |
| 195 | Hcaps00075_SRR25580013 | Ohiense | 2313 | 150138 | 37952969 | 31597 | 364 | 0 | 99.00% |
| 196 | Hcaps00116_SRR25580054 | Ohiense | 2425 | 211825 | 38504842 | 31573 | 389 | 0 | 98.80% |
| 197 | Hcaps00108_SRR25580046 | Ohiense | 1905 | 206370 | 38266003 | 45828 | 251 | 0 | 99.00% |
| 198 | Hcaps00115_SRR25580053 | Ohiense | 1816 | 192678 | 38199155 | 40451 | 296 | 0 | 99.10% |
| 199 | Hcaps00118_SRR25580056 | Ohiense | 1565 | 195775 | 37891040 | 45509 | 255 | 0 | 99.20% |
| 200 | Hcaps00107_SRR25580045 | Ohiense | 6363 | 136456 | 46923311 | 14704 | 856 | 0 | 97.50% |
| 201 | Hcaps00022_SRR21059730 | Ohiense | 1415 | 384060 | 38855437 | 61350 | 185 | 0 | 99.30% |
| 202 | Hcaps00077_SRR25580015 | Ohiense | 2151 | 187081 | 38622218 | 36324 | 322 | 0 | 99.00% |
| 203 | Hcaps00088_SRR25580026 | Ohiense | 3084 | 138227 | 38224293 | 24674 | 476 | 0 | 98.60% |
| 204 | Hcaps00050_SRR21059758 | Ohiense | 1170 | 305135 | 38665158 | 73099 | 159 | 0 | 99.30% |
| 205 | Hcaps00064_SRR21059772 | Ohiense | 1084 | 240896 | 37596587 | 75122 | 161 | 0 | 99.30% |
| 206 | Hcaps00098_SRR25580036 | Ohiense | 2106 | 230823 | 37965754 | 36493 | 328 | 0 | 99.10% |

| | | | | | | | | | |
|---|---|---|---|---|---|---|---|---|---|
| 207 | Hcaps00267_SRR6243658 | Ohiense | 2555 | 172987 | 36484667 | 35190 | 272 | 0 | 99.10% |
| 208 | Hcaps00114_SRR25580052 | Ohiense | 2901 | 178178 | 39891622 | 31439 | 384 | 0 | 97.90% |
| 209 | Hcaps00095_SRR25580033 | Ohiense | 3030 | 115500 | 38850374 | 26113 | 456 | 0 | 98.80% |
| 210 | Hcaps00129_SRR25580067 | Ohiense | 2565 | 171491 | 37966845 | 30326 | 389 | 0 | 98.80% |
| 211 | Hcaps00101_SRR25580039 | Ohiense | 2480 | 138009 | 38331740 | 32089 | 370 | 0 | 98.80% |
| 212 | Hcaps00262_SRR6243653 | Ohiense | 3115 | 155546 | 36686116 | 27655 | 366 | 0 | 99.00% |
| 213 | Hcaps00265_SRR6243656 | Ohiense | 1975 | 183469 | 36496427 | 41654 | 249 | 0 | 99.30% |
| 214 | Hcaps00266_SRR6243657 | Ohiense | 2187 | 218769 | 36542638 | 36524 | 280 | 0 | 99.30% |
| 215 | Hcaps00246_SRR6243637 | Ohiense | 2733 | 179805 | 36671030 | 28281 | 350 | 0 | 99.00% |
| 216 | Hcaps00264_SRR6243655 | Ohiense | 2122 | 161196 | 36840804 | 38551 | 283 | 0 | 99.10% |
| 217 | Hcaps00263_SRR6243654 | Ohiense | 2555 | 233514 | 36363773 | 36593 | 270 | 0 | 99.20% |
| 218 | Hcaps00127_SRR25580065 | Ohiense | 2502 | 163305 | 37561955 | 30233 | 377 | 0 | 98.80% |
| 219 | Hcaps00260_SRR6243651 | Ohiense | 1111 | 477469 | 38270042 | 105113 | 102 | 0 | 99.30% |
| 220 | Hcapsulatum_GCA017607445 | Ohiense | 12 | 8217122 | 39447273 | 6680169 | 3 | 0 | 99.20% |
| 221 | Hcaps00269_SRR6243660 | Ohiense | 2167 | 162300 | 37216251 | 35593 | 297 | 0 | 99.00% |
| 222 | Hcaps00130_SRR25580068 | Ohiense | 2084 | 180808 | 38081611 | 35147 | 336 | 0 | 99.10% |
| 223 | Hcaps00128_SRR25580066 | Ohiense | 1813 | 204694 | 37765110 | 41288 | 283 | 0 | 99.00% |
| 224 | Hcaps00117_SRR25580055 | Ohiense | 2330 | 154752 | 37984907 | 32387 | 356 | 0 | 98.80% |
| 225 | Hcaps00103_SRR25580041 | Ohiense | 2030 | 176068 | 37625123 | 37890 | 311 | 0 | 99.00% |
| 226 | Hcaps00060_SRR21059768 | Ohiense | 1096 | 398101 | 38706337 | 76884 | 156 | 0 | 99.20% |
| 227 | Hcaps00036_SRR21059744 | Ohiense | 1404 | 293572 | 38555504 | 61171 | 187 | 0 | 99.20% |
| 228 | Hcaps00133_SRR25580071 | Ohiense | 2692 | 141433 | 38387587 | 28493 | 410 | 0 | 98.90% |
| 229 | Hcaps00100_SRR25580038 | Ohiense | 2167 | 148501 | 38417053 | 36123 | 336 | 0 | 99.00% |
| 230 | Hcaps00268_SRR6243659 | Ohiense | 2123 | 385961 | 36043909 | 43346 | 220 | 0 | 99.30% |
| 231 | Hcaps00038_SRR21059746 | Ohiense | 1129 | 311804 | 37712707 | 75505 | 157 | 0 | 99.20% |
| 232 | Hcaps00037_SRR21059745 | Ohiense | 1299 | 329351 | 39077843 | 64102 | 190 | 0 | 99.20% |
| 233 | Hcaps00051_SRR21059759 | Ohiense | 1156 | 285877 | 38235966 | 71443 | 161 | 0 | 99.20% |
| 234 | Hcaps00035_SRR21059743 | Ohiense | 1274 | 318470 | 38363982 | 65588 | 179 | 0 | 99.30% |
| 235 | Hcaps00096_SRR25580034 | Ohiense | 1653 | 238454 | 37771963 | 46870 | 242 | 0 | 99.10% |
| 236 | Hcaps00122_SRR25580060 | Ohiense | 1923 | 197506 | 38574554 | 39723 | 292 | 0 | 99.10% |
| 237 | Hcaps00080_SRR25580018 | Ohiense | 1543 | 271743 | 38846681 | 55922 | 215 | 0 | 99.10% |
| 238 | Hcaps00131_SRR25580069 | Ohiense | 2417 | 157834 | 38493454 | 32247 | 371 | 0 | 99.00% |
| 239 | Hcaps00024_SRR21059732 | Ohiense | 1215 | 293341 | 39184339 | 64317 | 180 | 0 | 99.10% |
| 240 | Hcaps00058_SRR21059766 | Ohiense | 1330 | 309242 | 38951156 | 63123 | 182 | 0 | 99.20% |
| 241 | Hcaps00034_SRR21059742 | Ohiense | 1536 | 233434 | 38842535 | 52253 | 219 | 0 | 99.10% |

| | | | | | | | | | |
|---|---|---|---|---|---|---|---|---|---|
| 242 | Hcaps00027_SRR21059735 | Ohiense | 1441 | 250620 | 39111310 | 57590 | 203 | 0 | 99.20% |
| 243 | Hcaps00025_SRR21059733 | Ohiense | 1343 | 279028 | 39215373 | 65000 | 182 | 0 | 99.20% |
| 244 | Hcaps00030_SRR21059738 | Ohiense | 1385 | 396446 | 39320565 | 57058 | 199 | 0 | 99.20% |
| 245 | Hcaps00032_SRR21059740 | Ohiense | 1622 | 333185 | 39812597 | 56185 | 208 | 0 | 99.20% |
| 246 | Hcaps00033_SRR21059741 | Ohiense | 1486 | 294129 | 39654532 | 56369 | 216 | 0 | 99.20% |
| 247 | Hcaps00028_SRR21059736 | Ohiense | 1412 | 252522 | 38835008 | 58508 | 205 | 0 | 99.20% |
| 248 | Hcaps00029_SRR21059737 | Ohiense | 1437 | 252461 | 40553390 | 61777 | 195 | 0 | 99.30% |
| 249 | Hcaps00023_SRR21059731 | Ohiense | 1279 | 259393 | 39171075 | 62186 | 193 | 0 | 99.20% |
| 250 | Hcaps00104_SRR25580042 | Ohiense | 1946 | 190369 | 39628125 | 39350 | 305 | 0 | 99.10% |
| 251 | Hcaps00011_SRR17307790 | Ohiense | 2458 | 208018 | 41594943 | 39309 | 315 | 0 | 99.10% |
| 252 | Hcaps00017_SRR17307796 | Ohiense | 2265 | 184783 | 42164464 | 38360 | 327 | 0 | 98.90% |
| 253 | Hcaps00016_SRR17307795 | Ohiense | 2258 | 190133 | 41010648 | 38513 | 318 | 0 | 99.10% |
| 254 | Hcaps00009_SRR17307788 | Ohiense | 2196 | 187985 | 42202707 | 39812 | 315 | 0 | 99.00% |
| 255 | Hcaps00012_SRR17307791 | Ohiense | 2351 | 255231 | 41015435 | 36694 | 324 | 0 | 99.20% |
| 256 | Hcaps00097_SRR25580035 | Ohiense | 2733 | 159943 | 40352778 | 31191 | 388 | 0 | 99.10% |
| 257 | Hcaps00124_SRR25580062 | Ohiense | 2196 | 200075 | 38873639 | 36800 | 321 | 0 | 99.10% |
| 258 | Hcaps00073_SRR25580011 | Ohiense | 3100 | 130972 | 39600386 | 27121 | 440 | 0 | 98.80% |
| 259 | Hcaps00109_SRR25580047 | Ohiense | 2515 | 155020 | 40074822 | 31428 | 385 | 0 | 99.00% |
| 260 | Hcaps00074_SRR25580012 | Ohiense | 2851 | 123739 | 39413052 | 27659 | 439 | 0 | 98.70% |
| 261 | Hcaps00044_SRR21059752 | Ohiense | 1759 | 235028 | 40165586 | 50109 | 241 | 0 | 99.10% |
| 262 | Hcaps00113_SRR25580051 | Ohiense | 17477 | 67565 | 49194258 | 4797 | 2592 | 0 | 81.50% |
| 263 | Hcaps00084_SRR25580022 | Mississippiense | 1502 | 257657 | 30023469 | 51289 | 177 | 0 | 98.90% |
| 264 | Hcaps00076_SRR25580014 | Mississippiense | 1839 | 167244 | 30056672 | 36990 | 242 | 0 | 98.70% |
| 265 | Hcaps00089_SRR25580027 | Mississippiense | 2801 | 136773 | 29978290 | 21618 | 400 | 0 | 98.40% |
| 266 | Hcaps00081_SRR25580019 | Mississippiense | 1982 | 191409 | 30111338 | 33639 | 257 | 0 | 98.70% |
| 267 | Hcaps00087_SRR25580025 | Mississippiense | 2811 | 126137 | 29932142 | 22265 | 399 | 0 | 98.30% |
| 268 | Hcaps00082_SRR25580020 | Mississippiense | 1902 | 195441 | 30089235 | 35002 | 252 | 0 | 98.80% |
| 269 | Hcaps00125_SRR25580063 | Mississippiense | 2593 | 109625 | 29971992 | 24067 | 373 | 0 | 98.40% |
| 270 | Hcaps00091_SRR25580029 | Mississippiense | 2527 | 109848 | 29908512 | 24868 | 359 | 0 | 98.50% |
| 271 | Hcaps00092_SRR25580030 | Mississippiense | 3359 | 110051 | 29722253 | 18146 | 488 | 0 | 98.10% |
| 272 | Hcaps00085_SRR25580023 | Mississippiense | 2624 | 147402 | 30005754 | 22683 | 385 | 0 | 98.50% |
| 273 | Hcaps00079_SRR25580017 | Mississippiense | 1774 | 263105 | 30097091 | 40477 | 230 | 0 | 98.70% |
| 274 | Hcaps00078_SRR25580016 | Mississippiense | 1816 | 174928 | 30061422 | 38473 | 243 | 0 | 98.70% |
| 275 | Hcaps00102_SRR25580040 | Mississippiense | 2699 | 116467 | 29957935 | 23026 | 385 | 0 | 98.40% |
| 276 | Hcaps00136_SRR25758805 | Mississippiense | 2556 | 151861 | 29920567 | 28525 | 316 | 0 | 98.10% |

| | | | | | | | | | |
|---|---|---|---|---|---|---|---|---|---|
| 277 | Hcaps00090_SRR25580028 | Mississippiense | 3367 | 96182 | 29722943 | 17376 | 517 | 0 | 98.20% |
| 278 | Hcaps00126_SRR25580064 | Mississippiense | 3106 | 81180 | 29854174 | 19721 | 456 | 0 | 98.30% |
| 279 | Hcaps00251_SRR6243642 | Mississippiense | 1704 | 211785 | 29367023 | 41380 | 216 | 0 | 98.80% |
| 280 | Hcaps00134_SRR25636130 | Mississippiense | 2107 | 180462 | 29931170 | 37732 | 229 | 0 | 98.70% |
| 281 | Hcaps00112_SRR25580050 | Mississippiense | 2692 | 109785 | 29746702 | 23451 | 378 | 0 | 98.60% |
| 282 | Hcaps00086_SRR25580024 | Mississippiense | 2945 | 118032 | 29728336 | 20409 | 433 | 0 | 98.40% |
| 283 | Hcaps00010_SRR17307789 | Mississippiense | 2013 | 158061 | 30251973 | 32862 | 271 | 0 | 98.80% |
| 284 | Hcaps00083_SRR25580021 | Mississippiense | 1923 | 163772 | 29933585 | 34226 | 259 | 0 | 98.70% |
| 285 | Hcaps00041_SRR21059749 | Mississippiense | 1540 | 221524 | 30053579 | 43701 | 202 | 0 | 98.70% |
| 286 | Hcaps00247_SRR6243638 | Mississippiense | 2110 | 166253 | 29620323 | 31261 | 282 | 0 | 98.80% |
| 287 | Hcaps00243_SRR6243634 | Mississippiense | 2064 | 158251 | 29540470 | 32064 | 276 | 0 | 98.70% |
| 288 | Hcaps00245_SRR6243636 | Mississippiense | 1883 | 150829 | 29600911 | 36494 | 242 | 0 | 98.90% |
| 289 | Hcaps00244_SRR6243635 | Mississippiense | 2061 | 213053 | 29545221 | 33446 | 267 | 0 | 98.70% |
| 290 | Hcaps00119_SRR25580057 | Mississippiense | 2519 | 136752 | 29992252 | 24583 | 358 | 0 | 98.60% |
| 291 | Hcaps00257_SRR6243648 | Mississippiense | 1995 | 154322 | 29489249 | 34112 | 265 | 0 | 98.90% |
| 292 | Hcaps00271_SRR6243662 | Mississippiense | 2076 | 182954 | 29497403 | 32161 | 281 | 0 | 98.80% |
| 293 | Hcaps00250_SRR6243641 | Mississippiense | 1935 | 196186 | 29575350 | 35947 | 245 | 0 | 98.90% |
| 294 | Hcaps00242_SRR6243633 | Mississippiense | 1866 | 141790 | 29581130 | 36394 | 248 | 0 | 98.90% |
| 295 | Hcapsulatum_GCA017310585 | Mississippiense | 8 | 7576836 | 32531515 | 5328076 | 3 | 0 | 99.10% |
| 296 | Hcapsulatum_GCA000149585 | Mississippiense | 280 | 7343276 | 33030326 | 2885091 | 4 | 7208.57 | 93.50% |
| 297 | Hcaps00141_SRR27481867 | India | 3172 | 153305 | 29819834 | 25894 | 338 | 0 | 98.90% |
| 298 | Hcaps00150_SRR27481876 | India | 2932 | 153082 | 29800992 | 27226 | 328 | 0 | 98.90% |
| 299 | Hcaps00158_SRR31797080 | India | 2570 | 157443 | 29777477 | 32392 | 271 | 0 | 99.00% |
| 300 | Hcaps00146_SRR27481872 | India | 3031 | 153085 | 29741408 | 24492 | 350 | 0 | 99.00% |
| 301 | Hcaps00140_SRR27481866 | India | 3119 | 152893 | 29766041 | 26515 | 331 | 0 | 98.90% |
| 302 | Hcaps00152_SRR27481878 | India | 3192 | 153134 | 29705848 | 25704 | 338 | 0 | 98.90% |
| 303 | Hcaps00147_SRR27481873 | India | 3604 | 117914 | 29729728 | 23345 | 371 | 0 | 98.90% |
| 304 | Hcaps00145_SRR27481871 | India | 4230 | 113320 | 29560732 | 19074 | 446 | 0 | 98.50% |
| 305 | Hcaps00139_SRR27481865 | India | 3120 | 134945 | 29851050 | 26448 | 332 | 0 | 99.00% |
| 306 | Hcaps00143_SRR27481869 | India | 3152 | 134338 | 29847001 | 26688 | 331 | 0 | 98.90% |
| 307 | Hcaps00138_SRR27481864 | India | 3001 | 135012 | 29726479 | 26423 | 326 | 0 | 98.90% |
| 308 | Hcaps00151_SRR27481877 | India | 2733 | 136356 | 29462559 | 26716 | 321 | 0 | 98.90% |
| 309 | Hcaps00149_SRR27481875 | India | 2815 | 135046 | 29523001 | 27095 | 331 | 0 | 98.90% |
| 310 | Hcaps00142_SRR27481868 | India | 3078 | 134990 | 29596927 | 25910 | 343 | 0 | 98.90% |
| 311 | Hcaps00148_SRR27481874 | India | 3387 | 134968 | 29711769 | 24882 | 346 | 0 | 99.00% |

| | | | | | | | | | |
|---|---|---|---|---|---|---|---|---|---|
| 312 | Hcaps00137_SRR27481863 | India | 3261 | 134960 | 29887039 | 26032 | 337 | 0 | 98.90% |
| 313 | Hcaps00144_SRR27481870 | India | 9344 | 34461 | 29384500 | 5016 | 1774 | 0 | 84.50% |

**Outgroup genomes for tree rooting**

| Order | Species | Lineage | # contigs | Largest contig | Total length | N50 | L50 | #N's/100 kbp | BUSCO % |
|---|---|---|---|---|---|---|---|---|---|
| root | Blastomyces dermatitidis | outgroup | 3960 | 311976 | 62560380 | 43590 | 369 | 0 | 99.50% |
| root | Blastomyces parvus | outgroup | 181 | 1788946 | 27921030 | 623573 | 16 | 0 | 99.80% |
| root | Emergomyces pasteurianus | outgroup | 863 | 781711 | 32639620 | 149774 | 61 | 0 | 99.50% |
| root | Paracoccidioides brasiliensis | outgroup | 771 | 301564 | 29402480 | 84342 | 108 | 0 | 92.50% |

**Duplicate samples and derivatives of G217B/WU15**

| Order[3] | Assembly[2] | Lineage | # contigs | Largest contig | Total length | N50 | L50 | #N's/100 kbp | BUSCO % |
|---|---|---|---|---|---|---|---|---|---|
| 1 | Hcaps00306_SRR33400182 | Ohiense | 1217 | 441408 | 38228770 | 102235 | 111 | 0 | 99.40% |
| 2 | Hcaps00310_SRR33400186 | Ohiense | 1226 | 419972 | 38204702 | 106518 | 108 | 0 | 99.40% |
| 3 | Hcaps00308_SRR33400184 | Ohiense | 1307 | 440935 | 38213358 | 103456 | 107 | 0 | 99.40% |
| 4 | Hcaps00298_SRR33380105 | Ohiense | 1049 | 388237 | 38294403 | 107384 | 109 | 0 | 99.40% |
| 5 | Hcaps00302_SRR33380109 | Ohiense | 997 | 515775 | 38307353 | 110174 | 104 | 0 | 99.40% |
| 6 | Hcaps00303_SRR33380110 | Ohiense | 1187 | 441563 | 38262348 | 95881 | 115 | 0 | 99.40% |
| 7 | Hcaps00296_SRR33380103 | Ohiense | 985 | 477810 | 38295451 | 116665 | 97 | 0 | 99.40% |
| 8 | Hcaps00321_SRR33400197 | Ohiense | 1042 | 440805 | 38275190 | 111637 | 101 | 0 | 99.40% |
| 9 | Hcaps00311_SRR33400187 | Ohiense | 1173 | 478000 | 38215203 | 117852 | 92 | 0 | 99.30% |
| 10 | Hcaps00307_SRR33400183 | Ohiense | 1126 | 363863 | 38235527 | 108443 | 109 | 0 | 99.40% |
| 11 | Hcaps00309_SRR33400185 | Ohiense | 1179 | 495180 | 38228616 | 110097 | 105 | 0 | 99.40% |
| 12 | Hcaps00299_SRR33380106 | Ohiense | 996 | 441600 | 38288248 | 116824 | 99 | 0 | 99.30% |
| 13 | Hcaps00304_SRR33400180 | Ohiense | 988 | 495210 | 38249419 | 115340 | 104 | 0 | 99.40% |
| 14 | Hcaps00322_SRR33400198 | Ohiense | 1224 | 363892 | 38231390 | 107487 | 108 | 0 | 99.40% |
| 15 | Hcaps00305_SRR33400181 | Ohiense | 1170 | 413254 | 38208471 | 103648 | 109 | 0 | 99.30% |
| 16 | Hcaps00350_SRR33400226 | Ohiense | 1185 | 435756 | 38212541 | 103482 | 109 | 0 | 99.30% |
| 17 | Hcaps00372_SRR33400248 | Ohiense | 1197 | 441655 | 38210558 | 103503 | 110 | 0 | 99.40% |
| 18 | Hcaps00155_SRR31751697 | Ohiense | 934 | 549679 | 38279502 | 127610 | 90 | 0 | 99.40% |
| 19 | Hcaps00154_SRR31751696 | Ohiense | 1056 | 441631 | 38305164 | 110531 | 104 | 0 | 99.40% |
| 20 | Hcaps00153_SRR31751695 | Ohiense | 1000 | 372312 | 38281875 | 117944 | 100 | 0 | 99.40% |
| 21 | Hcaps00335_SRR33400211 | Ohiense | 1026 | 442158 | 38266376 | 114591 | 96 | 0 | 99.40% |
| 22 | Hcaps00356_SRR33400232 | Ohiense | 918 | 441624 | 38329201 | 123505 | 93 | 0 | 99.30% |
| 23 | Hcaps00336_SRR33400212 | Ohiense | 1040 | 500037 | 38271121 | 118197 | 99 | 0 | 99.30% |

| | | | | | | | | | |
|---|---|---|---|---|---|---|---|---|---|
| 24 | Hcaps00300_SRR33380107 | Ohiense | 978 | 441660 | 38290566 | 116332 | 98 | 0 | 99.30% |
| 25 | Hcaps00341_SRR33400217 | Ohiense | 1249 | 540455 | 38223455 | 104474 | 103 | 0 | 99.30% |
| 26 | Hcaps00349_SRR33400225 | Ohiense | 1155 | 428562 | 38231887 | 108918 | 105 | 0 | 99.40% |
| 27 | Hcaps00363_SRR33400239 | Ohiense | 1220 | 435752 | 38241745 | 106979 | 108 | 0 | 99.40% |
| 28 | Hcaps00334_SRR33400210 | Ohiense | 1062 | 441664 | 38283006 | 110557 | 102 | 0 | 99.30% |
| 29 | Hcaps00351_SRR33400227 | Ohiense | 1181 | 441632 | 38238539 | 101696 | 110 | 0 | 99.30% |
| 30 | Hcaps00370_SRR33400246 | Ohiense | 1305 | 519633 | 38208634 | 104135 | 104 | 0 | 99.30% |
| 31 | Hcaps00301_SRR33380108 | Ohiense | 1107 | 549774 | 38279034 | 111102 | 102 | 0 | 99.40% |
| 32 | Hcaps00365_SRR33400241 | Ohiense | 1161 | 435784 | 38257675 | 103039 | 110 | 0 | 99.40% |
| 33 | Hcaps00353_SRR33400229 | Ohiense | 1135 | 473456 | 38262809 | 94963 | 113 | 0 | 99.30% |
| 34 | Hcaps00352_SRR33400228 | Ohiense | 1126 | 441479 | 38247858 | 102536 | 108 | 0 | 99.40% |
| 35 | Hcaps00346_SRR33400222 | Ohiense | 1117 | 441715 | 38235708 | 112596 | 98 | 0 | 99.40% |
| 36 | Hcaps00358_SRR33400234 | Ohiense | 1446 | 549781 | 38166253 | 110533 | 92 | 0 | 99.40% |
| 37 | Hcaps00373_SRR33400249 | Ohiense | 1226 | 440935 | 38208857 | 93935 | 113 | 0 | 99.30% |
| 38 | Hcaps00354_SRR33400230 | Ohiense | 1241 | 435835 | 38222905 | 104497 | 107 | 0 | 99.40% |
| 39 | Hcaps00355_SRR33400231 | Ohiense | 1103 | 563898 | 38249683 | 116093 | 101 | 0 | 99.30% |
| 40 | Hcaps00329_SRR33400205 | Ohiense | 1126 | 435773 | 38234432 | 110098 | 103 | 0 | 99.40% |
| 41 | Hcaps00331_SRR33400207 | Ohiense | 1176 | 441686 | 38228595 | 108488 | 107 | 0 | 99.30% |
| 42 | Hcaps00337_SRR33400213 | Ohiense | 1085 | 443008 | 38268358 | 113561 | 99 | 0 | 99.30% |
| 43 | Hcaps00368_SRR33400244 | Ohiense | 1221 | 549692 | 38232125 | 110157 | 102 | 0 | 99.30% |
| 44 | Hcaps00369_SRR33400245 | Ohiense | 1149 | 426478 | 38255759 | 101668 | 112 | 0 | 99.40% |
| 45 | Hcaps00320_SRR33400196 | Ohiense | 1120 | 549729 | 38269214 | 104533 | 108 | 0 | 99.30% |
| 46 | Hcaps00366_SRR33400242 | Ohiense | 1178 | 435759 | 38224223 | 103901 | 108 | 0 | 99.40% |
| 47 | Hcaps00367_SRR33400243 | Ohiense | 1100 | 366281 | 38246017 | 111137 | 104 | 0 | 99.30% |
| 48 | Hcaps00371_SRR33400247 | Ohiense | 1322 | 441618 | 38218199 | 93772 | 115 | 0 | 99.40% |
| 49 | Hcaps00361_SRR33400237 | Ohiense | 1427 | 606220 | 38180825 | 96829 | 102 | 0 | 99.30% |
| 50 | Hcaps00360_SRR33400236 | Ohiense | 1654 | 416897 | 37995864 | 98354 | 111 | 0 | 99.40% |
| 51 | Hcaps00314_SRR33400190 | Ohiense | 1547 | 546677 | 38158713 | 95717 | 106 | 0 | 99.30% |
| 52 | Hcaps00330_SRR33400206 | Ohiense | 1370 | 839174 | 38194725 | 111981 | 100 | 0 | 99.40% |
| 53 | Hcaps00357_SRR33400233 | Ohiense | 1713 | 529373 | 38043226 | 93199 | 114 | 0 | 99.30% |
| 54 | Hcaps00359_SRR33400235 | Ohiense | 1474 | 564389 | 38180199 | 103383 | 102 | 0 | 99.30% |
| 55 | Hcaps00364_SRR33400240 | Ohiense | 1540 | 668381 | 38161668 | 104754 | 98 | 0 | 99.40% |
| 56 | Hcaps00313_SRR33400189 | Ohiense | 1488 | 636145 | 38147525 | 104088 | 100 | 0 | 99.40% |
| 57 | Hcaps00339_SRR33400215 | Ohiense | 2010 | 435651 | 38037748 | 99898 | 110 | 0 | 99.40% |
| 58 | Hcaps00338_SRR33400214 | Ohiense | 1690 | 605386 | 38101627 | 90380 | 107 | 0 | 99.30% |

| | | | | | | | | | |
|---|---|---|---|---|---|---|---|---|---|
| 59 | Hcaps00327_SRR33400203 | Ohiense | 2291 | 369941 | 38010495 | 67890 | 142 | 0 | 99.40% |
| 60 | Hcaps00325_SRR33400201 | Ohiense | 2278 | 374996 | 37988393 | 74856 | 132 | 0 | 99.40% |
| 61 | Hcaps00340_SRR33400216 | Ohiense | 1409 | 582981 | 38174891 | 105052 | 101 | 0 | 99.40% |
| 62 | Hcaps00323_SRR33400199 | Ohiense | 1493 | 584148 | 38145527 | 93734 | 112 | 0 | 99.40% |
| 63 | Hcaps00324_SRR33400200 | Ohiense | 1524 | 592603 | 38150547 | 98657 | 109 | 0 | 99.40% |
| 64 | Hcaps00326_SRR33400202 | Ohiense | 2022 | 444821 | 38065360 | 89267 | 120 | 0 | 99.40% |
| 65 | Hcaps00312_SRR33400188 | Ohiense | 1781 | 606167 | 38123324 | 93200 | 112 | 0 | 99.30% |
| 66 | Hcaps00342_SRR33400218 | Ohiense | 1510 | 563716 | 38125519 | 104841 | 98 | 0 | 99.30% |
| 67 | Hcaps00343_SRR33400219 | Ohiense | 1473 | 581649 | 38158116 | 108449 | 99 | 0 | 99.40% |
| 68 | Hcaps00332_SRR33400208 | Ohiense | 1633 | 442969 | 38099566 | 103685 | 103 | 0 | 99.40% |
| 69 | Hcaps00315_SRR33400191 | Ohiense | 1668 | 612112 | 38132842 | 92873 | 108 | 0 | 99.40% |
| 70 | Hcaps00319_SRR33400195 | Ohiense | 1482 | 435662 | 38161292 | 102808 | 106 | 0 | 99.30% |
| 71 | Hcaps00317_SRR33400193 | Ohiense | 2408 | 286213 | 38022209 | 62685 | 168 | 0 | 99.30% |
| 72 | Hcaps00318_SRR33400194 | Ohiense | 2387 | 363066 | 38004843 | 64460 | 158 | 0 | 99.40% |
| 73 | Hcaps00345_SRR33400221 | Ohiense | 3281 | 484081 | 36694998 | 67067 | 146 | 0 | 99.30% |
| 74 | Hcaps00003_SRR13433422 | Ohiense | 1186 | 369364 | 38319010 | 79244 | 142 | 0 | 99.30% |
| 75 | Hcaps00068_SRR24516103 | Ohiense | 3558 | 281386 | 31576682 | 58294 | 152 | 0 | 99.40% |
| 76 | Hcaps00070_SRR24516105 | Ohiense | 3629 | 322895 | 31367441 | 64830 | 139 | 0 | 99.30% |
| 77 | Hcaps00071_SRR24516106 | Ohiense | 3785 | 272047 | 31262968 | 57861 | 151 | 0 | 99.30% |
| 78 | Hcaps00069_SRR24516104 | Ohiense | 3743 | 283503 | 31224408 | 59907 | 151 | 0 | 99.20% |
| 79 | Hcaps00067_SRR24516102 | Ohiense | 3536 | 287145 | 31455819 | 69193 | 128 | 0 | 99.30% |
| 80 | Hcaps00066_SRR24516101 | Ohiense | 3606 | 355553 | 31396740 | 61582 | 144 | 0 | 99.30% |
| 81 | Hcaps00328_SRR33400204 | Ohiense | 3409 | 266485 | 33515517 | 51409 | 184 | 0 | 99.20% |
| 82 | Hcaps00270_SRR6243661 | Ohiense | 2097 | 222465 | 37189813 | 39270 | 260 | 0 | 99.00% |
| 83 | Hcaps00347_SRR33400223 | Ohiense | 1169 | 441610 | 38247822 | 101793 | 109 | 0 | 99.30% |
| 84 | Hcaps00295_SRR33380102 | Ohiense | 1016 | 393038 | 38294843 | 117580 | 96 | 0 | 99.30% |
| 85 | Hcaps00348_SRR33400224 | Ohiense | 1255 | 487727 | 38218796 | 95929 | 108 | 0 | 99.30% |
| 86 | Hcaps00157_SRR31751699 | Ohiense | 981 | 549730 | 38282612 | 120802 | 90 | 0 | 99.40% |
| 87 | Hcaps00297_SRR33380104 | Ohiense | 993 | 517319 | 38295396 | 114087 | 103 | 0 | 99.30% |
| 88 | Hcaps00344_SRR33400220 | Ohiense | 2095 | 361376 | 38050586 | 70859 | 139 | 0 | 99.40% |
| 89 | Hcaps00333_SRR33400209 | Ohiense | 1466 | 443073 | 38159206 | 111869 | 97 | 0 | 99.40% |
| 90 | Hcaps00316_SRR33400192 | Ohiense | 2296 | 442964 | 38034192 | 67846 | 148 | 0 | 99.40% |
| 91 | Hcaps00362_SRR33400238 | Ohiense | 1473 | 443121 | 38170804 | 95039 | 108 | 0 | 99.40% |
| 92 | Hcapsulatum_GCA017607445 | Ohiense | 12 | 8217122 | 39447273 | 6680169 | 3 | 0 | 99.20% |
| 93 | Hcaps00291_SRR8426586 | Ohiense | 4513 | 186422 | 31124409 | 27740 | 309 | 0 | 98.80% |

| | | | | | | | | | |
|---|---|---|---|---|---|---|---|---|---|
| 94 | Hcaps00156_SRR31751698 | Ohiense | 1003 | 549726 | 38298085 | 118172 | 100 | 0 | 99.30% |
| 95 | Hcapsulatum_GCA000170615 | Ohiense | 266 | 2014935 | 41285890 | 597872 | 22 | 0 | 99.40% |
| 96 | Hcaps00004_SRR13433424 | Capsu | 378 | 762042 | 30443794 | 164673 | 56 | 0 | 99.60% |
| 97 | Hcaps00006_SRR13508697 | Capsu | 445 | 610280 | 30293253 | 193026 | 51 | 0 | 99.60% |
| 98 | Hcapsulatum_GCA013420885 | Capsu | 285 | 3610552 | 30598358 | 1343918 | 7 | 1170.47 | 99.40% |
| 99 | Hcaps00259_SRR6243650 | Capsu | 511 | 512959 | 30400550 | 120619 | 80 | 0 | 99.60% |
| 100 | Hcapsulatum_GCA017607465 | Capsu | 12 | 9773074 | 30991520 | 4063218 | 3 | 0 | 99.70% |
| 101 | Hcapsulatum_GCA017355575 | Capsu | 7 | 7332157 | 31111494 | 5585295 | 3 | 0 | 99.70% |
| 102 | Hcapsulatum_GCA000150115 | Capsu | 88 | 3777229 | 30483324 | 1769553 | 6 | 660.58 | 99.30% |
| 103 | Hcapsulatum_GCA017310615 | Africa | 7 | 9359328 | 37996987 | 7000705 | 3 | 0 | 99.70% |
| 104 | Hcaps00002_SRR13433402 | Africa | 583 | 1096954 | 37312327 | 174701 | 65 | 0 | 99.70% |
| 105 | Hcapsulatum_GCA000151005 | Africa | 17 | 7050398 | 37943219 | 5001999 | 4 | 668.22 | 99.70% |
| 106 | Hcaps00005_SRR13508696 | Mississippiense | 1670 | 289703 | 29299085 | 58975 | 153 | 0 | 98.90% |
| 107 | Hcaps00249_SRR6243640 | Mississippiense | 2075 | 133601 | 29597718 | 31904 | 267 | 0 | 98.80% |
| 108 | Hcapsulatum_GCA017310585 | Mississippiense | 8 | 7576836 | 32531515 | 5328076 | 3 | 0 | 99.10% |
| 109 | Hcaps00141_SRR27481867 | India | 3172 | 153305 | 29819834 | 25894 | 338 | 0 | 98.90% |

**Excluded samples**

| Order | Assembly | Lineage | # contigs | Largest contig | Total length | N50 | L50 | #N's/100 kbp | BUSCO % |
|---|---|---|---|---|---|---|---|---|---|
| ND | Hcaps00388_SRR29905408 | Africa | 6425 | 231469 | 74961722 | 28866 | 653 | 0 | N/A |
| ND | Hcaps00171_SRR31893259 | Sura | 1626 | 912143 | 66979548 | 153047 | 115 | 0 | N/A |
| ND | Hcaps00216_SRR31893306 | Mz5-like | 1981 | 269790 | 69323015 | 67359 | 324 | 0 | N/A |

Footnotes:

1. Order of the samples in the wgMLST 0300 tree shown in Figure 3.
2. Assemblies labeled in light blue are the 60 assemblies used for generation of the wgMLST schemes and testing of the schemes (Fig. 2, S2, S3).
3. BUSCO percentage completenes based on the complete, the fragmented and the duplicate hits added up, compared to the 5,439 markers of the ajellomycetaceae_odb12.2 lineage database (Tegenfeldt et al., 2025).
4. Order of the samples in the wgMLST 0300 tree shown in Figure S4.

Cells highlighted in yellow indicate low quality assembly metrics (high number of N's, $N_{50}$ < 25kb, $L_{50}$ > 500, number of contigs > 5000)

An Excel spreadsheet version of this Table is available at Zenodo (doi: 10.5281/zenodo.22865196) and Figshare (doi: 10.6084/m9.figshare.33946246).

**Table S3.** Metadata of *Histoplasma capsulatum* samples/genome assemblies used in this study, including order in the wgMLST trees (300 nt minimum ORF length), accession and biosample numbers, isolation source, year of isolation, country/continent of isolation.

**Genomes used in main study**

| Order[1] | Assembly | (Sub)Lineage | SRA/ENA[2] | Assembly | Biosample[3] | Name | Host | Year | Country |
|---|---|---|---|---|---|---|---|---|---|
| 1 | Hcaps00164_SRR31893252 | Sura | SRR31893252 | GCA_988282835 | SAMEA123467319 | HC970591 | Human | 2021 | Venezuela |
| 2 | Hcaps00166_SRR31893254 | Sura | SRR31893254 | GCA_988282755 | SAMEA123467321 | HC4137 | Human | 2021 | Venezuela |
| 3 | Hcaps00170_SRR31893258 | Sura | SRR31893258 | GCA_988282815 | SAMEA123467324 | HC1070058-2 | Human | 2021 | Venezuela |
| 4 | Hcaps00290_SRR8347492 | Sura | SRR8347492 | GCA_988269055 | SAMN10132218 | Hc7909 | Human | 2000 | Venezuela |
| 5 | Hcaps00172_SRR31893260 | Sura | SRR31893260 | GCA_988282715 | SAMEA123467325 | HC7090 | Human | 2021 | Venezuela |
| 6 | Hcaps00173_SRR31893261 | Sura | SRR31893261 | GCA_988282345 | SAMEA123467326 | HC7072a | Human | 2021 | Venezuela |
| 7 | Hcaps00163_SRR31893251 | Sura | SRR31893251 | GCA_988282255 | SAMEA123467318 | hcjb | Human | 2021 | Venezuela |
| 8 | Hcaps00168_SRR31893256 | Sura | SRR31893256 | GCA_988271725 | SAMN46055077 | HC3066 | Human | 2021 | Venezuela |
| 9 | Hcaps00165_SRR31893253 | Sura | SRR31893253 | GCA_988283395 | SAMEA123467320 | HC776 | Human | 2021 | Venezuela |
| 10 | Hcaps00167_SRR31893255 | Sura | SRR31893255 | GCA_988282635 | SAMEA123467322 | HC394 | Human | 2021 | Venezuela |
| 11 | Hcaps00120_SRR25580058 | Sura | SRR25580058 | GCA_988269845 | SAMN36890447 | HistoB05614 | Human | missing | India |
| 12 | Hcaps00020_SRR21059728 | Sura | SRR21059728 | GCA_988269005 | SAMN30261154 | B16560 | Human | 2018 | USA |
| 13 | Hcaps00135_SRR25636145 | Sura | SRR25636145 | GCA_988270025 | SAMN36986136 | 600 | Human | 2020 | USA |
| 14 | Hcaps00014_SRR17307793 | Sura | SRR17307793 | GCA_988272655 | SAMN24252190 | H3 | Human | 2013 | Canada |
| 15 | Hcaps00177_SRR31893265 | Sura | SRR31893265 | GCA_988282645 | SAMEA123467330 | CM5679 | Human | 2009 | Spain |
| 16 | Hcaps00176_SRR31893264 | Sura | SRR31893264 | GCA_988282375 | SAMEA123467329 | CM5692 | Human | 2009 | Spain |
| 17 | Hcaps00254_SRR6243645 | Sura | SRR6243645 | GCA_988271805 | SAMN07964917 | 3_11 | Human | 2011 | Guatemala |
| 18 | Hcaps00178_SRR31893266 | Sura | SRR31893266 | GCA_988282985 | SAMEA123467331 | B06379 | Human | 2021 | Nicaragua |
| 19 | Hcaps00175_SRR31893263 | Sura | SRR31893263 | GCA_988282915 | SAMEA123467328 | CM7057 | Human | 2012 | Spain |
| 20 | Hcaps00252_SRR6243643 | Sura | SRR6243643 | GCA_988271875 | SAMN07964919 | 21_14 | Human | 2014 | Guatemala |
| 21 | Hcaps00110_SRR25580048 | Sura | SRR25580048 | GCA_988272195 | SAMN36890428 | Histo2134 | Human | missing | USA |
| 22 | Hcaps00007_SRR17307786 | Sura | SRR17307786 | GCA_988273305 | SAMN24252197 | H12 | Human | 2018 | Canada |
| 23 | Hcaps00174_SRR31893262 | Sura | SRR31893262 | GCA_988282455 | SAMEA123467327 | CM7717 | Human | 2015 | Spain |
| 24 | Hcaps00015_SRR17307794 | Sura | SRR17307794 | GCA_988268915 | SAMN24252198 | H13 | Human | 2019 | Canada |
| 25 | Hcaps00026_SRR21059734 | Sura | SRR21059734 | GCA_988269475 | SAMN30261191 | B18188 | Human | 2019 | USA |
| 26 | Hcaps00013_SRR17307792 | Sura | SRR17307792 | GCA_988270765 | SAMN24252191 | H4 | Human | 2015 | Canada |
| 27 | Hcaps00008_SRR17307787 | Sura | SRR17307787 | GCA_988270405 | SAMN24252196 | H11 | Human | 2018 | Canada |
| 28 | Hcaps00406_SRR29905426 | Farciminosum | SRR29905426 | GCA_988272675 | SAMN42690067 | 19VMG-15 | Horse | 2019 | South Africa |
| 29 | Hcaps00395_SRR29905415 | Farciminosum | SRR29905415 | GCA_988272625 | SAMN42690068 | 20VMK-11 | Horse | 2020 | South Africa |
| 30 | Hcaps00382_SRR29905402 | Farciminosum | SRR29905402 | GCA_988269375 | SAMN42690066 | CM7436 | Horse | 2023 | Egypt |

| | | | | | | | | | |
|---|---|---|---|---|---|---|---|---|---|
| 31 | Hcaps00105_SRR25580043 | Sura/Farci | SRR25580043 | GCA_988270035 | SAMN36890459 | HistoB10265 | Human | 2013 | USA |
| 32 | Hcaps00001_ERR7414368 | Sura/Farci | ERR7414368 | GCA_988283025 | SAMEA123467297 | 09-0599 | Human | 2009 | Germany |
| 33 | Hcapsulatum_GCA000313325 | Sura/Farci | N/A | GCA_000313325.1 | SAMN02981464 | Tmu | Human | 2011 | Taiwan |
| 34 | Hcaps00275_SRR8084707 | RJ | SRR8084707 | GCA_988271895 | SAMN10257874 | AC05 | soil | 1985 | Brazil |
| 35 | Hcaps00276_SRR8084708 | RJ | SRR8084708 | GCA_988269175 | SAMN10257881 | RPS35 | soil | 1984 | Brazil |
| 36 | Hcaps00277_SRR8084709 | RJ | SRR8084709 | GCA_988271925 | SAMN10257880 | CO4 | soil | 1984 | Brazil |
| 37 | Hcaps00273_SRR8084705 | RJ | SRR8084705 | GCA_988269085 | SAMN10257876 | RS36 | Mouse | 1981 | Brazil |
| 38 | Hcaps00274_SRR8084706 | RJ | SRR8084706 | GCA_988272685 | SAMN10257875 | IT04 | soil | 1983 | Brazil |
| 39 | Hcaps00272_SRR8084704 | RJ | SRR8084704 | GCA_988271325 | SAMN10257877 | TI01 | soil | 1984 | Brazil |
| 40 | Hcaps00282_SRR8084714 | RJ | SRR8084714 | GCA_988268985 | SAMN10257885 | 07_12 | Dog | 2012 | Brazil |
| 41 | Hcaps00284_SRR8084716 | RJ | SRR8084716 | GCA_988270725 | SAMN10257887 | 24_11 | Human | 2011 | Brazil |
| 42 | Hcaps00285_SRR8084717 | RJ | SRR8084717 | GCA_988269095 | SAMN10257886 | 28_11 | Human | 2011 | Brazil |
| 43 | Hcaps00289_SRR8084721 | RJ | SRR8084721 | GCA_988272045 | SAMN10257890 | 27_11 | Human | 2011 | Brazil |
| 44 | Hcaps00283_SRR8084715 | RJ | SRR8084715 | GCA_988271625 | SAMN10257884 | 23_11 | Human | 2011 | Brazil |
| 45 | Hcaps00288_SRR8084720 | RJ | SRR8084720 | GCA_988268865 | SAMN10257891 | 06_12 | Human | 2012 | Brazil |
| 46 | Hcaps00287_SRR8084719 | RJ | SRR8084719 | GCA_988269415 | SAMN10257888 | 02_14 | Human | 2014 | Brazil |
| 47 | Hcaps00278_SRR8084710 | RJ | SRR8084710 | GCA_988269045 | SAMN10257879 | 39439 | Human | 2000 | Brazil |
| 48 | Hcaps00193_SRR31893282 | Amazon-2 | SRR31893282 | GCA_988282955 | SAMEA123467346 | FGACO2005 | Human | 2013 | French Guiana |
| 49 | Hcaps00192_SRR31893281 | Amazon-2 | SRR31893281 | GCA_988282615 | SAMEA123467345 | FGBRU0291 | Human | 2006 | French Guiana |
| 50 | Hcaps00189_SRR31893278 | Amazon-2 | SRR31893278 | GCA_988283405 | SAMEA123467342 | SDOR2042 | Human | 2014 | Suriname |
| 51 | Hcaps00191_SRR31893280 | Amazon-2 | SRR31893280 | GCA_988282775 | SAMEA123467344 | FGPERS2034 | Human | 2016 | French Guiana |
| 52 | Hcaps00280_SRR8084712 | NEB | SRR8084712 | GCA_988269135 | SAMN10257883 | 09_12 | Human | 2012 | Brazil |
| 53 | Hcaps00281_SRR8084713 | NEB | SRR8084713 | GCA_988271815 | SAMN10257882 | 01_16 | Human | 2016 | Brazil |
| 54 | Hcaps00210_SRR31893300 | Mz5-like | SRR31893300 | GCA_988283325 | SAMEA123467363 | FGPIN2043 | Human | 2010 | French Guiana |
| 55 | Hcaps00196_SRR31893286 | Mz5-like | SRR31893286 | GCA_988282705 | SAMEA123467349 | FGZAA2004 | Human | 2007 | French Guiana |
| 56 | Hcaps00180_SRR31893268 | Mz5-like | SRR31893268 | GCA_988282305 | SAMEA123467333 | FGCLE2015 | Human | 2012 | French Guiana |
| 57 | Hcaps00199_SRR31893289 | Mz5-like | SRR31893289 | GCA_988282905 | SAMEA123467352 | FGBIK2051 | Human | 2017 | French Guiana |
| 58 | Hcaps00169_SRR31893257 | Mz5-like | SRR31893257 | GCA_988282725 | SAMEA123467323 | FGCRI2041 | Human | 2010 | French Guiana |
| 59 | Hcaps00203_SRR31893293 | Mz5-like | SRR31893293 | GCA_988283065 | SAMEA123467356 | FGVER2032 | Human | 2009 | French Guiana |
| 60 | Hcaps00198_SRR31893288 | Mz5-like | SRR31893288 | GCA_988283125 | SAMEA123467351 | SPA2057 | Human | 2015 | Suriname |
| 61 | Hcaps00201_SRR31893291 | Mz5-like | SRR31893291 | GCA_988283085 | SAMEA123467354 | SADA2068 | Human | 2014 | Suriname |
| 62 | Hcaps00222_SRR31893312 | Mz5-like | SRR31893312 | GCA_988282765 | SAMEA123467374 | FGAMA2041 | Human | 2010 | French Guiana |
| 63 | Hcaps00212_SRR31893302 | Mz5-like | SRR31893302 | GCA_988283385 | SAMEA123467365 | FGPIE2055 | Human | 2017 | French Guiana |
| 64 | Hcaps00218_SRR31893308 | Mz5-like | SRR31893308 | GCA_988283285 | SAMEA123467370 | FGJOS2044 | Human | 2016 | French Guiana |
| 65 | Hcaps00208_SRR31893298 | Mz5-like | SRR31893298 | GCA_988283175 | SAMEA123467361 | FGPOE2043 | Human | 2012 | French Guiana |

| | | | | | | | | | |
|---|---|---|---|---|---|---|---|---|---|
| 66 | Hcaps00217_SRR31893307 | Mz5-like | SRR31893307 | GCA_988283375 | SAMEA123467369 | FGKOU2024 | Human | 2017 | French Guiana |
| 67 | Hcaps00223_SRR31893313 | Mz5-like | SRR31893313 | GCA_988282315 | SAMEA123467375 | FGADA2079 | Human | 2013 | French Guiana |
| 68 | Hcaps00204_SRR31893294 | Mz5-like | SRR31893294 | GCA_988283095 | SAMEA123467357 | SMIS2065 | Human | 2014 | Suriname |
| 69 | Hcaps00161_SRR31893249 | Mz5-like | SRR31893249 | GCA_988282405 | SAMEA123467316 | FGDAS2062 | Human | 2013 | French Guiana |
| 70 | Hcaps00207_SRR31893297 | Mz5-like | SRR31893297 | GCA_988283145 | SAMEA123467360 | SASA2073 | Human | 2014 | Suriname |
| 71 | Hcaps00200_SRR31893290 | Mz5-like | SRR31893290 | GCA_988282895 | SAMEA123467353 | SASAAZP1 | Human | 2013 | Suriname |
| 72 | Hcaps00190_SRR31893279 | Mz5-like | SRR31893279 | GCA_988282335 | SAMEA123467343 | FGBON2001 | Human | 2015 | French Guiana |
| 73 | Hcaps00219_SRR31893309 | Mz5-like | SRR31893309 | GCA_988282505 | SAMEA123467371 | FGGRE2022 | Human | 2016 | French Guiana |
| 74 | Hcaps00211_SRR31893301 | Mz5-like | SRR31893301 | GCA_988283295 | SAMEA123467364 | FGBEL2002 | Human | 2017 | French Guiana |
| 75 | Hcaps00221_SRR31893311 | Mz5-like | SRR31893311 | GCA_988282355 | SAMEA123467373 | FGFAN2059 | Human | 2017 | French Guiana |
| 76 | Hcaps00215_SRR31893305 | Mz5-like | SRR31893305 | GCA_988282625 | SAMEA123467368 | FGMEL2036 | Human | 2007 | French Guiana |
| 77 | Hcaps00213_SRR31893303 | Mz5-like | SRR31893303 | GCA_988282515 | SAMEA123467366 | FGPIA2052 | Human | 2009 | French Guiana |
| 78 | Hcaps00202_SRR31893292 | Mz5-like | SRR31893292 | GCA_988283195 | SAMEA123467355 | FGZUL2036 | Human | 2009 | French Guiana |
| 79 | Hcaps00209_SRR31893299 | Mz5-like | SRR31893299 | GCA_988283155 | SAMEA123467362 | FGPIR2086 | Human | 2009 | French Guiana |
| 80 | Hcaps00160_SRR31893248 | Mz5-like | SRR31893248 | GCA_988283415 | SAMEA123467315 | FGDEE2016 | Human | 2010 | French Guiana |
| 81 | Hcaps00206_SRR31893296 | Mz5-like | SRR31893296 | GCA_988283075 | SAMEA123467359 | FGROD2046 | Human | 2010 | French Guiana |
| 82 | Hcaps00214_SRR31893304 | Mz5-like | SRR31893304 | GCA_988282675 | SAMEA123467367 | FGNON2028 | Human | 2017 | French Guiana |
| 83 | Hcaps00159_SRR31893247 | Mz5-like | SRR31893247 | GCA_988282685 | SAMEA123467314 | FGDEO2088 | Human | 2014 | French Guiana |
| 84 | Hcaps00228_SRR32121389 | Mz5-like | SRR32121389 | GCA_988283235 | SAMEA123467380 | 4337-Belem17 | Human | 2015 | Brazil |
| 85 | Hcaps00227_SRR32121388 | Mz5-like | SRR32121388 | GCA_988283115 | SAMEA123467379 | 4729-Belem16 | Human | 2016 | Brazil |
| 86 | Hcaps00241_SRR32121402 | Mz5-like | SRR32121402 | GCA_988282435 | SAMEA123467393 | 11571-Belem1 | Human | 2008 | Brazil |
| 87 | Hcaps00240_SRR32121401 | Mz5-like | SRR32121401 | GCA_988282465 | SAMEA123467392 | 11572-Belem2 | Human | 2008 | Brazil |
| 88 | Hcaps00226_SRR32121387 | Mz5-like | SRR32121387 | GCA_988283035 | SAMEA123467378 | 47692-Belem14 | Human | 2016 | Brazil |
| 89 | Hcaps00230_SRR32121391 | Mz5-like | SRR32121391 | GCA_988282265 | SAMEA123467382 | 3948-Belem7 | Human | 2015 | Brazil |
| 90 | Hcaps00220_SRR31893310 | Mz5-like | SRR31893310 | GCA_988282805 | SAMEA123467372 | FGFER2036 | Human | 2017 | French Guiana |
| 91 | Hcaps00231_SRR32121392 | Mz5-like | SRR32121392 | GCA_988282445 | SAMEA123467383 | 2202-Belem5 | Human | 2011 | Brazil |
| 92 | Hcaps00205_SRR31893295 | Mz5-like | SRR31893295 | GCA_988282995 | SAMEA123467358 | FGSOU0318 | Human | 2006 | French Guiana |
| 93 | Hcaps00229_SRR32121390 | Mz5-like | SRR32121390 | GCA_988282535 | SAMEA123467381 | 4336-Belem10 | Human | 2015 | Brazil |
| 94 | Hcaps00258_SRR6243649 | Mz5-like | SRR6243649 | GCA_988282285 | SAMEA123467395 | MZ5 | Human | missing | Colombia |
| 95 | Hcaps00374_SRR33700783 | Mz5-like | SRR33700783 | GCA_988282855 | SAMEA123467400 | CM5788 | Human | 2020 | Brazil |
| 96 | Hcaps00042_SRR21059750 | Mz5-like | SRR21059750 | GCA_988270085 | SAMN30261150 | B16384 | Human | 2018 | USA |
| 97 | Hcaps00195_SRR31893285 | Amazon-1 | SRR31893285 | GCA_988282875 | SAMEA123467348 | FGBRE2013 | Human | 2010 | French Guiana |
| 98 | Hcaps00194_SRR31893284 | Amazon-1 | SRR31893284 | GCA_988282975 | SAMEA123467347 | FGPIC2055 | Human | 2016 | French Guiana |
| 99 | Hcaps00294_SRR31893283 | Amazon-1 | SRR31893283 | GCA_988282525 | SAMEA123467399 | HCM-H | Human | 2021 | Venezuela |
| 100 | Hcaps00197_SRR31893287 | Amazon-1 | SRR31893287 | GCA_988283315 | SAMEA123467350 | HCAM | Human | 2021 | Venezuela |

| | | | | | | | | | |
|---|---|---|---|---|---|---|---|---|---|
| 101 | Hcaps00255_SRR6243646 | 27-14 | SRR6243646 | GCA_988268855 | SAMN07964918 | 27-14 | Human | 2014 | Guatemala |
| 102 | Hcaps00279_SRR8084711 | LAmB | SRR8084711 | GCA_988271765 | SAMN10257878 | 03_16 | Human | 2011 | Brazil |
| 103 | Hcaps00162_SRR31893250 | LAmB | SRR31893250 | GCA_988282695 | SAMEA123467317 | A290302130 | Human | 2009 | Martinique |
| 104 | Hcaps00286_SRR8084718 | LAmB | SRR8084718 | GCA_988271835 | SAMN10257889 | 11_12 | Human | 2012 | Brazil |
| 105 | Hcaps00400_SRR29905420 | Africa | SRR29905420 | GCA_988271465 | SAMN42690087 | B05181 | Human | 2023 | Cote d'Ivoire |
| 106 | Hcaps00402_SRR29905422 | Africa | SRR29905422 | GCA_988271105 | SAMN42690076 | SA811 | Human | 2015 | South Africa |
| 107 | Hcaps00391_SRR29905411 | Africa | SRR29905411 | GCA_988271235 | SAMN42690090 | ES2_84 | Human | 2023 | Cameroon |
| 108 | Hcaps00385_SRR29905405 | Africa | SRR29905405 | GCA_988270265 | SAMN42690096 | ES2_90 | Human | 2023 | Cote d'Ivoire |
| 109 | Hcaps00392_SRR29905412 | Africa | SRR29905412 | GCA_988273125 | SAMN42690089 | ES2_83 | Human | 2023 | South Africa |
| 110 | Hcaps00253_SRR6243644 | Africa | SRR6243644 | GCA_988282295 | SAMEA123467394 | DUBOISIIA | Human | missing | "Africa" |
| 111 | Hcapsulatum_GCA017310615 | Africa | N/A | GCA_017310615.1 | SAMN17004946 | H88 | Human | 1975 | Belgium |
| 112 | Hcaps00248_SRR6243639 | Africa | SRR6243639 | GCA_988268845 | SAMN07964921 | DUBOISIIB | Human | missing | "Africa" |
| 113 | Hcaps00396_SRR29905416 | Africa | SRR29905416 | GCA_988271015 | SAMN42690085 | CM7767 | Human | 2015 | Nigeria |
| 114 | Hcaps00393_SRR29905413 | Africa | SRR29905413 | GCA_988271285 | SAMN42690088 | ES2_82 | Human | 2023 | Cote d'Ivoire |
| 115 | Hcaps00375_SRR29905395 | Africa | SRR29905395 | GCA_988272105 | SAMN42690073 | SA355 | Human | 2014 | South Africa |
| 116 | Hcaps00379_SRR29905399 | Africa | SRR29905399 | GCA_988268885 | SAMN42690078 | SA1436 | Human | 2019 | South Africa |
| 117 | Hcaps00387_SRR29905407 | Africa | SRR29905407 | GCA_988270365 | SAMN42690094 | ES2_88 | Human | 2023 | Cameroon |
| 118 | Hcaps00390_SRR29905410 | Africa | SRR29905410 | GCA_988272125 | SAMN42690091 | ES2_85 | Human | 2023 | Cameroon |
| 119 | Hcaps00397_SRR29905417 | Africa | SRR29905417 | GCA_988271265 | SAMN42690084 | CM7704 | Human | 2015 | Ghana |
| 120 | Hcaps00404_SRR29905424 | Africa | SRR29905424 | GCA_988271085 | SAMN42690070 | SA47 | Human | 2010 | South Africa |
| 121 | Hcaps00405_SRR29905425 | Africa | SRR29905425 | GCA_988271025 | SAMN42690069 | SA46 | Human | 2010 | South Africa |
| 122 | Hcaps00381_SRR29905401 | Africa | SRR29905401 | GCA_988269275 | SAMN42690077 | SA19 | Human | 2023 | South Africa |
| 123 | Hcaps00384_SRR29905404 | Africa | SRR29905404 | GCA_988269355 | SAMN42690081 | SA1704 | Human | 2021 | South Africa |
| 124 | Hcaps00377_SRR29905397 | Africa | SRR29905397 | GCA_988269435 | SAMN42690071 | SA297 | Human | 2013 | South Africa |
| 125 | Hcaps00378_SRR29905398 | Africa | SRR29905398 | GCA_988271825 | SAMN42690080 | SA1556 | Human | 2020 | South Africa |
| 126 | Hcaps00398_SRR29905418 | Africa | SRR29905418 | GCA_988271065 | SAMN42690083 | CM7434 | Human | 2023 | South Africa |
| 127 | Hcapsulatum_GCA000151035 | Africa | N/A | GCA_000151035.1 | SAMN02953731 | H143 | Human | 1954 | South Africa |
| 128 | Hcaps00389_SRR29905409 | Africa | SRR29905409 | GCA_988271865 | SAMN42690092 | ES2_86 | Human | 2023 | Cameroon |
| 129 | Hcaps00383_SRR29905403 | Africa | SRR29905403 | GCA_988268965 | SAMN42690097 | ES2_91 | Human | 2023 | Gabon |
| 130 | Hcaps00403_SRR29905423 | Africa | SRR29905423 | GCA_988272755 | SAMN42690075 | SA602 | Human | 2015 | South Africa |
| 131 | Hcaps00376_SRR29905396 | Africa | SRR29905396 | GCA_988272115 | SAMN42690072 | SA302 | Human | 2013 | South Africa |
| 132 | Hcaps00394_SRR29905414 | Africa | SRR29905414 | GCA_988271165 | SAMN42690086 | CM8013 | Human | 2016 | Ghana |
| 133 | Hcaps00399_SRR29905419 | Africa | SRR29905419 | GCA_988272695 | SAMN42690082 | CM7001 | Human | 2012 | Eq. Guinea |
| 134 | Hcaps00401_SRR29905421 | Africa | SRR29905421 | GCA_988271045 | SAMN42690074 | SA562 | Human | 2014 | South Africa |
| 135 | Hcaps00380_SRR29905400 | Africa | SRR29905400 | GCA_988269265 | SAMN42690079 | SA1371 | Human | 2019 | South Africa |

| | | | | | | | | | |
|---|---|---|---|---|---|---|---|---|---|
| 136 | Hcaps00386_SRR29905406 | Africa | SRR29905406 | GCA_988271055 | SAMN42690095 | ES2_89 | Human | 2023 | DR Congo |
| 137 | Hcaps00182_SRR31893271 | Capsu | SRR31893271 | GCA_988282425 | SAMEA123467335 | STAN2075 | Human | 2014 | Suriname |
| 138 | Hcaps00183_SRR31893272 | Capsu | SRR31893272 | GCA_988282415 | SAMEA123467336 | SROB2039 | Human | 2014 | Suriname |
| 139 | Hcaps00185_SRR31893274 | Capsu | SRR31893274 | GCA_988282595 | SAMEA123467338 | FGWIL2021 | Human | 2008 | French Guiana |
| 140 | Hcaps00179_SRR31893267 | Capsu | SRR31893267 | GCA_988282555 | SAMEA123467332 | HCRP | Human | 2021 | Venezuela |
| 141 | Hcaps00232_SRR32121393 | Capsu | SRR32121393 | GCA_988282385 | SAMEA123467384 | CM6556 | Human | 2022 | Venezuela |
| 142 | Hcaps00181_SRR31893270 | Capsu | SRR31893270 | GCA_988282395 | SAMEA123467334 | hc3645 | Human | 2021 | Venezuela |
| 143 | Hcaps00186_SRR31893275 | Capsu | SRR31893275 | GCA_988283305 | SAMEA123467339 | FGSAT2037 | Human | 2009 | French Guiana |
| 144 | Hcaps00187_SRR31893276 | Capsu | SRR31893276 | GCA_988282845 | SAMEA123467340 | FGMARTIN | Human | 2017 | French Guiana |
| 145 | Hcaps00188_SRR31893277 | Capsu | SRR31893277 | GCA_988282865 | SAMEA123467341 | FGDOS0487 | Human | 2007 | French Guiana |
| 146 | Hcaps00293_SRR31893269 | Capsu | SRR31893269 | GCA_988282735 | SAMEA123467398 | HC970588 | Human | 2021 | Venezuela |
| 147 | Hcaps00184_SRR31893273 | Capsu | SRR31893273 | GCA_988282745 | SAMEA123467337 | GJJ | Human | 2015 | Guyana |
| 148 | Hcaps00261_SRR6243652 | Capsu | SRR6243652 | GCA_988282785 | SAMEA123467397 | MV3 | Human | 1981 | Colombia |
| 149 | Hcaps00259_SRR6243650 | Capsu | SRR6243650 | GCA_988269975 | SAMN07964914 | G186A | Human | 1967 | Panama |
| 150 | Hcaps00256_SRR6243647 | Capsu | SRR6243647 | GCA_988270225 | SAMN07964915 | G184A | Human | 1967 | Panama |
| 151 | Hcapsulatum_GCA017607465 | Capsu | N/A | GCA_017607465.1 | SAMN17004945 | G184AR | Human | 1967 | Panama |
| 152 | Hcaps00235_SRR32121396 | Capsu | SRR32121396 | GCA_988282545 | SAMEA123467387 | B05368 | Human | 2022 | Brazil |
| 153 | Hcaps00234_SRR32121395 | Capsu | SRR32121395 | GCA_988282795 | SAMEA123467386 | B05593-Histo79 | Human | 2022 | Panama |
| 154 | Hcapsulatum_GCA017355575 | Capsu | N/A | GCA_017355575.1 | SAMN17004902 | G186AR | Human | 1967 | Panama |
| 155 | Hcaps00233_SRR32121394 | Capsu | SRR32121394 | GCA_988282485 | SAMEA123467385 | CM6408 | Human | 2022 | Panama |
| 156 | Hcaps00237_SRR32121398 | Capsu | SRR32121398 | GCA_988282235 | SAMEA123467389 | 4809-Belem13 | Human | 2016 | Brazil |
| 157 | Hcaps00236_SRR32121397 | Capsu | SRR32121397 | GCA_988282495 | SAMEA123467388 | 52292-Belem19 | Human | 2016 | Brazil |
| 158 | Hcaps00238_SRR32121399 | Capsu | SRR32121399 | GCA_988282605 | SAMEA123467390 | 4182-Belem9 | Human | 2015 | Brazil |
| 159 | Hcaps00225_SRR32121386 | Capsu | SRR32121386 | GCA_988283205 | SAMEA123467377 | 1014-Belem3 | Human | 2020 | Brazil |
| 160 | Hcaps00224_SRR32121385 | Capsu | SRR32121385 | GCA_988283105 | SAMEA123467376 | 2363-Belem6 | Human | 2011 | Brazil |
| 161 | Hcaps00239_SRR32121400 | Capsu | SRR32121400 | GCA_988282325 | SAMEA123467391 | 3865-Belem8 | Human | 2015 | Brazil |
| 162 | Hcaps00046_SRR21059754 | Ohiense | SRR21059754 | GCA_988271545 | SAMN30261173 | B17708 | Human | 2019 | USA |
| 163 | Hcaps00057_SRR21059765 | Ohiense | SRR21059765 | GCA_988272765 | SAMN30261163 | B16716 | Human | 2018 | USA |
| 164 | Hcaps00056_SRR21059764 | Ohiense | SRR21059764 | GCA_988269805 | SAMN30261164 | B16717 | Human | 2018 | USA |
| 165 | Hcaps00062_SRR21059770 | Ohiense | SRR21059770 | GCA_988272945 | SAMN30261158 | B16689 | Human | 2018 | USA |
| 166 | Hcaps00054_SRR21059762 | Ohiense | SRR21059762 | GCA_988269525 | SAMN30261166 | B17545 | Human | 2019 | USA |
| 167 | Hcaps00040_SRR21059748 | Ohiense | SRR21059748 | GCA_988270995 | SAMN30261178 | B17927 | Human | 2019 | USA |
| 168 | Hcaps00055_SRR21059763 | Ohiense | SRR21059763 | GCA_988270655 | SAMN30261165 | B17544 | Human | 2019 | USA |
| 169 | Hcaps00031_SRR21059739 | Ohiense | SRR21059739 | GCA_988272215 | SAMN30261151 | B16385 | Human | 2018 | USA |
| 170 | Hcaps00407_SRR29905427 | Ohiense | SRR29905427 | GCA_988272475 | SAMN42690065 | SA15 | Human | 2023 | South Africa |

| | | | | | | | | | |
|---|---|---|---|---|---|---|---|---|---|
| 171 | Hcaps00019_SRR21059727 | Ohiense | SRR21059727 | GCA_988270905 | SAMN30261155 | B16680 | Human | 2018 | USA |
| 172 | Hcaps00018_SRR21059726 | Ohiense | SRR21059726 | GCA_988271955 | SAMN30261156 | B16687 | Human | 2018 | USA |
| 173 | Hcaps00045_SRR21059753 | Ohiense | SRR21059753 | GCA_988270155 | SAMN30261174 | B17812 | Human | 2019 | USA |
| 174 | Hcaps00063_SRR21059771 | Ohiense | SRR21059771 | GCA_988271515 | SAMN30261157 | B16688 | Human | 2018 | USA |
| 175 | Hcaps00048_SRR21059756 | Ohiense | SRR21059756 | GCA_988271525 | SAMN30261171 | B17633 | Human | 2019 | USA |
| 176 | Hcaps00059_SRR21059767 | Ohiense | SRR21059767 | GCA_988272905 | SAMN30261161 | B16714 | Human | 2018 | USA |
| 177 | Hcaps00052_SRR21059760 | Ohiense | SRR21059760 | GCA_988269695 | SAMN30261167 | B17546 | Human | 2019 | USA |
| 178 | Hcaps00021_SRR21059729 | Ohiense | SRR21059729 | GCA_988269395 | SAMN30261153 | B16530 | Human | 2018 | USA |
| 179 | Hcaps00053_SRR21059761 | Ohiense | SRR21059761 | GCA_988269635 | SAMN30261149 | B16383 | Human | 2018 | USA |
| 180 | Hcaps00049_SRR21059757 | Ohiense | SRR21059757 | GCA_988270425 | SAMN30261170 | B17565 | Human | 2019 | USA |
| 181 | Hcaps00111_SRR25580049 | Ohiense | SRR25580049 | GCA_988271475 | SAMN36890454 | HistoB06553 | Human | 2003 | USA |
| 182 | Hcaps00093_SRR25580031 | Ohiense | SRR25580031 | GCA_988272375 | SAMN36890470 | HistoB11773 | Human | 2013 | USA |
| 183 | Hcaps00043_SRR21059751 | Ohiense | SRR21059751 | GCA_988271495 | SAMN30261176 | B17925 | Human | 2019 | USA |
| 184 | Hcaps00047_SRR21059755 | Ohiense | SRR21059755 | GCA_988270205 | SAMN30261172 | B17666 | Human | 2019 | USA |
| 185 | Hcaps00039_SRR21059747 | Ohiense | SRR21059747 | GCA_988272165 | SAMN30261179 | B17928 | Human | 2019 | USA |
| 186 | Hcaps00061_SRR21059769 | Ohiense | SRR21059769 | GCA_988272745 | SAMN30261159 | B16710 | Human | 2018 | USA |
| 187 | Hcaps00123_SRR25580061 | Ohiense | SRR25580061 | GCA_988270435 | SAMN36890427 | Histo1009_H18 | Human | 2021 | USA |
| 188 | Hcaps00132_SRR25580070 | Ohiense | SRR25580070 | GCA_988270325 | SAMN36890436 | Histo2767_H130 | Human | 2020 | USA |
| 189 | Hcaps00099_SRR25580037 | Ohiense | SRR25580037 | GCA_988272435 | SAMN36890429 | Histo2311 | Human | 2003 | USA |
| 190 | Hcaps00072_SRR25580010 | Ohiense | SRR25580010 | GCA_988269215 | SAMN36890434 | Histo2436_H86 | Human | 1974 | USA |
| 191 | Hcaps00065_SRR21059773 | Ohiense | SRR21059773 | GCA_988269245 | SAMN30261147 | B15363 | Human | 2018 | USA |
| 192 | Hcaps00106_SRR25580044 | Ohiense | SRR25580044 | GCA_988270175 | SAMN36890458 | HistoB09607 | Human | 2012 | USA |
| 193 | Hcaps00121_SRR25580059 | Ohiense | SRR25580059 | GCA_988271585 | SAMN36890446 | HistoB05583 | Human | missing | USA |
| 194 | Hcaps00094_SRR25580032 | Ohiense | SRR25580032 | GCA_988272185 | SAMN36890469 | HistoB11347 | Human | 2012 | USA |
| 195 | Hcaps00075_SRR25580013 | Ohiense | SRR25580013 | GCA_988269035 | SAMN36890486 | HistoB9348 | Human | missing | USA |
| 196 | Hcaps00116_SRR25580054 | Ohiense | SRR25580054 | GCA_988272285 | SAMN36890426 | Histo1008 | Human | missing | USA |
| 197 | Hcaps00108_SRR25580046 | Ohiense | SRR25580046 | GCA_988271995 | SAMN36890456 | HistoB08830 | Human | 2010 | USA |
| 198 | Hcaps00115_SRR25580053 | Ohiense | SRR25580053 | GCA_988271225 | SAMN36890450 | HistoB06279 | Human | missing | USA |
| 199 | Hcaps00118_SRR25580056 | Ohiense | SRR25580056 | GCA_988269905 | SAMN36890449 | HistoB06278 | Human | missing | USA |
| 200 | Hcaps00107_SRR25580045 | Ohiense | SRR25580045 | GCA_988272535 | SAMN36890457 | HistoB08855 | Cat | 2010 | USA |
| 201 | Hcaps00022_SRR21059730 | Ohiense | SRR21059730 | GCA_988270805 | SAMN30261152 | B16524 | Human | 2018 | USA |
| 202 | Hcaps00077_SRR25580015 | Ohiense | SRR25580015 | GCA_988270305 | SAMN36890431 | Histo2317 | Human | missing | USA |
| 203 | Hcaps00088_SRR25580026 | Ohiense | SRR25580026 | GCA_988269785 | SAMN36890430 | Histo2314 | Human | 2014 | USA |
| 204 | Hcaps00050_SRR21059758 | Ohiense | SRR21059758 | GCA_988272145 | SAMN30261169 | B17564 | Human | 2019 | USA |
| 205 | Hcaps00064_SRR21059772 | Ohiense | SRR21059772 | GCA_988271685 | SAMN30261148 | B16336 | Human | 2018 | USA |

| | | | | | | | | | |
|---|---|---|---|---|---|---|---|---|---|
| 206 | Hcaps00098_SRR25580036 | Ohiense | SRR25580036 | GCA_988272235 | SAMN36890465 | HistoB11020 | Human | 2014 | USA |
| 207 | Hcaps00267_SRR6243658 | Ohiense | SRR6243658 | GCA_988268835 | SAMN07964935 | CI_9 | Human | 2016 | USA |
| 208 | Hcaps00114_SRR25580052 | Ohiense | SRR25580052 | GCA_988269875 | SAMN36890451 | HistoB06301 | Human | missing | USA |
| 209 | Hcaps00095_SRR25580033 | Ohiense | SRR25580033 | GCA_988269885 | SAMN36890468 | HistoB11303 | Human | 2010 | USA |
| 210 | Hcaps00129_SRR25580067 | Ohiense | SRR25580067 | GCA_988270145 | SAMN36890439 | HistoB04440 | Human | 2020 | USA |
| 211 | Hcaps00101_SRR25580039 | Ohiense | SRR25580039 | GCA_988273115 | SAMN36890463 | HistoB10907 | Human | 2014 | USA |
| 212 | Hcaps00262_SRR6243653 | Ohiense | SRR6243653 | GCA_988270095 | SAMN07964940 | CI_35 | Human | 1983 | USA |
| 213 | Hcaps00265_SRR6243656 | Ohiense | SRR6243656 | GCA_988271335 | SAMN07964937 | CI_17 | Human | 1984 | USA |
| 214 | Hcaps00266_SRR6243657 | Ohiense | SRR6243657 | GCA_988271315 | SAMN07964936 | CI_10 | Human | 2011 | USA |
| 215 | Hcaps00246_SRR6243637 | Ohiense | SRR6243637 | GCA_988269295 | SAMN07964923 | CI_4 | Human | missing | USA |
| 216 | Hcaps00264_SRR6243655 | Ohiense | SRR6243655 | GCA_988270185 | SAMN07964938 | CI_18 | Human | 1984 | USA |
| 217 | Hcaps00263_SRR6243654 | Ohiense | SRR6243654 | GCA_988271115 | SAMN07964939 | CI_30 | Human | 1985 | USA |
| 218 | Hcaps00127_SRR25580065 | Ohiense | SRR25580065 | GCA_988270505 | SAMN36890441 | HistoB04447 | Human | 2020 | USA |
| 219 | Hcaps00260_SRR6243651 | Ohiense | SRR6243651 | GCA_988282965 | SAMEA123467396 | Hc1986 | Human | 1984 | Colombia |
| 220 | Hcapsulatum_GCA017607445 | Ohiense | N/A | GCA_017607445.1 | SAMN17004899 | G217B | Human | 1973 | USA |
| 221 | Hcaps00269_SRR6243660 | Ohiense | SRR6243660 | GCA_988272915 | SAMN07964933 | G222B | Human | 2016 | USA |
| 222 | Hcaps00130_SRR25580068 | Ohiense | SRR25580068 | GCA_988270215 | SAMN36890438 | HistoB02798 | Human | 2020 | USA |
| 223 | Hcaps00128_SRR25580066 | Ohiense | SRR25580066 | GCA_988269285 | SAMN36890440 | HistoB04443 | Human | 2020 | USA |
| 224 | Hcaps00117_SRR25580055 | Ohiense | SRR25580055 | GCA_988269765 | SAMN36890425 | Histo1005 | Human | missing | USA |
| 225 | Hcaps00103_SRR25580041 | Ohiense | SRR25580041 | GCA_988271445 | SAMN36890461 | HistoB10351 | Human | 2013 | USA |
| 226 | Hcaps00060_SRR21059768 | Ohiense | SRR21059768 | GCA_988271575 | SAMN30261160 | B16711 | Human | 2018 | USA |
| 227 | Hcaps00036_SRR21059744 | Ohiense | SRR21059744 | GCA_988270885 | SAMN30261182 | B18179 | Human | 2019 | USA |
| 228 | Hcaps00133_SRR25580071 | Ohiense | SRR25580071 | GCA_988270235 | SAMN36890435 | Histo2472_H97 | Human | 2020 | USA |
| 229 | Hcaps00100_SRR25580038 | Ohiense | SRR25580038 | GCA_988271145 | SAMN36890464 | HistoB10920 | Human | 2014 | USA |
| 230 | Hcaps00268_SRR6243659 | Ohiense | SRR6243659 | GCA_988272995 | SAMN07964934 | CI_6 | Human | 2012 | USA |
| 231 | Hcaps00038_SRR21059746 | Ohiense | SRR21059746 | GCA_988272395 | SAMN30261180 | B17958 | Human | missing | USA |
| 232 | Hcaps00037_SRR21059745 | Ohiense | SRR21059745 | GCA_988273015 | SAMN30261181 | B17959 | Human | missing | USA |
| 233 | Hcaps00051_SRR21059759 | Ohiense | SRR21059759 | GCA_988270635 | SAMN30261168 | B17551 | Human | 2019 | USA |
| 234 | Hcaps00035_SRR21059743 | Ohiense | SRR21059743 | GCA_988272175 | SAMN30261183 | B18180 | Human | 2019 | USA |
| 235 | Hcaps00096_SRR25580034 | Ohiense | SRR25580034 | GCA_988272485 | SAMN36890467 | HistoB11298 | Human | 2010 | USA |
| 236 | Hcaps00122_SRR25580060 | Ohiense | SRR25580060 | GCA_988271655 | SAMN36890445 | HistoB05581 | Human | 2022 | USA |
| 237 | Hcaps00080_SRR25580018 | Ohiense | SRR25580018 | GCA_988271195 | SAMN36890482 | HistoB15421 | Dog | missing | USA |
| 238 | Hcaps00131_SRR25580069 | Ohiense | SRR25580069 | GCA_988271605 | SAMN36890437 | HistoB02529 | Human | 2020 | USA |
| 239 | Hcaps00024_SRR21059732 | Ohiense | SRR21059732 | GCA_988272455 | SAMN30261193 | B18192 | Human | 2019 | USA |
| 240 | Hcaps00058_SRR21059766 | Ohiense | SRR21059766 | GCA_988269165 | SAMN30261162 | B16715 | Human | 2018 | USA |

| | | | | | | | | | |
|---|---|---|---|---|---|---|---|---|---|
| 241 | Hcaps00034_SRR21059742 | Ohiense | SRR21059742 | GCA_988270705 | SAMN30261184 | B18181 | Human | 2019 | USA |
| 242 | Hcaps00027_SRR21059735 | Ohiense | SRR21059735 | GCA_988271425 | SAMN30261190 | B18187 | Human | 2019 | USA |
| 243 | Hcaps00025_SRR21059733 | Ohiense | SRR21059733 | GCA_988272835 | SAMN30261192 | B18190 | Human | 2019 | USA |
| 244 | Hcaps00030_SRR21059738 | Ohiense | SRR21059738 | GCA_988270935 | SAMN30261187 | B18184 | Human | 2019 | USA |
| 245 | Hcaps00032_SRR21059740 | Ohiense | SRR21059740 | GCA_988269505 | SAMN30261186 | B18183 | Human | 2019 | USA |
| 246 | Hcaps00033_SRR21059741 | Ohiense | SRR21059741 | GCA_988272465 | SAMN30261185 | B18182 | Human | 2019 | USA |
| 247 | Hcaps00028_SRR21059736 | Ohiense | SRR21059736 | GCA_988272665 | SAMN30261189 | B18186 | Human | 2019 | USA |
| 248 | Hcaps00029_SRR21059737 | Ohiense | SRR21059737 | GCA_988269795 | SAMN30261188 | B18185 | Human | 2019 | USA |
| 249 | Hcaps00023_SRR21059731 | Ohiense | SRR21059731 | GCA_988270715 | SAMN30261194 | B18193 | Human | 2019 | USA |
| 250 | Hcaps00104_SRR25580042 | Ohiense | SRR25580042 | GCA_988272315 | SAMN36890460 | HistoB10349 | Human | 2013 | USA |
| 251 | Hcaps00011_SRR17307790 | Ohiense | SRR17307790 | GCA_988270415 | SAMN24252193 | H6 | Human | 2016 | Canada |
| 252 | Hcaps00017_SRR17307796 | Ohiense | SRR17307796 | GCA_988273095 | SAMN24252188 | H1 | Human | 2003 | Canada |
| 253 | Hcaps00016_SRR17307795 | Ohiense | SRR17307795 | GCA_988272605 | SAMN24252189 | H2 | Human | 2011 | Canada |
| 254 | Hcaps00009_SRR17307788 | Ohiense | SRR17307788 | GCA_988270795 | SAMN24252195 | H8 | Human | 2018 | Canada |
| 255 | Hcaps00012_SRR17307791 | Ohiense | SRR17307791 | GCA_988270845 | SAMN24252192 | H5 | Human | 2016 | Canada |
| 256 | Hcaps00097_SRR25580035 | Ohiense | SRR25580035 | GCA_988269855 | SAMN36890466 | HistoB11242 | Human | 2006 | USA |
| 257 | Hcaps00124_SRR25580062 | Ohiense | SRR25580062 | GCA_988269105 | SAMN36890444 | HistoB05474 | Human | 2022 | USA |
| 258 | Hcaps00073_SRR25580011 | Ohiense | SRR25580011 | GCA_988269445 | SAMN36890433 | Histo2434_H84 | Oppossum | missing | USA |
| 259 | Hcaps00109_SRR25580047 | Ohiense | SRR25580047 | GCA_988270465 | SAMN36890455 | HistoB06895 | Human | 2006 | USA |
| 260 | Hcaps00074_SRR25580012 | Ohiense | SRR25580012 | GCA_988268925 | SAMN36890432 | Histo2404_H77 | Rat | missing | USA |
| 261 | Hcaps00044_SRR21059752 | Ohiense | SRR21059752 | GCA_988271645 | SAMN30261175 | B17836 | Human | 2019 | USA |
| 262 | Hcaps00113_SRR25580051 | Ohiense | SRR25580051 | GCA_988271415 | SAMN36890452 | HistoB06377 | Human | missing | USA |
| 263 | Hcaps00084_SRR25580022 | Mississippiense | SRR25580022 | GCA_988269515 | SAMN36890478 | HistoB15141 | Cat | 2015 | USA |
| 264 | Hcaps00076_SRR25580014 | Mississippiense | SRR25580014 | GCA_988272075 | SAMN36890485 | HistoB15424 | Cat | missing | USA |
| 265 | Hcaps00089_SRR25580027 | Mississippiense | SRR25580027 | GCA_988269565 | SAMN36890474 | HistoB13424 | Dog | 2014 | USA |
| 266 | Hcaps00081_SRR25580019 | Mississippiense | SRR25580019 | GCA_988273135 | SAMN36890481 | HistoB15414 | Dog | 2016 | USA |
| 267 | Hcaps00087_SRR25580025 | Mississippiense | SRR25580025 | GCA_988270575 | SAMN36890475 | HistoB13425 | Dog | missing | USA |
| 268 | Hcaps00082_SRR25580020 | Mississippiense | SRR25580020 | GCA_988271215 | SAMN36890480 | HistoB15413 | Dog | 2015 | USA |
| 269 | Hcaps00125_SRR25580063 | Mississippiense | SRR25580063 | GCA_988270455 | SAMN36890443 | HistoB05371 | Human | 2020 | USA |
| 270 | Hcaps00091_SRR25580029 | Mississippiense | SRR25580029 | GCA_988270535 | SAMN36890472 | HistoB12410-2 | Cat | 2013 | USA |
| 271 | Hcaps00092_SRR25580030 | Mississippiense | SRR25580030 | GCA_988272245 | SAMN36890471 | HistoB12410-1 | Cat | 2013 | USA |
| 272 | Hcaps00085_SRR25580023 | Mississippiense | SRR25580023 | GCA_988271965 | SAMN36890477 | HistoB13438 | Cat | 2015 | USA |
| 273 | Hcaps00079_SRR25580017 | Mississippiense | SRR25580017 | GCA_988271205 | SAMN36890483 | HistoB15422 | Cat | missing | USA |
| 274 | Hcaps00078_SRR25580016 | Mississippiense | SRR25580016 | GCA_988270295 | SAMN36890484 | HistoB15423 | Cat | missing | USA |
| 275 | Hcaps00102_SRR25580040 | Mississippiense | SRR25580040 | GCA_988272865 | SAMN36890462 | HistoB10816 | Human | 2014 | USA |

| | | | | | | | | | |
|---|---|---|---|---|---|---|---|---|---|
| 276 | Hcaps00136_SRR25758805 | Mississippiense | SRR25758805 | GCA_988270335 | SAMN37147750 | 1170 | Human | 2020 | USA |
| 277 | Hcaps00090_SRR25580028 | Mississippiense | SRR25580028 | GCA_988269575 | SAMN36890473 | HistoB12411 | Cat | 2014 | USA |
| 278 | Hcaps00126_SRR25580064 | Mississippiense | SRR25580064 | GCA_988268905 | SAMN36890442 | HistoB05316 | Human | 2020 | USA |
| 279 | Hcaps00251_SRR6243642 | Mississippiense | SRR6243642 | GCA_988269065 | SAMN07964930 | UCLA_531 | Human | missing | USA |
| 280 | Hcaps00134_SRR25636130 | Mississippiense | SRR25636130 | GCA_988271745 | SAMN36986150 | 1009 | Human | 2020 | USA |
| 281 | Hcaps00112_SRR25580050 | Mississippiense | SRR25580050 | GCA_988273075 | SAMN36890453 | HistoB06378 | Human | missing | USA |
| 282 | Hcaps00086_SRR25580024 | Mississippiense | SRR25580024 | GCA_988269545 | SAMN36890476 | HistoB13435 | Dog | 2014 | USA |
| 283 | Hcaps00010_SRR17307789 | Mississippiense | SRR17307789 | GCA_988270595 | SAMN24252194 | H7 | Human | 2018 | Canada |
| 284 | Hcaps00083_SRR25580021 | Mississippiense | SRR25580021 | GCA_988271035 | SAMN36890479 | HistoB15376 | Cat | 2015 | USA |
| 285 | Hcaps00041_SRR21059749 | Mississippiense | SRR21059749 | GCA_988270055 | SAMN30261177 | B17926 | Human | 2019 | USA |
| 286 | Hcaps00247_SRR6243638 | Mississippiense | SRR6243638 | GCA_988269405 | SAMN07964924 | CI_7 | Human | missing | USA |
| 287 | Hcaps00243_SRR6243634 | Mississippiense | SRR6243634 | GCA_988268935 | SAMN07964928 | CI_42 | Human | missing | USA |
| 288 | Hcaps00245_SRR6243636 | Mississippiense | SRR6243636 | GCA_988271665 | SAMN07964926 | CI_22 | Human | missing | USA |
| 289 | Hcaps00244_SRR6243635 | Mississippiense | SRR6243635 | GCA_988269225 | SAMN07964925 | CI_19 | Human | missing | USA |
| 290 | Hcaps00119_SRR25580057 | Mississippiense | SRR25580057 | GCA_988269585 | SAMN36890448 | HistoB05822 | Human | missing | USA |
| 291 | Hcaps00257_SRR6243648 | Mississippiense | SRR6243648 | GCA_988269965 | SAMN07964916 | 505 | Human | missing | missing |
| 292 | Hcaps00271_SRR6243662 | Mississippiense | SRR6243662 | GCA_988271985 | SAMN07964931 | DOWNS | Human | 2011 | USA |
| 293 | Hcaps00250_SRR6243641 | Mississippiense | SRR6243641 | GCA_988271775 | SAMN07964929 | CI_43 | Human | missing | USA |
| 294 | Hcaps00242_SRR6243633 | Mississippiense | SRR6243633 | GCA_988269335 | SAMN07964927 | CI_24 | Human | missing | USA |
| 295 | Hcapsulatum_GCA017310585 | Mississippiense | N/A | GCA_017310585.1 | SAMN17004911 | WU24 | Human | 2004 | USA |
| 296 | Hcapsulatum_GCA000149585 | Mississippiense | N/A | GCA_000149585.1 | SAMN02953633 | NAm1 | Human | missing | USA |
| 297 | Hcaps00141_SRR27481867 | India | SRR27481867 | GCA_988283135 | SAMEA123467302 | 136_P_07_S6 | Human | 2022 | India |
| 298 | Hcaps00150_SRR27481876 | India | SRR27481876 | GCA_988283345 | SAMEA123467311 | 327_P_12_S7 | Human | 2021 | India |
| 299 | Hcaps00158_SRR31797080 | India | SRR31797080 | GCA_988271735 | SAMN41842359 | Histo-485P20 | Human | 2021 | India |
| 300 | Hcaps00146_SRR27481872 | India | SRR27481872 | GCA_988282925 | SAMEA123467307 | WGS_S14 | Human | 2020 | India |
| 301 | Hcaps00140_SRR27481866 | India | SRR27481866 | GCA_988283045 | SAMEA123467301 | 144_p_08_S14 | Human | 2020 | India |
| 302 | Hcaps00152_SRR27481878 | India | SRR27481878 | GCA_988282565 | SAMEA123467313 | 104_p_06_S19 | Human | 2021 | India |
| 303 | Hcaps00147_SRR27481873 | India | SRR27481873 | GCA_988282245 | SAMEA123467308 | WGS_S11 | Human | 2021 | India |
| 304 | Hcaps00145_SRR27481871 | India | SRR27481871 | GCA_988283015 | SAMEA123467306 | WGS_S16 | Human | 2022 | India |
| 305 | Hcaps00139_SRR27481865 | India | SRR27481865 | GCA_988283005 | SAMEA123467300 | 1517_p_17_S20 | Human | 2020 | India |
| 306 | Hcaps00143_SRR27481869 | India | SRR27481869 | GCA_988283185 | SAMEA123467304 | 117_p_12_S17 | Human | 2022 | India |
| 307 | Hcaps00138_SRR27481864 | India | SRR27481864 | GCA_988283055 | SAMEA123467299 | 256_P_18_S2 | Human | 2020 | India |
| 308 | Hcaps00151_SRR27481877 | India | SRR27481877 | GCA_988282665 | SAMEA123467312 | 104_P_19_S5 | Human | 2021 | India |
| 309 | Hcaps00149_SRR27481875 | India | SRR27481875 | GCA_988282275 | SAMEA123467310 | 343_p_18_S1 | Human | 2021 | India |
| 310 | Hcaps00142_SRR27481868 | India | SRR27481868 | GCA_988283265 | SAMEA123467303 | 122_p_10_B_S15 | Human | 2022 | India |

| | | | | | | | | | |
|---|---|---|---|---|---|---|---|---|---|
| 311 | Hcaps00148_SRR27481874 | India | SRR27481874 | GCA_988282575 | SAMEA123467309 | 388_p_11_S16 | Human | 2021 | India |
| 312 | Hcaps00137_SRR27481863 | India | SRR27481863 | GCA_988282885 | SAMEA123467298 | 316_p_10_S18 | Human | 2020 | India |
| 313 | Hcaps00144_SRR27481870 | India | SRR27481870 | GCA_988283165 | SAMEA123467305 | 107_P_06_S1 | Human | 2022 | India |

**Outgroup genomes for tree rooting**

| Order | Species | Lineage | SRA/ENA | Assembly | Biosample | Name | Host | Year | Country |
|---|---|---|---|---|---|---|---|---|---|
| root | Blastomyces dermatitidis | outgroup | SRR20306113 | GCA_988264835 | SAMN29815857 | D19b | Human | 1990 | China |
| root | Blastomyces parvus | outgroup | SRR6147078 | GCA_988264845 | SAMN02854802 | UAMH_130 | missing | missing | missing |
| root | Emergomyces pasteurianus | outgroup | SRR1575175 | GCA_988264875 | SAMN02584703 | Ep9510 | missing | missing | missing |
| root | Paracoccidioides brasiliensis | outgroup | SRR12032014 | GCA_988264885 | SAMN05171542 | Pb03 | Human | missing | Brazil |

**Duplicate samples and derivatives of G217B/WU15**

| Order[4] | Assembly | Lineage | SRA/ENA | Assembly | Biosample | Name | Description[5] | Year | Country |
|---|---|---|---|---|---|---|---|---|---|
| 1 | Hcaps00306_SRR33400182 | Ohiense | SRR33400182 | GCA_988265625 | SAMN48284832 | G217BMix8 | Duplicate G217B | N/A | N/A |
| 2 | Hcaps00310_SRR33400186 | Ohiense | SRR33400186 | GCA_988265735 | SAMN48284828 | G217BMix4 | Duplicate G217B | N/A | N/A |
| 3 | Hcaps00308_SRR33400184 | Ohiense | SRR33400184 | GCA_988265485 | SAMN48284830 | G217BMix6 | Duplicate G217B | N/A | N/A |
| 4 | Hcaps00298_SRR33380105 | Ohiense | SRR33380105 | GCA_988265515 | SAMN48208354 | SHU-A-F | Mutant G217B | N/A | N/A |
| 5 | Hcaps00302_SRR33380109 | Ohiense | SRR33380109 | GCA_988265655 | SAMN48208353 | SHU-C-E | Mutant G217B | N/A | N/A |
| 6 | Hcaps00303_SRR33380110 | Ohiense | SRR33380110 | GCA_988265695 | SAMN48208352 | SHU-C-A | Mutant G217B | N/A | N/A |
| 7 | Hcaps00296_SRR33380103 | Ohiense | SRR33380103 | GCA_988265815 | SAMN48208357 | SHU-B-C | Mutant G217B | N/A | N/A |
| 8 | Hcaps00321_SRR33400197 | Ohiense | SRR33400197 | GCA_988266015 | SAMN48284826 | G217BMix2 | Duplicate G217B | N/A | N/A |
| 9 | Hcaps00311_SRR33400187 | Ohiense | SRR33400187 | GCA_988266045 | SAMN48284827 | G217BMix3 | Duplicate G217B | N/A | N/A |
| 10 | Hcaps00307_SRR33400183 | Ohiense | SRR33400183 | GCA_988265355 | SAMN48284831 | G217BMix7 | Duplicate G217B | N/A | N/A |
| 11 | Hcaps00309_SRR33400185 | Ohiense | SRR33400185 | GCA_988265405 | SAMN48284829 | G217BMix5 | Duplicate G217B | N/A | N/A |
| 12 | Hcaps00299_SRR33380106 | Ohiense | SRR33380106 | GCA_988265605 | SAMN48208356 | SHU-B-A | Mutant G217B | N/A | N/A |
| 13 | Hcaps00304_SRR33400180 | Ohiense | SRR33400180 | GCA_988265465 | SAMN48284834 | G217BMix10 | Duplicate G217B | N/A | N/A |
| 14 | Hcaps00322_SRR33400198 | Ohiense | SRR33400198 | GCA_988265775 | SAMN48284825 | G217BMix1 | Duplicate G217B | N/A | N/A |
| 15 | Hcaps00305_SRR33400181 | Ohiense | SRR33400181 | GCA_988265495 | SAMN48284833 | G217BMix9 | Duplicate G217B | N/A | N/A |
| 16 | Hcaps00350_SRR33400226 | Ohiense | SRR33400226 | GCA_988266155 | SAMN48284769 | Z18R | Mutant G217B | N/A | N/A |
| 17 | Hcaps00372_SRR33400248 | Ohiense | SRR33400248 | GCA_988266115 | SAMN48284768 | Z1S | Mutant G217B | N/A | N/A |
| 18 | Hcaps00155_SRR31751697 | Ohiense | SRR31751697 | GCA_988265805 | SAMN45877844 | R6_u_05 | Mutant G217B | N/A | N/A |
| 19 | Hcaps00154_SRR31751696 | Ohiense | SRR31751696 | GCA_988265645 | SAMN45877845 | AVR9_u_01 | Mutant G217B | N/A | N/A |
| 20 | Hcaps00153_SRR31751695 | Ohiense | SRR31751695 | GCA_988265385 | SAMN45877846 | u_u | Mutant G217B | N/A | N/A |
| 21 | Hcaps00335_SRR33400211 | Ohiense | SRR33400211 | GCA_988266085 | SAMN48284793 | X11R | Mutant G217B | N/A | N/A |
| 22 | Hcaps00356_SRR33400232 | Ohiense | SRR33400232 | GCA_988265925 | SAMN48284836 | WU15_S | Mutant G217B | N/A | N/A |
| 23 | Hcaps00336_SRR33400212 | Ohiense | SRR33400212 | GCA_988265905 | SAMN48284792 | Z18S | Mutant G217B | N/A | N/A |

| | | | | | | | | | |
|---|---|---|---|---|---|---|---|---|---|
| 24 | Hcaps00300_SRR33380107 | Ohiense | SRR33380107 | GCA_988265595 | SAMN48208358 | URA5-SB | Mutant G217B | N/A | N/A |
| 25 | Hcaps00341_SRR33400217 | Ohiense | SRR33400217 | GCA_988266225 | SAMN48284772 | A23S | Mutant G217B | N/A | N/A |
| 26 | Hcaps00349_SRR33400225 | Ohiense | SRR33400225 | GCA_988266295 | SAMN48284787 | C2S | Mutant G217B | N/A | N/A |
| 27 | Hcaps00363_SRR33400239 | Ohiense | SRR33400239 | GCA_988266285 | SAMN48284771 | A20S | Mutant G217B | N/A | N/A |
| 28 | Hcaps00334_SRR33400210 | Ohiense | SRR33400210 | GCA_988265895 | SAMN48284794 | A12R | Mutant G217B | N/A | N/A |
| 29 | Hcaps00351_SRR33400227 | Ohiense | SRR33400227 | GCA_988266335 | SAMN48284786 | B13R | Mutant G217B | N/A | N/A |
| 30 | Hcaps00370_SRR33400246 | Ohiense | SRR33400246 | GCA_988266165 | SAMN48284778 | B5S | Mutant G217B | N/A | N/A |
| 31 | Hcaps00301_SRR33380108 | Ohiense | SRR33380108 | GCA_988265675 | SAMN48208360 | SH-Agr-YB | Mutant G217B | N/A | N/A |
| 32 | Hcaps00365_SRR33400241 | Ohiense | SRR33400241 | GCA_988265955 | SAMN48284783 | B2R | Mutant G217B | N/A | N/A |
| 33 | Hcaps00353_SRR33400229 | Ohiense | SRR33400229 | GCA_988265995 | SAMN48284784 | B3R | Mutant G217B | N/A | N/A |
| 34 | Hcaps00352_SRR33400228 | Ohiense | SRR33400228 | GCA_988265975 | SAMN48284785 | B12R | Mutant G217B | N/A | N/A |
| 35 | Hcaps00346_SRR33400222 | Ohiense | SRR33400222 | GCA_988266355 | SAMN48284790 | C15R | Mutant G217B | N/A | N/A |
| 36 | Hcaps00358_SRR33400234 | Ohiense | SRR33400234 | GCA_988266195 | SAMN48284811 | L12P | Mutant G217B | N/A | N/A |
| 37 | Hcaps00373_SRR33400249 | Ohiense | SRR33400249 | GCA_988266065 | SAMN48284767 | X11S | Mutant G217B | N/A | N/A |
| 38 | Hcaps00354_SRR33400230 | Ohiense | SRR33400230 | GCA_988266315 | SAMN48284776 | B3S | Mutant G217B | N/A | N/A |
| 39 | Hcaps00355_SRR33400231 | Ohiense | SRR33400231 | GCA_988266305 | SAMN48284775 | B2S | Mutant G217B | N/A | N/A |
| 40 | Hcaps00329_SRR33400205 | Ohiense | SRR33400205 | GCA_988266025 | SAMN48284774 | A23R | Mutant G217B | N/A | N/A |
| 41 | Hcaps00331_SRR33400207 | Ohiense | SRR33400207 | GCA_988266125 | SAMN48284770 | A12S | Mutant G217B | N/A | N/A |
| 42 | Hcaps00337_SRR33400213 | Ohiense | SRR33400213 | GCA_988266145 | SAMN48284791 | C19R | Mutant G217B | N/A | N/A |
| 43 | Hcaps00368_SRR33400244 | Ohiense | SRR33400244 | GCA_988266255 | SAMN48284780 | B12S | Mutant G217B | N/A | N/A |
| 44 | Hcaps00369_SRR33400245 | Ohiense | SRR33400245 | GCA_988266135 | SAMN48284779 | B6S | Mutant G217B | N/A | N/A |
| 45 | Hcaps00320_SRR33400196 | Ohiense | SRR33400196 | GCA_988265745 | SAMN48284773 | A28S | Mutant G217B | N/A | N/A |
| 46 | Hcaps00366_SRR33400242 | Ohiense | SRR33400242 | GCA_988266075 | SAMN48284782 | B15S | Mutant G217B | N/A | N/A |
| 47 | Hcaps00367_SRR33400243 | Ohiense | SRR33400243 | GCA_988266245 | SAMN48284781 | B13S | Mutant G217B | N/A | N/A |
| 48 | Hcaps00371_SRR33400247 | Ohiense | SRR33400247 | GCA_988266215 | SAMN48284777 | B4S | Mutant G217B | N/A | N/A |
| 49 | Hcaps00361_SRR33400237 | Ohiense | SRR33400237 | GCA_988266345 | SAMN48284808 | L16N | Mutant G217B | N/A | N/A |
| 50 | Hcaps00360_SRR33400236 | Ohiense | SRR33400236 | GCA_988266325 | SAMN48284809 | K13R | Mutant G217B | N/A | N/A |
| 51 | Hcaps00314_SRR33400190 | Ohiense | SRR33400190 | GCA_988265705 | SAMN48284803 | K11S | Mutant G217B | N/A | N/A |
| 52 | Hcaps00330_SRR33400206 | Ohiense | SRR33400206 | GCA_988265965 | SAMN48284797 | E8S | Mutant G217B | N/A | N/A |
| 53 | Hcaps00357_SRR33400233 | Ohiense | SRR33400233 | GCA_988265915 | SAMN48284812 | L16Q | Mutant G217B | N/A | N/A |
| 54 | Hcaps00359_SRR33400235 | Ohiense | SRR33400235 | GCA_988266055 | SAMN48284810 | L7O | Mutant G217B | N/A | N/A |
| 55 | Hcaps00364_SRR33400240 | Ohiense | SRR33400240 | GCA_988266035 | SAMN48284806 | L7L | Mutant G217B | N/A | N/A |
| 56 | Hcaps00313_SRR33400189 | Ohiense | SRR33400189 | GCA_988265635 | SAMN48284804 | K13S | Mutant G217B | N/A | N/A |
| 57 | Hcaps00339_SRR33400215 | Ohiense | SRR33400215 | GCA_988266005 | SAMN48284818 | 4_Ms_Smooth | Mutant G217B | N/A | N/A |
| 58 | Hcaps00338_SRR33400214 | Ohiense | SRR33400214 | GCA_988265985 | SAMN48284819 | 5_Ms_Smooth | Mutant G217B | N/A | N/A |

| | | | | | | | | | |
|---|---|---|---|---|---|---|---|---|---|
| 59 | Hcaps00327_SRR33400203 | Ohiense | SRR33400203 | GCA_988265535 | SAMN48284820 | 6_Ms_Rough | Mutant G217B | N/A | N/A |
| 60 | Hcaps00325_SRR33400201 | Ohiense | SRR33400201 | GCA_988265615 | SAMN48284822 | 8_Ms_Rough | Mutant G217B | N/A | N/A |
| 61 | Hcaps00340_SRR33400216 | Ohiense | SRR33400216 | GCA_988266175 | SAMN48284817 | 3_Ms_Smooth | Mutant G217B | N/A | N/A |
| 62 | Hcaps00323_SRR33400199 | Ohiense | SRR33400199 | GCA_988265585 | SAMN48284824 | 10_Ms_Rough | Mutant G217B | N/A | N/A |
| 63 | Hcaps00324_SRR33400200 | Ohiense | SRR33400200 | GCA_988265665 | SAMN48284823 | 9_Ms_Rough | Mutant G217B | N/A | N/A |
| 64 | Hcaps00326_SRR33400202 | Ohiense | SRR33400202 | GCA_988265445 | SAMN48284821 | 7_Ms_Rough | Mutant G217B | N/A | N/A |
| 65 | Hcaps00312_SRR33400188 | Ohiense | SRR33400188 | GCA_988265525 | SAMN48284805 | K14S | Mutant G217B | N/A | N/A |
| 66 | Hcaps00342_SRR33400218 | Ohiense | SRR33400218 | GCA_988266105 | SAMN48284816 | 2_Ms_Smooth | Mutant G217B | N/A | N/A |
| 67 | Hcaps00343_SRR33400219 | Ohiense | SRR33400219 | GCA_988266375 | SAMN48284815 | 1_Ms_Smooth | Mutant G217B | N/A | N/A |
| 68 | Hcaps00332_SRR33400208 | Ohiense | SRR33400208 | GCA_988265875 | SAMN48284796 | D16S | Mutant G217B | N/A | N/A |
| 69 | Hcaps00315_SRR33400191 | Ohiense | SRR33400191 | GCA_988265755 | SAMN48284802 | E8R | Mutant G217B | N/A | N/A |
| 70 | Hcaps00319_SRR33400195 | Ohiense | SRR33400195 | GCA_988265935 | SAMN48284798 | D4R | Mutant G217B | N/A | N/A |
| 71 | Hcaps00317_SRR33400193 | Ohiense | SRR33400193 | GCA_988265565 | SAMN48284800 | D12R | Mutant G217B | N/A | N/A |
| 72 | Hcaps00318_SRR33400194 | Ohiense | SRR33400194 | GCA_988265365 | SAMN48284799 | D11R | Mutant G217B | N/A | N/A |
| 73 | Hcaps00345_SRR33400221 | Ohiense | SRR33400221 | GCA_988266265 | SAMN48284813 | D | Mutant G217B | N/A | N/A |
| 74 | Hcaps00003_SRR13433422 | Ohiense | SRR13433422 | GCA_988265845 | SAMN17004899 | G217B | Duplicate G217B | N/A | N/A |
| 75 | Hcaps00068_SRR24516103 | Ohiense | SRR24516103 | GCA_988265765 | SAMN35051543 | G217B_BJAS01 | Mutant G217B | N/A | N/A |
| 76 | Hcaps00070_SRR24516105 | Ohiense | SRR24516105 | GCA_988265395 | SAMN35051541 | G217B_BJAS01 | Mutant G217B | N/A | N/A |
| 77 | Hcaps00071_SRR24516106 | Ohiense | SRR24516106 | GCA_988265725 | SAMN35051540 | G217B_BJAS01 | Mutant G217B | N/A | N/A |
| 78 | Hcaps00069_SRR24516104 | Ohiense | SRR24516104 | GCA_988265835 | SAMN35051542 | G217B_BJAS01 | Mutant G217B | N/A | N/A |
| 79 | Hcaps00067_SRR24516102 | Ohiense | SRR24516102 | GCA_988265825 | SAMN35051544 | G217B_BJAS01 | Mutant G217B | N/A | N/A |
| 80 | Hcaps00066_SRR24516101 | Ohiense | SRR24516101 | GCA_988265475 | SAMN35051545 | G217B_BJAS01 | Mutant G217B | N/A | N/A |
| 81 | Hcaps00328_SRR33400204 | Ohiense | SRR33400204 | GCA_988265715 | SAMN48284835 | WU15_R | Mutant G217B | N/A | N/A |
| 82 | Hcaps00270_SRR6243661 | Ohiense | SRR6243661 | GCA_988265415 | SAMN07964932 | G217B | Duplicate G217B | N/A | N/A |
| 83 | Hcaps00347_SRR33400223 | Ohiense | SRR33400223 | GCA_988265945 | SAMN48284789 | C19S | Mutant G217B | N/A | N/A |
| 84 | Hcaps00295_SRR33380102 | Ohiense | SRR33380102 | GCA_988265795 | SAMN48208359 | URA5 | Mutant G217B | N/A | N/A |
| 85 | Hcaps00348_SRR33400224 | Ohiense | SRR33400224 | GCA_988266095 | SAMN48284788 | C15S | Mutant G217B | N/A | N/A |
| 86 | Hcaps00157_SRR31751699 | Ohiense | SRR31751699 | GCA_988265375 | SAMN45877842 | R10_u_02 | Mutant G217B | N/A | N/A |
| 87 | Hcaps00297_SRR33380104 | Ohiense | SRR33380104 | GCA_988266385 | SAMN48208355 | SHU-A-G | Mutant G217B | N/A | N/A |
| 88 | Hcaps00344_SRR33400220 | Ohiense | SRR33400220 | GCA_988266235 | SAMN48284814 | L | Mutant G217B | N/A | N/A |
| 89 | Hcaps00333_SRR33400209 | Ohiense | SRR33400209 | GCA_988265885 | SAMN48284795 | D4S | Mutant G217B | N/A | N/A |
| 90 | Hcaps00316_SRR33400192 | Ohiense | SRR33400192 | GCA_988265685 | SAMN48284801 | D16R | Mutant G217B | N/A | N/A |
| 91 | Hcaps00362_SRR33400238 | Ohiense | SRR33400238 | GCA_988266365 | SAMN48284807 | L12M | Mutant G217B | N/A | N/A |
| 92 | Hcapsulatum_GCA017607445 | Ohiense | N/A | GCA_017607445.1 | SAMN17004899 | G217B | Comparator | N/A | N/A |
| 93 | Hcaps00291_SRR8426586 | Ohiense | SRR8426586 | GCA_988265425 | SAMN10720093 | S29 | Mutant G217B | N/A | N/A |

| | | | | | | | | | |
|---|---|---|---|---|---|---|---|---|---|
| 94 | Hcaps00156_SRR31751698 | Ohiense | SRR31751698 | GCA_988265785 | SAMN45877843 | R1_u_02 | Mutant G217B | N/A | N/A |
| 95 | Hcapsulatum_GCA000170615 | Ohiense | N/A | GCA_000170615.1 | SAMN02953691 | G217B | Duplicate G217B | N/A | N/A |
| 96 | Hcaps00004_SRR13433424 | Capsu | SRR13433424 | GCA_988265505 | SAMN17004945 | G184AR | Duplicate G184 | N/A | N/A |
| 97 | Hcaps00006_SRR13508697 | Capsu | SRR13508697 | GCA_988265575 | SAMN17004902 | G186AR | Duplicate G186 | N/A | N/A |
| 98 | Hcapsulatum_GCA013420885 | Capsu | N/A | GCA_013420885.1 | SAMN14558693 | G186A | Duplicate G186 | N/A | N/A |
| 99 | Hcaps00259_SRR6243650 | Capsu | SRR6243650 | GCA_988269975 | SAMN07964914 | G186A | Comparator | N/A | N/A |
| 100 | Hcapsulatum_GCA017607465 | Capsu | N/A | GCA_017607465.1 | SAMN17004945 | G184AR | Comparator | N/A | N/A |
| 101 | Hcapsulatum_GCA017355575 | Capsu | N/A | GCA_017355575.1 | SAMN17004902 | G186AR | Comparator | N/A | N/A |
| 102 | Hcapsulatum_GCA000150115 | Capsu | N/A | GCA_000150115.1 | SAMN02953690 | G186AR | Duplicate G186 | N/A | N/A |
| 103 | Hcapsulatum_GCA017310615 | Africa | N/A | GCA_017310615.1 | SAMN17004946 | H88 | Comparator | N/A | N/A |
| 104 | Hcaps00002_SRR13433402 | Africa | SRR13433402 | GCA_988265455 | SAMN17004946 | H88 | Duplicate H88 | N/A | N/A |
| 105 | Hcapsulatum_GCA000151005 | Africa | N/A | GCA_000151005.2 | SAMN02953730 | H88 | Duplicate H88 | N/A | N/A |
| 106 | Hcaps00005_SRR13508696 | Mississippiense | SRR13508696 | GCA_988265545 | SAMN17004911 | WU24 | Duplicate WU24 | N/A | N/A |
| 107 | Hcaps00249_SRR6243640 | Mississippiense | SRR6243640 | GCA_988265865 | SAMN07964922 | WU24 | Duplicate WU24 | N/A | N/A |
| 108 | Hcapsulatum_GCA017310585 | Mississippiense | N/A | GCA_017310585.1 | SAMN17004911 | WU24 | Comparator | N/A | N/A |
| 109 | Hcaps00141_SRR27481867 | India | SRR27481867 | GCA_988283135 | SAMEA123467302 | 136_P_07_S6 | Root | N/A | N/A |

**Excluded samples**

| **Order** | **Assembly** | **Lineage** | **SRA/ENA** | **Assembly** | **Biosample** | **Name** | **Host** | **Year** | **Country** |
|---|---|---|---|---|---|---|---|---|---|
| ND | Hcaps00388_SRR29905408 | Africa | SRR29905408 | N/A | SAMN42690093 | ES2_87 | Human | 2023 | DR Congo |
| ND | Hcaps00171_SRR31893259 | Sura | SRR31893259 | N/A | SAMN46055075 | SDIJ2058 | Human | 2015 | Suriname |
| ND | Hcaps00216_SRR31893306 | Mz5-like | SRR31893306 | N/A | SAMN46055032 | FGLIN2055 | Human | 2014 | French Guiana |

Footnotes:

5. Order of the samples in the wgMLST0300 tree shown in Figure 3.
6. SRA/ENA codes are only shown for samples used for assembly generation in this study. N/A represent Genbank-derived assemblies.
7. New biosample numbers were generated for samples failing validation for ENA submission. The Zenodo/Figshare versions of Table S3 contain both biosample codes.
8. Order of the samples in the wgMLST 0300 tree shown in Figure S4.
9. The derivatives of G217B are primarily mutant isolates of *H. capsulatum* WU15 (G217B *ura5*$^{-}$) (Heater et al, 2026a, 2026b).

An Excel spreadsheet version of this Table is available at Zenodo (doi: 10.5281/zenodo.22865196) and Figshare (doi: 10.6084/m9.figshare.33946246).

**Table S4.** Information on the number of annotated protein coding genes for *Histoplasma capsulatum* genomes annotated with Funannotate, and annotated *H. capsulatum* genomes downloaded from NCBI Genome.

**Annotation metrics for genomes used for the fanMLST schemes (Funannotate)**

| Assembly | Lineage | #CDS | Annotation (nt) | Total Genome (nt) | Coding % |
|---|---|---|---|---|---|
| Hcaps00176_SRR31893264 | Sura | 7,778 | 12,933,019 | 33,280,898 | 38.9% |
| Hcaps00289_SRR8084721 | RJ | 7,708 | 12,882,388 | 31,460,371 | 40.9% |
| Hcaps00189_SRR31893278 | Amazon-2 | 7,629 | 12,836,044 | 29,738,317 | 43.2% |
| Hcaps00281_SRR8084713 | NEB | 7,523 | 13,304,562 | 31,935,425 | 41.7% |
| Hcaps00228_SRR32121389 | Mz5-like | 7,500 | 13,305,638 | 32,928,890 | 40.4% |
| Hcaps00195_SRR31893285 | Amazon-1 | 7,736 | 12,846,195 | 33,182,976 | 38.7% |
| Hcaps00255_SRR6243646 | 27-14 | 7,512 | 13,261,487 | 32,305,380 | 41.1% |
| Hcaps00279_SRR8084711 | LAmB | 7,775 | 12,911,046 | 29,977,273 | 43.1% |
| Hcapsulatum_GCA017310615 | Africa | 7,643 | 13,452,255 | 37,996,987 | 35.4% |
| Hcapsulatum_GCA017355575 | Capsulatum | 7,643 | 13,558,782 | 31,111,494 | 43.6% |
| Hcapsulatum_GCA017607445 | Ohiense | 7,819 | 13,346,019 | 39,447,273 | 33.8% |
| Hcapsulatum_GCA017310585 | Mississippiense | 8,927 | 14,022,220 | 32,531,515 | 43.1% |
| Hcaps00158_SRR31797080 | India | 8,711 | 13,548,231 | 29,777,477 | 45.5% |
| **average** | | **7,839** | **13,246,760** | **32,744,175** | **40.7%** |

**Annotation metrics for genomes used for the annMLST schemes (NCBI Genome)**

| Assembly | Lineage | #CDS | Annotation (nt) | Total Genome (nt) | Coding % |
|---|---|---|---|---|---|
| Hcapsulatum_GCA017310615 | Africa | 9,312 | 12,406,274 | 37,996,987 | 32.7% |
| Hcapsulatum_GCA017355575 | Capsulatum | 9,462 | 12,407,544 | 31,111,494 | 39.9% |
| Hcapsulatum_GCA017607465 | Capsulatum | 9,415 | 12,335,760 | 30,991,520 | 39.8% |
| Hcapsulatum_GCA051313235 | Ohiense | 9,524 | 12,748,783 | 39,447,273 | 32.3% |
| Hcapsulatum_GCA017310585 | Mississippiense | 9,080 | 13,796,061 | 32,531,515 | 42.4% |
| **average** | | **9,359** | **12,738,884** | **34,415,758** | **37.4%** |

An Excel spreadsheet version of this Table is available at Zenodo (doi: 10.5281/zenodo.22865196) and Figshare (doi: 10.6084/m9.figshare.33946246).

**Table S5.** Order of samples in the phylogenetic trees shown in Figures 2, 3, S2, S4, S5-S7.

**Positions of genomes in trees shown in Figures 2 and S3.**

| Node_names | Lineage | ParSNP | wg0300 | wg0500 | ann0300 | ann0500 | fan0300 | fan0500 |
|---|---|---|---|---|---|---|---|---|
| Hcaps00218_SRR31893308 | Mz5-like | 1 | 2 | 16 | 14 | 3 | 16 | 16 |
| Hcaps00210_SRR31893300 | Mz5-like | 2 | 5 | 18 | 16 | 5 | 18 | 18 |
| Hcaps00200_SRR31893290 | Mz5-like | 3 | 8 | 21 | 19 | 8 | 21 | 21 |
| Hcaps00223_SRR31893313 | Mz5-like | 4 | 1 | 15 | 13 | 1 | 15 | 15 |
| Hcaps00213_SRR31893303 | Mz5-like | 5 | 4 | 17 | 15 | 4 | 17 | 17 |
| Hcaps00221_SRR31893311 | Mz5-like | 6 | 3 | 14 | 12 | 2 | 14 | 14 |
| Hcaps00230_SRR32121391 | Mz5-like | 7 | 6 | 19 | 17 | 6 | 19 | 19 |
| Hcaps00228_SRR32121389 | Mz5-like | 8 | 7 | 20 | 18 | 7 | 20 | 20 |
| Hcaps00294_SRR31893283 | Amazon-1 | 9 | 9 | 23 | 20 | 9 | 22 | 22 |
| Hcaps00195_SRR31893285 | Amazon-1 | 10 | 10 | 22 | 21 | 10 | 23 | 23 |
| Hcaps00289_SRR8084721 | RJ | 11 | 12 | 11 | 7 | 11 | 7 | 9 |
| Hcaps00284_SRR8084716 | RJ | 12 | 13 | 12 | 9 | 13 | 9 | 10 |
| Hcaps00276_SRR8084708 | RJ | 13 | 14 | 13 | 10 | 14 | 10 | 11 |
| Hcaps00287_SRR8084719 | RJ | 14 | 11 | 10 | 8 | 12 | 8 | 8 |
| Hcaps00192_SRR31893281 | Amazon-2 | 15 | 16 | 8 | 22 | 15 | 13 | 12 |
| Hcaps00189_SRR31893278 | Amazon-2 | 16 | 17 | 9 | 23 | 16 | 12 | 13 |
| Hcaps00281_SRR8084713 | NEB | 17 | 15 | 7 | 11 | 17 | 11 | 7 |
| Hcaps00164_SRR31893252 | Sura | 18 | 18 | 6 | 3 | 20 | 3 | 3 |
| Hcaps00167_SRR31893255 | Sura | 19 | 19 | 5 | 2 | 18 | 1 | 2 |
| Hcaps00176_SRR31893264 | Sura | 20 | 20 | 4 | 1 | 19 | 2 | 1 |
| Hcaps00007_SRR17307786 | Sura | 21 | 21 | 2 | 6 | 23 | 5 | 6 |
| Hcaps00026_SRR21059734 | Sura | 22 | 22 | 1 | 5 | 22 | 4 | 5 |
| Hcaps00395_SRR29905415 | Farciminosum | 23 | 23 | 3 | 4 | 21 | 6 | 4 |
| Hcaps00255_SRR6243646 | 27-14 | 24 | 24 | 24 | 24 | 24 | 24 | 24 |
| Hcaps00253_SRR6243644 | Africa | 25 | 30 | 26 | 26 | 26 | 25 | 25 |
| Hcaps00248_SRR6243639 | Africa | 26 | 29 | 25 | 31 | 31 | 31 | 31 |
| Hcapsulatum_GCA017310615 | Africa | 27 | 31 | 27 | 25 | 25 | 26 | 26 |
| Hcaps00378_SRR29905398 | Africa | 28 | 27 | 28 | 27 | 27 | 27 | 27 |
| Hcaps00405_SRR29905425 | Africa | 29 | 28 | 29 | 28 | 28 | 28 | 28 |
| Hcaps00393_SRR29905413 | Africa | 30 | 25 | 31 | 30 | 30 | 30 | 30 |
| Hcaps00390_SRR29905410 | Africa | 31 | 26 | 30 | 29 | 29 | 29 | 29 |
| Hcaps00376_SRR29905396 | Africa | 32 | 32 | 32 | 32 | 32 | 32 | 32 |
| Hcaps00386_SRR29905406 | Africa | 33 | 33 | 33 | 33 | 33 | 33 | 33 |
| Hcapsulatum_GCA017607465 | Capsulatum | 34 | 34 | 34 | 35 | 35 | 35 | 35 |
| Hcapsulatum_GCA017355575 | Capsulatum | 35 | 35 | 35 | 34 | 34 | 34 | 34 |
| Hcaps00187_SRR31893276 | Capsulatum | 36 | 36 | 38 | 37 | 36 | 37 | 38 |
| Hcaps00182_SRR31893271 | Capsulatum | 37 | 37 | 37 | 38 | 38 | 39 | 39 |
| Hcaps00261_SRR6243652 | Capsulatum | 38 | 38 | 36 | 36 | 39 | 36 | 36 |
| Hcaps00236_SRR32121397 | Capsulatum | 39 | 39 | 39 | 39 | 37 | 38 | 37 |
| Hcaps00239_SRR32121400 | Capsulatum | 40 | 40 | 40 | 40 | 40 | 40 | 40 |
| Hcaps00162_SRR31893250 | LAmB | 41 | 41 | 41 | 41 | 41 | 41 | 41 |
| Hcaps00157_SRR31751699 | Ohiense | 42 | 50 | 45 | 43 | 42 | 42 | 42 |
| Hcapsulatum_GCA017607445 | Ohiense | 43 | 49 | 50 | 42 | 43 | 43 | 43 |
| Hcaps00065_SRR21059773 | Ohiense | 44 | 45 | 47 | 46 | 46 | 46 | 46 |
| Hcaps00031_SRR21059739 | Ohiense | 45 | 44 | 46 | 48 | 48 | 48 | 48 |
| Hcaps00051_SRR21059759 | Ohiense | 46 | 42 | 43 | 45 | 45 | 45 | 45 |

| | | | | | | | | |
|---|---|---|---|---|---|---|---|---|
| Hcaps00080_SRR25580018 | Ohiense | 47 | 43 | 42 | 44 | 44 | 44 | 44 |
| Hcaps00030_SRR21059738 | Ohiense | 48 | 47 | 44 | 49 | 49 | 49 | 49 |
| Hcaps00017_SRR17307796 | Ohiense | 49 | 48 | 49 | 50 | 50 | 50 | 50 |
| Hcaps00044_SRR21059752 | Ohiense | 50 | 46 | 48 | 47 | 47 | 47 | 47 |
| Hcaps00271_SRR6243662 | Mississippiense | 51 | 53 | 53 | 51 | 51 | 51 | 52 |
| Hcapsulatum_GCA017310585 | Mississippiense | 52 | 55 | 55 | 52 | 52 | 52 | 51 |
| Hcaps00079_SRR25580017 | Mississippiense | 53 | 52 | 52 | 54 | 54 | 54 | 54 |
| Hcaps00041_SRR21059749 | Mississippiense | 54 | 54 | 54 | 55 | 55 | 55 | 55 |
| Hcaps00084_SRR25580022 | Mississippiense | 55 | 51 | 51 | 53 | 53 | 53 | 53 |
| Hcaps00142_SRR27481868 | India | 56 | 60 | 60 | 60 | 60 | 60 | 60 |
| Hcaps00151_SRR27481877 | India | 57 | 58 | 59 | 59 | 56 | 58 | 58 |
| Hcaps00143_SRR27481869 | India | 58 | 59 | 58 | 58 | 57 | 59 | 59 |
| Hcaps00158_SRR31797080 | India | 59 | 56 | 56 | 56 | 59 | 57 | 57 |
| Hcaps00152_SRR27481878 | India | 60 | 57 | 57 | 57 | 58 | 56 | 56 |

**Positions of genomes in trees shown in Figure S4.**

| Node_names | Lineage | ParSNP | wg0300 | wg0500 | ann0300 | ann0500 | fan0300 | fan0500 |
|---|---|---|---|---|---|---|---|---|
| Hcaps00311_SRR33400187 | Ohiense | 1 | 9 | 37 | 67 | 44 | 40 | 11 |
| Hcaps00321_SRR33400197 | Ohiense | 2 | 8 | 38 | 69 | 46 | 39 | 10 |
| Hcaps00309_SRR33400185 | Ohiense | 3 | 11 | 42 | 64 | 5 | 43 | 41 |
| Hcaps00299_SRR33380106 | Ohiense | 4 | 12 | 39 | 65 | 42 | 38 | 20 |
| Hcaps00296_SRR33380103 | Ohiense | 5 | 7 | 33 | 60 | 40 | 37 | 21 |
| Hcaps00303_SRR33380110 | Ohiense | 6 | 6 | 35 | 62 | 43 | 42 | 37 |
| Hcaps00302_SRR33380109 | Ohiense | 7 | 5 | 36 | 61 | 39 | 44 | 19 |
| Hcaps00297_SRR33380104 | Ohiense | 8 | 87 | 84 | 83 | 75 | 91 | 91 |
| Hcapsulatum_GCA017607445 | Ohiense | 9 | 92 | 93 | 93 | 93 | 93 | 93 |
| Hcaps00003_SRR13433422 | Ohiense | 10 | 74 | 73 | 84 | 83 | 82 | 82 |
| Hcaps00308_SRR33400184 | Ohiense | 11 | 3 | 40 | 68 | 45 | 34 | 12 |
| Hcaps00310_SRR33400186 | Ohiense | 12 | 2 | 44 | 70 | 14 | 36 | 13 |
| Hcaps00306_SRR33400182 | Ohiense | 13 | 1 | 43 | 71 | 13 | 33 | 14 |
| Hcaps00307_SRR33400183 | Ohiense | 14 | 10 | 41 | 63 | 47 | 41 | 36 |
| Hcaps00298_SRR33380105 | Ohiense | 15 | 4 | 34 | 59 | 41 | 35 | 18 |
| Hcaps00070_SRR24516105 | Ohiense | 16 | 76 | 77 | 85 | 85 | 76 | 74 |
| Hcaps00066_SRR24516101 | Ohiense | 17 | 80 | 80 | 89 | 88 | 75 | 76 |
| Hcaps00071_SRR24516106 | Ohiense | 18 | 77 | 75 | 87 | 89 | 77 | 77 |
| Hcaps00068_SRR24516103 | Ohiense | 19 | 75 | 76 | 86 | 86 | 74 | 75 |
| Hcaps00345_SRR33400221 | Ohiense | 20 | 73 | 81 | 37 | 84 | 51 | 69 |
| Hcaps00069_SRR24516104 | Ohiense | 21 | 78 | 74 | 91 | 90 | 78 | 78 |
| Hcaps00366_SRR33400242 | Ohiense | 22 | 46 | 8 | 29 | 48 | 14 | 26 |
| Hcaps00365_SRR33400241 | Ohiense | 23 | 32 | 28 | 20 | 62 | 16 | 68 |
| Hcaps00368_SRR33400244 | Ohiense | 24 | 43 | 9 | 4 | 27 | 8 | 31 |
| Hcaps00353_SRR33400229 | Ohiense | 25 | 33 | 30 | 18 | 24 | 13 | 17 |
| Hcaps00370_SRR33400246 | Ohiense | 26 | 30 | 10 | 9 | 26 | 29 | 34 |
| Hcaps00367_SRR33400243 | Ohiense | 27 | 47 | 7 | 5 | 28 | 7 | 32 |
| Hcaps00354_SRR33400230 | Ohiense | 28 | 38 | 13 | 12 | 70 | 4 | 48 |
| Hcaps00355_SRR33400231 | Ohiense | 29 | 39 | 4 | 11 | 68 | 5 | 44 |
| Hcaps00369_SRR33400245 | Ohiense | 30 | 44 | 3 | 27 | 36 | 6 | 29 |
| Hcaps00371_SRR33400247 | Ohiense | 31 | 48 | 5 | 2 | 7 | 9 | 33 |
| Hcaps00351_SRR33400227 | Ohiense | 32 | 29 | 12 | 6 | 72 | 30 | 25 |
| Hcaps00352_SRR33400228 | Ohiense | 33 | 34 | 29 | 21 | 63 | 18 | 24 |
| Hcaps00322_SRR33400198 | Ohiense | 34 | 14 | 45 | 72 | 22 | 45 | 45 |
| Hcaps00316_SRR33400192 | Ohiense | 35 | 90 | 85 | 82 | 74 | 83 | 87 |

| | | | | | | | | |
|---|---|---|---|---|---|---|---|---|
| Hcaps00314_SRR33400190 | Ohiense | 36 | 51 | 59 | 46 | 19 | 67 | 50 |
| Hcaps00323_SRR33400199 | Ohiense | 37 | 62 | 65 | 44 | 60 | 66 | 66 |
| Hcaps00347_SRR33400223 | Ohiense | 38 | 83 | 91 | 74 | 76 | 87 | 90 |
| Hcaps00270_SRR6243661 | Ohiense | 39 | 82 | 82 | 92 | 92 | 81 | 81 |
| Hcaps00328_SRR33400204 | Ohiense | 40 | 81 | 78 | 88 | 87 | 80 | 79 |
| Hcaps00291_SRR8426586 | Ohiense | 41 | 93 | 94 | 94 | 94 | 94 | 94 |
| Hcaps00155_SRR31751697 | Ohiense | 42 | 18 | 16 | 13 | 69 | 1 | 47 |
| Hcaps00340_SRR33400216 | Ohiense | 43 | 61 | 62 | 39 | 65 | 64 | 6 |
| Hcaps00305_SRR33400181 | Ohiense | 44 | 15 | 46 | 73 | 1 | 49 | 54 |
| Hcaps00356_SRR33400232 | Ohiense | 45 | 22 | 19 | 17 | 67 | 15 | 22 |
| Hcapsulatum_GCA000170615 | Ohiense | 46 | 95 | 95 | 95 | 95 | 95 | 95 |
| Hcaps00362_SRR33400238 | Ohiense | 47 | 91 | 86 | 80 | 73 | 84 | 88 |
| Hcaps00344_SRR33400220 | Ohiense | 48 | 88 | 89 | 77 | 77 | 90 | 89 |
| Hcaps00359_SRR33400235 | Ohiense | 49 | 54 | 52 | 49 | 17 | 54 | 58 |
| Hcaps00357_SRR33400233 | Ohiense | 50 | 53 | 54 | 50 | 16 | 55 | 60 |
| Hcaps00361_SRR33400237 | Ohiense | 51 | 49 | 63 | 55 | 59 | 57 | 63 |
| Hcaps00364_SRR33400240 | Ohiense | 52 | 55 | 53 | 52 | 20 | 56 | 42 |
| Hcaps00358_SRR33400234 | Ohiense | 53 | 36 | 51 | 51 | 18 | 52 | 59 |
| Hcaps00342_SRR33400218 | Ohiense | 54 | 66 | 49 | 48 | 3 | 48 | 56 |
| Hcaps00332_SRR33400208 | Ohiense | 55 | 68 | 57 | 53 | 21 | 21 | 3 |
| Hcaps00156_SRR31751698 | Ohiense | 56 | 94 | 92 | 76 | 81 | 89 | 85 |
| Hcaps00319_SRR33400195 | Ohiense | 57 | 70 | 50 | 56 | 50 | 62 | 61 |
| Hcaps00363_SRR33400239 | Ohiense | 58 | 27 | 23 | 10 | 35 | 11 | 15 |
| Hcaps00318_SRR33400194 | Ohiense | 59 | 72 | 72 | 36 | 55 | 60 | 72 |
| Hcaps00341_SRR33400217 | Ohiense | 60 | 25 | 18 | 1 | 2 | 50 | 55 |
| Hcaps00313_SRR33400189 | Ohiense | 61 | 56 | 67 | 45 | 61 | 17 | 52 |
| Hcaps00333_SRR33400209 | Ohiense | 62 | 89 | 88 | 81 | 80 | 85 | 83 |
| Hcaps00315_SRR33400191 | Ohiense | 63 | 69 | 60 | 34 | 9 | 73 | 53 |
| Hcaps00338_SRR33400214 | Ohiense | 64 | 58 | 69 | 31 | 52 | 69 | 71 |
| Hcaps00337_SRR33400213 | Ohiense | 65 | 42 | 24 | 19 | 64 | 12 | 16 |
| Hcaps00335_SRR33400211 | Ohiense | 66 | 21 | 25 | 15 | 29 | 23 | 7 |
| Hcaps00324_SRR33400200 | Ohiense | 67 | 63 | 66 | 41 | 15 | 65 | 65 |
| Hcaps00360_SRR33400236 | Ohiense | 68 | 50 | 64 | 54 | 58 | 58 | 64 |
| Hcaps00348_SRR33400224 | Ohiense | 69 | 85 | 90 | 78 | 78 | 92 | 92 |
| Hcaps00334_SRR33400210 | Ohiense | 70 | 28 | 15 | 7 | 71 | 31 | 40 |
| Hcaps00346_SRR33400222 | Ohiense | 71 | 35 | 2 | 30 | 51 | 59 | 62 |
| Hcaps00325_SRR33400201 | Ohiense | 72 | 60 | 71 | 35 | 57 | 61 | 73 |
| Hcaps00153_SRR31751695 | Ohiense | 73 | 20 | 32 | 25 | 12 | 32 | 35 |
| Hcaps00343_SRR33400219 | Ohiense | 74 | 67 | 55 | 40 | 11 | 20 | 1 |
| Hcaps00330_SRR33400206 | Ohiense | 75 | 52 | 58 | 47 | 4 | 47 | 51 |
| Hcaps00336_SRR33400212 | Ohiense | 76 | 23 | 27 | 23 | 38 | 22 | 8 |
| Hcaps00326_SRR33400202 | Ohiense | 77 | 64 | 68 | 43 | 53 | 68 | 67 |
| Hcaps00301_SRR33380108 | Ohiense | 78 | 31 | 26 | 8 | 37 | 27 | 9 |
| Hcaps00373_SRR33400249 | Ohiense | 79 | 37 | 1 | 3 | 32 | 53 | 30 |
| Hcaps00312_SRR33400188 | Ohiense | 80 | 65 | 56 | 42 | 10 | 70 | 57 |
| Hcaps00372_SRR33400248 | Ohiense | 81 | 17 | 11 | 16 | 30 | 26 | 28 |
| Hcaps00304_SRR33400180 | Ohiense | 82 | 13 | 47 | 66 | 23 | 46 | 38 |
| Hcaps00350_SRR33400226 | Ohiense | 83 | 16 | 22 | 14 | 31 | 3 | 43 |
| Hcaps00320_SRR33400196 | Ohiense | 84 | 45 | 6 | 24 | 6 | 10 | 39 |
| Hcaps00349_SRR33400225 | Ohiense | 85 | 26 | 17 | 32 | 54 | 2 | 46 |
| Hcaps00331_SRR33400207 | Ohiense | 86 | 41 | 21 | 28 | 33 | 24 | 2 |
| Hcaps00295_SRR33380102 | Ohiense | 87 | 84 | 83 | 79 | 82 | 86 | 86 |

| | | | | | | | | |
|---|---|---|---|---|---|---|---|---|
| Hcaps00329_SRR33400205 | Ohiense | 88 | 40 | 20 | 26 | 25 | 25 | 5 |
| Hcaps00154_SRR31751696 | Ohiense | 89 | 19 | 31 | 22 | 49 | 19 | 27 |
| Hcaps00327_SRR33400203 | Ohiense | 90 | 59 | 61 | 33 | 8 | 72 | 49 |
| Hcaps00317_SRR33400193 | Ohiense | 91 | 71 | 70 | 38 | 56 | 71 | 70 |
| Hcaps00339_SRR33400215 | Ohiense | 92 | 57 | 48 | 57 | 34 | 63 | 4 |
| Hcaps00300_SRR33380107 | Ohiense | 93 | 24 | 14 | 58 | 66 | 28 | 23 |
| Hcaps00157_SRR31751699 | Ohiense | 94 | 86 | 87 | 75 | 79 | 88 | 84 |
| Hcaps00067_SRR24516102 | Ohiense | 95 | 79 | 79 | 90 | 91 | 79 | 80 |
| Hcaps00005_SRR13508696 | Mississippiense | 96 | 106 | 108 | 107 | 106 | 106 | 106 |
| Hcapsulatum_GCA017310585 | Mississippiense | 97 | 108 | 107 | 108 | 108 | 108 | 108 |
| Hcaps00249_SRR6243640 | Mississippiense | 98 | 107 | 106 | 106 | 107 | 107 | 107 |
| Hcapsulatum_GCA017355575 | Capsu | 99 | 101 | 100 | 101 | 101 | 101 | 101 |
| Hcaps00006_SRR13508697 | Capsu | 100 | 97 | 97 | 97 | 97 | 96 | 96 |
| Hcapsulatum_GCA000150115 | Capsu | 101 | 102 | 102 | 102 | 102 | 102 | 102 |
| Hcapsulatum_GCA013420885 | Capsu | 102 | 98 | 101 | 100 | 100 | 100 | 100 |
| Hcaps00259_SRR6243650 | Capsu | 103 | 99 | 96 | 96 | 96 | 97 | 97 |
| Hcaps00004_SRR13433424 | Capsu | 104 | 96 | 98 | 98 | 98 | 98 | 98 |
| Hcapsulatum_GCA017607465 | Capsu | 105 | 100 | 99 | 99 | 99 | 99 | 99 |
| Hcaps00002_SRR13433402 | Africa | 106 | 104 | 104 | 103 | 103 | 104 | 104 |
| Hcapsulatum_GCA017310615 | Africa | 107 | 103 | 103 | 104 | 104 | 103 | 103 |
| Hcapsulatum_GCA000151005 | Africa | 108 | 105 | 105 | 105 | 105 | 105 | 105 |
| Hcaps00141_SRR27481867 | India (root) | 109 | 109 | 109 | 109 | 109 | 109 | 109 |

**Positions of genomes in trees shown in Figures 3 and S5-S7.**

| **Node_names** | **Lineage** | **wg0300** | **wg0500** | **ann0300** | **ann0500** | **fan0300** | **fan0500** |
|---|---|---|---|---|---|---|---|
| Hcaps00164_SRR31893252 | Sura | 1 | 2 | 2 | 69 | 2 | 69 |
| Hcaps00166_SRR31893254 | Sura | 2 | 1 | 1 | 68 | 1 | 68 |
| Hcaps00170_SRR31893258 | Sura | 3 | 3 | 3 | 70 | 3 | 70 |
| Hcaps00290_SRR8347492 | Sura | 4 | 5 | 5 | 72 | 5 | 72 |
| Hcaps00172_SRR31893260 | Sura | 5 | 4 | 4 | 71 | 4 | 71 |
| Hcaps00173_SRR31893261 | Sura | 6 | 6 | 6 | 73 | 6 | 73 |
| Hcaps00163_SRR31893251 | Sura | 7 | 7 | 7 | 74 | 7 | 74 |
| Hcaps00168_SRR31893256 | Sura | 8 | 8 | 9 | 75 | 9 | 75 |
| Hcaps00165_SRR31893253 | Sura | 9 | 9 | 8 | 76 | 8 | 76 |
| Hcaps00167_SRR31893255 | Sura | 10 | 10 | 10 | 77 | 10 | 77 |
| Hcaps00120_SRR25580058 | Sura | 11 | 11 | 11 | 78 | 11 | 78 |
| Hcaps00020_SRR21059728 | Sura | 12 | 12 | 12 | 79 | 12 | 79 |
| Hcaps00135_SRR25636145 | Sura | 13 | 13 | 13 | 80 | 13 | 80 |
| Hcaps00014_SRR17307793 | Sura | 14 | 14 | 14 | 81 | 14 | 81 |
| Hcaps00177_SRR31893265 | Sura | 15 | 16 | 16 | 82 | 15 | 83 |
| Hcaps00176_SRR31893264 | Sura | 16 | 15 | 15 | 83 | 16 | 82 |
| Hcaps00254_SRR6243645 | Sura | 17 | 17 | 17 | 84 | 17 | 84 |
| Hcaps00178_SRR31893266 | Sura | 18 | 18 | 18 | 85 | 18 | 85 |
| Hcaps00175_SRR31893263 | Sura | 19 | 19 | 19 | 86 | 19 | 86 |
| Hcaps00252_SRR6243643 | Sura | 20 | 20 | 20 | 87 | 20 | 87 |
| Hcaps00110_SRR25580048 | Sura | 21 | 21 | 21 | 88 | 21 | 88 |
| Hcaps00007_SRR17307786 | Sura | 22 | 23 | 23 | 90 | 23 | 90 |
| Hcaps00174_SRR31893262 | Sura | 23 | 22 | 22 | 89 | 22 | 89 |
| Hcaps00015_SRR17307794 | Sura | 24 | 24 | 24 | 91 | 24 | 91 |
| Hcaps00026_SRR21059734 | Sura | 25 | 25 | 25 | 92 | 25 | 92 |
| Hcaps00013_SRR17307792 | Sura | 26 | 26 | 26 | 93 | 26 | 93 |
| Hcaps00008_SRR17307787 | Sura | 27 | 27 | 27 | 94 | 27 | 94 |

| | | | | | | | |
|---|---|---|---|---|---|---|---|
| Hcaps00406_SRR29905426 | Farciminosum | 28 | 28 | 29 | 95 | 28 | 95 |
| Hcaps00395_SRR29905415 | Farciminosum | 29 | 29 | 28 | 96 | 29 | 96 |
| Hcaps00382_SRR29905402 | Farciminosum | 30 | 30 | 30 | 97 | 30 | 97 |
| Hcaps00105_SRR25580043 | Sura/Farci | 31 | 31 | 31 | 98 | 31 | 98 |
| Hcaps00001_ERR7414368 | Sura/Farci | 32 | 33 | 33 | 100 | 33 | 100 |
| Hcapsulatum_GCA000313325 | Sura/Farci | 33 | 32 | 32 | 99 | 32 | 99 |
| Hcaps00275_SRR8084707 | RJ | 34 | 34 | 36 | 48 | 36 | 48 |
| Hcaps00276_SRR8084708 | RJ | 35 | 35 | 35 | 49 | 35 | 49 |
| Hcaps00277_SRR8084709 | RJ | 36 | 36 | 37 | 50 | 37 | 51 |
| Hcaps00273_SRR8084705 | RJ | 37 | 37 | 34 | 51 | 34 | 50 |
| Hcaps00274_SRR8084706 | RJ | 38 | 38 | 38 | 52 | 38 | 52 |
| Hcaps00272_SRR8084704 | RJ | 39 | 39 | 39 | 53 | 39 | 53 |
| Hcaps00282_SRR8084714 | RJ | 40 | 41 | 40 | 54 | 43 | 54 |
| Hcaps00284_SRR8084716 | RJ | 41 | 40 | 41 | 55 | 44 | 55 |
| Hcaps00285_SRR8084717 | RJ | 42 | 42 | 42 | 56 | 41 | 56 |
| Hcaps00289_SRR8084721 | RJ | 43 | 43 | 43 | 57 | 40 | 57 |
| Hcaps00283_SRR8084715 | RJ | 44 | 44 | 44 | 58 | 42 | 58 |
| Hcaps00288_SRR8084720 | RJ | 45 | 45 | 45 | 59 | 45 | 59 |
| Hcaps00287_SRR8084719 | RJ | 46 | 46 | 46 | 60 | 46 | 60 |
| Hcaps00278_SRR8084710 | RJ | 47 | 47 | 47 | 61 | 47 | 61 |
| Hcaps00193_SRR31893282 | Amazon-2 | 48 | 50 | 50 | 64 | 48 | 64 |
| Hcaps00192_SRR31893281 | Amazon-2 | 49 | 51 | 52 | 65 | 50 | 66 |
| Hcaps00189_SRR31893278 | Amazon-2 | 50 | 52 | 51 | 66 | 49 | 65 |
| Hcaps00191_SRR31893280 | Amazon-2 | 51 | 53 | 53 | 67 | 51 | 67 |
| Hcaps00280_SRR8084712 | NEB | 52 | 49 | 48 | 62 | 52 | 62 |
| Hcaps00281_SRR8084713 | NEB | 53 | 48 | 49 | 63 | 53 | 63 |
| Hcaps00210_SRR31893300 | Mz5-like | 54 | 54 | 54 | 2 | 63 | 11 |
| Hcaps00196_SRR31893286 | Mz5-like | 55 | 55 | 55 | 1 | 64 | 12 |
| Hcaps00180_SRR31893268 | Mz5-like | 56 | 56 | 57 | 3 | 66 | 13 |
| Hcaps00199_SRR31893289 | Mz5-like | 57 | 57 | 56 | 4 | 65 | 14 |
| Hcaps00169_SRR31893257 | Mz5-like | 58 | 63 | 63 | 12 | 60 | 7 |
| Hcaps00203_SRR31893293 | Mz5-like | 59 | 61 | 62 | 9 | 70 | 18 |
| Hcaps00198_SRR31893288 | Mz5-like | 60 | 59 | 60 | 6 | 69 | 17 |
| Hcaps00201_SRR31893291 | Mz5-like | 61 | 60 | 59 | 7 | 68 | 16 |
| Hcaps00222_SRR31893312 | Mz5-like | 62 | 62 | 61 | 8 | 71 | 19 |
| Hcaps00212_SRR31893302 | Mz5-like | 63 | 58 | 58 | 5 | 67 | 15 |
| Hcaps00218_SRR31893308 | Mz5-like | 64 | 67 | 64 | 14 | 54 | 2 |
| Hcaps00208_SRR31893298 | Mz5-like | 65 | 66 | 65 | 13 | 55 | 1 |
| Hcaps00217_SRR31893307 | Mz5-like | 66 | 68 | 66 | 15 | 56 | 3 |
| Hcaps00223_SRR31893313 | Mz5-like | 67 | 69 | 68 | 18 | 59 | 6 |
| Hcaps00204_SRR31893294 | Mz5-like | 68 | 70 | 69 | 17 | 58 | 5 |
| Hcaps00161_SRR31893249 | Mz5-like | 69 | 71 | 67 | 16 | 57 | 4 |
| Hcaps00207_SRR31893297 | Mz5-like | 70 | 65 | 71 | 10 | 61 | 9 |
| Hcaps00200_SRR31893290 | Mz5-like | 71 | 64 | 70 | 11 | 62 | 8 |
| Hcaps00190_SRR31893279 | Mz5-like | 72 | 72 | 72 | 19 | 72 | 10 |
| Hcaps00219_SRR31893309 | Mz5-like | 73 | 75 | 74 | 20 | 74 | 21 |
| Hcaps00211_SRR31893301 | Mz5-like | 74 | 76 | 73 | 21 | 73 | 20 |
| Hcaps00221_SRR31893311 | Mz5-like | 75 | 73 | 75 | 22 | 75 | 22 |
| Hcaps00215_SRR31893305 | Mz5-like | 76 | 74 | 76 | 24 | 76 | 24 |
| Hcaps00213_SRR31893303 | Mz5-like | 77 | 78 | 77 | 25 | 78 | 26 |
| Hcaps00202_SRR31893292 | Mz5-like | 78 | 77 | 78 | 26 | 77 | 25 |
| Hcaps00209_SRR31893299 | Mz5-like | 79 | 79 | 79 | 23 | 79 | 23 |

| | | | | | | | |
|---|---|---|---|---|---|---|---|
| Hcaps00160_SRR31893248 | Mz5-like | 80 | 80 | 80 | 27 | 80 | 27 |
| Hcaps00206_SRR31893296 | Mz5-like | 81 | 82 | 81 | 30 | 81 | 29 |
| Hcaps00214_SRR31893304 | Mz5-like | 82 | 83 | 82 | 29 | 82 | 30 |
| Hcaps00159_SRR31893247 | Mz5-like | 83 | 81 | 83 | 28 | 83 | 28 |
| Hcaps00228_SRR32121389 | Mz5-like | 84 | 85 | 84 | 32 | 85 | 31 |
| Hcaps00227_SRR32121388 | Mz5-like | 85 | 84 | 85 | 31 | 84 | 32 |
| Hcaps00241_SRR32121402 | Mz5-like | 86 | 87 | 87 | 34 | 86 | 34 |
| Hcaps00240_SRR32121401 | Mz5-like | 87 | 86 | 86 | 33 | 87 | 33 |
| Hcaps00226_SRR32121387 | Mz5-like | 88 | 88 | 88 | 35 | 88 | 35 |
| Hcaps00230_SRR32121391 | Mz5-like | 89 | 89 | 89 | 36 | 89 | 36 |
| Hcaps00220_SRR31893310 | Mz5-like | 90 | 90 | 90 | 38 | 91 | 38 |
| Hcaps00231_SRR32121392 | Mz5-like | 91 | 91 | 91 | 37 | 90 | 37 |
| Hcaps00205_SRR31893295 | Mz5-like | 92 | 92 | 92 | 39 | 92 | 39 |
| Hcaps00229_SRR32121390 | Mz5-like | 93 | 93 | 93 | 40 | 93 | 40 |
| Hcaps00258_SRR6243649 | Mz5-like | 94 | 94 | 95 | 42 | 94 | 41 |
| Hcaps00374_SRR33700783 | Mz5-like | 95 | 95 | 94 | 41 | 95 | 42 |
| Hcaps00042_SRR21059750 | Mz5-like | 96 | 96 | 96 | 43 | 96 | 43 |
| Hcaps00195_SRR31893285 | Amazon-1 | 97 | 98 | 98 | 45 | 98 | 45 |
| Hcaps00194_SRR31893284 | Amazon-1 | 98 | 97 | 97 | 44 | 97 | 44 |
| Hcaps00294_SRR31893283 | Amazon-1 | 99 | 99 | 99 | 47 | 99 | 47 |
| Hcaps00197_SRR31893287 | Amazon-1 | 100 | 100 | 100 | 46 | 100 | 46 |
| Hcaps00255_SRR6243646 | 27-14 | 101 | 101 | 101 | 101 | 101 | 101 |
| Hcaps00279_SRR8084711 | LAmB | 102 | 102 | 159 | 102 | 102 | 102 |
| Hcaps00162_SRR31893250 | LAmB | 103 | 103 | 160 | 103 | 103 | 103 |
| Hcaps00286_SRR8084718 | LAmB | 104 | 104 | 161 | 104 | 104 | 104 |
| Hcaps00400_SRR29905420 | Africa | 105 | 105 | 102 | 117 | 106 | 117 |
| Hcaps00402_SRR29905422 | Africa | 106 | 106 | 103 | 116 | 105 | 116 |
| Hcaps00391_SRR29905411 | Africa | 107 | 107 | 104 | 118 | 107 | 118 |
| Hcaps00385_SRR29905405 | Africa | 108 | 108 | 105 | 119 | 108 | 119 |
| Hcaps00392_SRR29905412 | Africa | 109 | 109 | 106 | 120 | 109 | 120 |
| Hcaps00253_SRR6243644 | Africa | 110 | 111 | 108 | 113 | 111 | 114 |
| Hcapsulatum_GCA017310615 | Africa | 111 | 110 | 107 | 114 | 110 | 113 |
| Hcaps00248_SRR6243639 | Africa | 112 | 112 | 109 | 115 | 112 | 115 |
| Hcaps00396_SRR29905416 | Africa | 113 | 113 | 111 | 122 | 114 | 122 |
| Hcaps00393_SRR29905413 | Africa | 114 | 114 | 110 | 121 | 113 | 121 |
| Hcaps00375_SRR29905395 | Africa | 115 | 115 | 112 | 123 | 115 | 123 |
| Hcaps00379_SRR29905399 | Africa | 116 | 116 | 113 | 124 | 116 | 124 |
| Hcaps00387_SRR29905407 | Africa | 117 | 117 | 114 | 125 | 117 | 125 |
| Hcaps00390_SRR29905410 | Africa | 118 | 118 | 115 | 126 | 118 | 126 |
| Hcaps00397_SRR29905417 | Africa | 119 | 119 | 116 | 127 | 119 | 127 |
| Hcaps00404_SRR29905424 | Africa | 120 | 120 | 117 | 105 | 120 | 105 |
| Hcaps00405_SRR29905425 | Africa | 121 | 121 | 118 | 106 | 121 | 106 |
| Hcaps00381_SRR29905401 | Africa | 122 | 122 | 119 | 107 | 122 | 107 |
| Hcaps00384_SRR29905404 | Africa | 123 | 123 | 120 | 108 | 123 | 108 |
| Hcaps00377_SRR29905397 | Africa | 124 | 124 | 121 | 109 | 124 | 109 |
| Hcaps00378_SRR29905398 | Africa | 125 | 126 | 123 | 111 | 126 | 111 |
| Hcaps00398_SRR29905418 | Africa | 126 | 125 | 122 | 110 | 125 | 110 |
| Hcapsulatum_GCA000151035 | Africa | 127 | 127 | 124 | 112 | 127 | 112 |
| Hcaps00389_SRR29905409 | Africa | 128 | 128 | 125 | 128 | 128 | 128 |
| Hcaps00383_SRR29905403 | Africa | 129 | 129 | 126 | 129 | 129 | 129 |
| Hcaps00403_SRR29905423 | Africa | 130 | 130 | 127 | 130 | 131 | 131 |
| Hcaps00376_SRR29905396 | Africa | 131 | 131 | 128 | 131 | 130 | 130 |

| | | | | | | | |
|---|---|---|---|---|---|---|---|
| Hcaps00394_SRR29905414 | Africa | 132 | 133 | 129 | 132 | 132 | 132 |
| Hcaps00399_SRR29905419 | Africa | 133 | 132 | 130 | 133 | 133 | 133 |
| Hcaps00401_SRR29905421 | Africa | 134 | 134 | 131 | 134 | 134 | 134 |
| Hcaps00380_SRR29905400 | Africa | 135 | 135 | 132 | 135 | 135 | 135 |
| Hcaps00386_SRR29905406 | Africa | 136 | 136 | 133 | 136 | 136 | 136 |
| Hcaps00182_SRR31893271 | Capsu | 137 | 138 | 134 | 142 | 141 | 142 |
| Hcaps00183_SRR31893272 | Capsu | 138 | 137 | 135 | 141 | 142 | 141 |
| Hcaps00185_SRR31893274 | Capsu | 139 | 139 | 136 | 143 | 143 | 143 |
| Hcaps00179_SRR31893267 | Capsu | 140 | 140 | 141 | 146 | 145 | 146 |
| Hcaps00232_SRR32121393 | Capsu | 141 | 141 | 142 | 145 | 144 | 144 |
| Hcaps00181_SRR31893270 | Capsu | 142 | 142 | 143 | 144 | 146 | 145 |
| Hcaps00186_SRR31893275 | Capsu | 143 | 143 | 137 | 137 | 137 | 138 |
| Hcaps00187_SRR31893276 | Capsu | 144 | 144 | 138 | 138 | 138 | 137 |
| Hcaps00188_SRR31893277 | Capsu | 145 | 145 | 139 | 139 | 139 | 139 |
| Hcaps00293_SRR31893269 | Capsu | 146 | 146 | 140 | 140 | 140 | 140 |
| Hcaps00184_SRR31893273 | Capsu | 147 | 147 | 144 | 147 | 147 | 147 |
| Hcaps00261_SRR6243652 | Capsu | 148 | 148 | 145 | 148 | 148 | 148 |
| Hcaps00259_SRR6243650 | Capsu | 149 | 150 | 147 | 152 | 150 | 150 |
| Hcaps00256_SRR6243647 | Capsu | 150 | 149 | 146 | 151 | 149 | 149 |
| Hcapsulatum_GCA017607465 | Capsu | 151 | 152 | 150 | 155 | 153 | 153 |
| Hcaps00235_SRR32121396 | Capsu | 152 | 153 | 149 | 154 | 152 | 152 |
| Hcaps00234_SRR32121395 | Capsu | 153 | 151 | 148 | 153 | 151 | 151 |
| Hcapsulatum_GCA017355575 | Capsu | 154 | 154 | 151 | 156 | 154 | 154 |
| Hcaps00233_SRR32121394 | Capsu | 155 | 155 | 152 | 157 | 155 | 155 |
| Hcaps00237_SRR32121398 | Capsu | 156 | 156 | 153 | 158 | 156 | 156 |
| Hcaps00236_SRR32121397 | Capsu | 157 | 161 | 155 | 150 | 157 | 161 |
| Hcaps00238_SRR32121399 | Capsu | 158 | 160 | 154 | 149 | 158 | 160 |
| Hcaps00225_SRR32121386 | Capsu | 159 | 157 | 157 | 159 | 159 | 157 |
| Hcaps00224_SRR32121385 | Capsu | 160 | 158 | 156 | 160 | 160 | 158 |
| Hcaps00239_SRR32121400 | Capsu | 161 | 159 | 158 | 161 | 161 | 159 |
| Hcaps00046_SRR21059754 | Ohiense | 162 | 163 | 162 | 163 | 168 | 168 |
| Hcaps00057_SRR21059765 | Ohiense | 163 | 162 | 163 | 162 | 166 | 166 |
| Hcaps00056_SRR21059764 | Ohiense | 164 | 164 | 164 | 164 | 167 | 169 |
| Hcaps00062_SRR21059770 | Ohiense | 165 | 165 | 165 | 165 | 169 | 167 |
| Hcaps00054_SRR21059762 | Ohiense | 166 | 167 | 170 | 170 | 171 | 171 |
| Hcaps00040_SRR21059748 | Ohiense | 167 | 166 | 171 | 171 | 170 | 170 |
| Hcaps00055_SRR21059763 | Ohiense | 168 | 168 | 172 | 172 | 172 | 172 |
| Hcaps00031_SRR21059739 | Ohiense | 169 | 171 | 173 | 174 | 173 | 174 |
| Hcaps00407_SRR29905427 | Ohiense | 170 | 172 | 187 | 178 | 178 | 179 |
| Hcaps00019_SRR21059727 | Ohiense | 171 | 170 | 175 | 175 | 175 | 175 |
| Hcaps00018_SRR21059726 | Ohiense | 172 | 169 | 174 | 173 | 174 | 173 |
| Hcaps00045_SRR21059753 | Ohiense | 173 | 173 | 176 | 176 | 176 | 176 |
| Hcaps00063_SRR21059771 | Ohiense | 174 | 174 | 168 | 166 | 165 | 163 |
| Hcaps00048_SRR21059756 | Ohiense | 175 | 175 | 169 | 167 | 164 | 162 |
| Hcaps00059_SRR21059767 | Ohiense | 176 | 176 | 167 | 169 | 163 | 165 |
| Hcaps00052_SRR21059760 | Ohiense | 177 | 177 | 166 | 168 | 162 | 164 |
| Hcaps00021_SRR21059729 | Ohiense | 178 | 178 | 199 | 189 | 200 | 200 |
| Hcaps00053_SRR21059761 | Ohiense | 179 | 183 | 179 | 188 | 186 | 188 |
| Hcaps00049_SRR21059757 | Ohiense | 180 | 182 | 180 | 187 | 187 | 189 |
| Hcaps00111_SRR25580049 | Ohiense | 181 | 188 | 177 | 191 | 183 | 181 |
| Hcaps00093_SRR25580031 | Ohiense | 182 | 189 | 178 | 192 | 184 | 182 |
| Hcaps00043_SRR21059751 | Ohiense | 183 | 180 | 182 | 184 | 194 | 185 |

| | | | | | | | |
|---|---|---|---|---|---|---|---|
| Hcaps00047_SRR21059755 | Ohiense | 184 | 179 | 181 | 185 | 193 | 186 |
| Hcaps00039_SRR21059747 | Ohiense | 185 | 181 | 183 | 186 | 195 | 187 |
| Hcaps00061_SRR21059769 | Ohiense | 186 | 194 | 184 | 177 | 177 | 177 |
| Hcaps00123_SRR25580061 | Ohiense | 187 | 190 | 198 | 193 | 185 | 180 |
| Hcaps00132_SRR25580070 | Ohiense | 188 | 191 | 196 | 194 | 197 | 198 |
| Hcaps00099_SRR25580037 | Ohiense | 189 | 192 | 197 | 195 | 196 | 197 |
| Hcaps00072_SRR25580010 | Ohiense | 190 | 184 | 188 | 179 | 179 | 178 |
| Hcaps00065_SRR21059773 | Ohiense | 191 | 185 | 200 | 190 | 199 | 199 |
| Hcaps00106_SRR25580044 | Ohiense | 192 | 187 | 189 | 182 | 181 | 194 |
| Hcaps00121_SRR25580059 | Ohiense | 193 | 186 | 190 | 183 | 182 | 195 |
| Hcaps00094_SRR25580032 | Ohiense | 194 | 196 | 191 | 181 | 180 | 183 |
| Hcaps00075_SRR25580013 | Ohiense | 195 | 195 | 186 | 180 | 188 | 184 |
| Hcaps00116_SRR25580054 | Ohiense | 196 | 193 | 185 | 204 | 198 | 205 |
| Hcaps00108_SRR25580046 | Ohiense | 197 | 198 | 194 | 199 | 192 | 192 |
| Hcaps00115_SRR25580053 | Ohiense | 198 | 197 | 195 | 198 | 191 | 193 |
| Hcaps00118_SRR25580056 | Ohiense | 199 | 199 | 192 | 196 | 189 | 190 |
| Hcaps00107_SRR25580045 | Ohiense | 200 | 200 | 193 | 197 | 190 | 191 |
| Hcaps00022_SRR21059730 | Ohiense | 201 | 204 | 201 | 205 | 201 | 202 |
| Hcaps00077_SRR25580015 | Ohiense | 202 | 205 | 207 | 206 | 203 | 201 |
| Hcaps00088_SRR25580026 | Ohiense | 203 | 206 | 206 | 207 | 202 | 203 |
| Hcaps00050_SRR21059758 | Ohiense | 204 | 202 | 203 | 201 | 205 | 207 |
| Hcaps00064_SRR21059772 | Ohiense | 205 | 201 | 202 | 200 | 204 | 196 |
| Hcaps00098_SRR25580036 | Ohiense | 206 | 203 | 204 | 202 | 206 | 206 |
| Hcaps00267_SRR6243658 | Ohiense | 207 | 209 | 205 | 210 | 207 | 210 |
| Hcaps00114_SRR25580052 | Ohiense | 208 | 208 | 209 | 209 | 209 | 209 |
| Hcaps00095_SRR25580033 | Ohiense | 209 | 207 | 208 | 208 | 208 | 208 |
| Hcaps00129_SRR25580067 | Ohiense | 210 | 210 | 210 | 203 | 210 | 204 |
| Hcaps00101_SRR25580039 | Ohiense | 211 | 211 | 229 | 241 | 227 | 214 |
| Hcaps00262_SRR6243653 | Ohiense | 212 | 230 | 227 | 246 | 214 | 228 |
| Hcaps00265_SRR6243656 | Ohiense | 213 | 218 | 238 | 237 | 213 | 221 |
| Hcaps00266_SRR6243657 | Ohiense | 214 | 217 | 216 | 231 | 216 | 218 |
| Hcaps00246_SRR6243637 | Ohiense | 215 | 216 | 215 | 230 | 215 | 217 |
| Hcaps00264_SRR6243655 | Ohiense | 216 | 229 | 235 | 235 | 211 | 219 |
| Hcaps00263_SRR6243654 | Ohiense | 217 | 223 | 231 | 233 | 212 | 222 |
| Hcaps00127_SRR25580065 | Ohiense | 218 | 224 | 230 | 234 | 230 | 216 |
| Hcaps00260_SRR6243651 | Ohiense | 219 | 220 | 212 | 238 | 218 | 212 |
| Hcapsulatum_GCA017607445 | Ohiense | 220 | 221 | 213 | 240 | 220 | 213 |
| Hcaps00269_SRR6243660 | Ohiense | 221 | 219 | 211 | 239 | 219 | 211 |
| Hcaps00130_SRR25580068 | Ohiense | 222 | 238 | 214 | 245 | 221 | 230 |
| Hcaps00128_SRR25580066 | Ohiense | 223 | 222 | 217 | 232 | 228 | 215 |
| Hcaps00117_SRR25580055 | Ohiense | 224 | 227 | 233 | 248 | 234 | 237 |
| Hcaps00103_SRR25580041 | Ohiense | 225 | 225 | 222 | 250 | 233 | 235 |
| Hcaps00060_SRR21059768 | Ohiense | 226 | 228 | 234 | 236 | 236 | 220 |
| Hcaps00036_SRR21059744 | Ohiense | 227 | 226 | 232 | 251 | 235 | 236 |
| Hcaps00133_SRR25580071 | Ohiense | 228 | 212 | 236 | 255 | 237 | 233 |
| Hcaps00100_SRR25580038 | Ohiense | 229 | 213 | 237 | 254 | 238 | 238 |
| Hcaps00268_SRR6243659 | Ohiense | 230 | 231 | 218 | 242 | 217 | 229 |
| Hcaps00038_SRR21059746 | Ohiense | 231 | 235 | 226 | 249 | 224 | 223 |
| Hcaps00037_SRR21059745 | Ohiense | 232 | 233 | 220 | 252 | 231 | 226 |
| Hcaps00051_SRR21059759 | Ohiense | 233 | 237 | 223 | 244 | 223 | 224 |
| Hcaps00035_SRR21059743 | Ohiense | 234 | 236 | 224 | 243 | 222 | 225 |
| Hcaps00096_SRR25580034 | Ohiense | 235 | 232 | 228 | 247 | 229 | 234 |

| | | | | | | | |
|---|---|---|---|---|---|---|---|
| Hcaps00122_SRR25580060 | Ohiense | 236 | 215 | 221 | 256 | 226 | 232 |
| Hcaps00080_SRR25580018 | Ohiense | 237 | 234 | 219 | 253 | 232 | 227 |
| Hcaps00131_SRR25580069 | Ohiense | 238 | 214 | 225 | 257 | 225 | 231 |
| Hcaps00024_SRR21059732 | Ohiense | 239 | 240 | 239 | 214 | 239 | 239 |
| Hcaps00058_SRR21059766 | Ohiense | 240 | 239 | 240 | 215 | 240 | 240 |
| Hcaps00034_SRR21059742 | Ohiense | 241 | 241 | 241 | 216 | 241 | 241 |
| Hcaps00027_SRR21059735 | Ohiense | 242 | 242 | 242 | 218 | 243 | 242 |
| Hcaps00025_SRR21059733 | Ohiense | 243 | 243 | 243 | 217 | 242 | 243 |
| Hcaps00030_SRR21059738 | Ohiense | 244 | 244 | 245 | 211 | 244 | 245 |
| Hcaps00032_SRR21059740 | Ohiense | 245 | 245 | 244 | 212 | 245 | 244 |
| Hcaps00033_SRR21059741 | Ohiense | 246 | 246 | 246 | 213 | 246 | 246 |
| Hcaps00028_SRR21059736 | Ohiense | 247 | 247 | 247 | 219 | 247 | 247 |
| Hcaps00029_SRR21059737 | Ohiense | 248 | 248 | 249 | 221 | 249 | 248 |
| Hcaps00023_SRR21059731 | Ohiense | 249 | 249 | 248 | 220 | 248 | 249 |
| Hcaps00104_SRR25580042 | Ohiense | 250 | 250 | 250 | 222 | 250 | 250 |
| Hcaps00011_SRR17307790 | Ohiense | 251 | 251 | 251 | 223 | 252 | 251 |
| Hcaps00017_SRR17307796 | Ohiense | 252 | 252 | 252 | 224 | 251 | 252 |
| Hcaps00016_SRR17307795 | Ohiense | 253 | 253 | 253 | 225 | 253 | 253 |
| Hcaps00009_SRR17307788 | Ohiense | 254 | 254 | 254 | 227 | 254 | 255 |
| Hcaps00012_SRR17307791 | Ohiense | 255 | 255 | 255 | 226 | 255 | 254 |
| Hcaps00097_SRR25580035 | Ohiense | 256 | 256 | 256 | 229 | 257 | 256 |
| Hcaps00124_SRR25580062 | Ohiense | 257 | 257 | 257 | 228 | 256 | 257 |
| Hcaps00073_SRR25580011 | Ohiense | 258 | 258 | 258 | 259 | 258 | 258 |
| Hcaps00109_SRR25580047 | Ohiense | 259 | 259 | 259 | 258 | 259 | 259 |
| Hcaps00074_SRR25580012 | Ohiense | 260 | 260 | 260 | 260 | 260 | 260 |
| Hcaps00044_SRR21059752 | Ohiense | 261 | 261 | 261 | 261 | 261 | 261 |
| Hcaps00113_SRR25580051 | Ohiense | 262 | 262 | 262 | 262 | 262 | 262 |
| Hcaps00084_SRR25580022 | Mississippiense | 263 | 265 | 272 | 266 | 263 | 266 |
| Hcaps00076_SRR25580014 | Mississippiense | 264 | 266 | 271 | 265 | 274 | 265 |
| Hcaps00089_SRR25580027 | Mississippiense | 265 | 263 | 269 | 264 | 264 | 263 |
| Hcaps00081_SRR25580019 | Mississippiense | 266 | 264 | 270 | 275 | 273 | 275 |
| Hcaps00087_SRR25580025 | Mississippiense | 267 | 267 | 263 | 270 | 267 | 270 |
| Hcaps00082_SRR25580020 | Mississippiense | 268 | 268 | 264 | 269 | 268 | 269 |
| Hcaps00125_SRR25580063 | Mississippiense | 269 | 269 | 265 | 267 | 266 | 267 |
| Hcaps00091_SRR25580029 | Mississippiense | 270 | 270 | 267 | 272 | 269 | 272 |
| Hcaps00092_SRR25580030 | Mississippiense | 271 | 271 | 266 | 271 | 270 | 271 |
| Hcaps00085_SRR25580023 | Mississippiense | 272 | 272 | 268 | 263 | 265 | 264 |
| Hcaps00079_SRR25580017 | Mississippiense | 273 | 273 | 276 | 273 | 271 | 273 |
| Hcaps00078_SRR25580016 | Mississippiense | 274 | 274 | 275 | 274 | 272 | 274 |
| Hcaps00102_SRR25580040 | Mississippiense | 275 | 275 | 277 | 277 | 277 | 277 |
| Hcaps00136_SRR25758805 | Mississippiense | 276 | 276 | 278 | 278 | 278 | 276 |
| Hcaps00090_SRR25580028 | Mississippiense | 277 | 277 | 273 | 268 | 276 | 268 |
| Hcaps00126_SRR25580064 | Mississippiense | 278 | 278 | 274 | 276 | 275 | 278 |
| Hcaps00251_SRR6243642 | Mississippiense | 279 | 280 | 280 | 280 | 279 | 280 |
| Hcaps00134_SRR25636130 | Mississippiense | 280 | 279 | 279 | 279 | 280 | 279 |
| Hcaps00112_SRR25580050 | Mississippiense | 281 | 281 | 281 | 281 | 281 | 281 |
| Hcaps00086_SRR25580024 | Mississippiense | 282 | 282 | 283 | 283 | 283 | 282 |
| Hcaps00010_SRR17307789 | Mississippiense | 283 | 283 | 284 | 284 | 284 | 284 |
| Hcaps00083_SRR25580021 | Mississippiense | 284 | 284 | 282 | 282 | 282 | 283 |
| Hcaps00041_SRR21059749 | Mississippiense | 285 | 285 | 285 | 285 | 285 | 285 |
| Hcaps00247_SRR6243638 | Mississippiense | 286 | 288 | 289 | 289 | 289 | 289 |
| Hcaps00243_SRR6243634 | Mississippiense | 287 | 289 | 288 | 288 | 288 | 288 |

| | | | | | | | |
|---|---|---|---|---|---|---|---|
| Hcaps00245_SRR6243636 | Mississippiense | 288 | 287 | 287 | 286 | 286 | 287 |
| Hcaps00244_SRR6243635 | Mississippiense | 289 | 286 | 286 | 287 | 287 | 286 |
| Hcaps00119_SRR25580057 | Mississippiense | 290 | 290 | 290 | 290 | 290 | 290 |
| Hcaps00257_SRR6243648 | Mississippiense | 291 | 292 | 291 | 292 | 292 | 292 |
| Hcaps00271_SRR6243662 | Mississippiense | 292 | 291 | 292 | 291 | 291 | 291 |
| Hcaps00250_SRR6243641 | Mississippiense | 293 | 293 | 294 | 294 | 293 | 293 |
| Hcaps00242_SRR6243633 | Mississippiense | 294 | 294 | 293 | 293 | 294 | 294 |
| Hcapsulatum_GCA017310585 | Mississippiense | 295 | 295 | 295 | 295 | 295 | 295 |
| Hcapsulatum_GCA000149585 | Mississippiense | 296 | 296 | 296 | 296 | 296 | 296 |
| Hcaps00141_SRR27481867 | India | 297 | 299 | 301 | 298 | 300 | 299 |
| Hcaps00150_SRR27481876 | India | 298 | 298 | 297 | 297 | 299 | 297 |
| Hcaps00158_SRR31797080 | India | 299 | 297 | 302 | 302 | 298 | 298 |
| Hcaps00146_SRR27481872 | India | 300 | 300 | 298 | 301 | 297 | 300 |
| Hcaps00140_SRR27481866 | India | 301 | 303 | 300 | 303 | 301 | 301 |
| Hcaps00152_SRR27481878 | India | 302 | 301 | 299 | 299 | 302 | 302 |
| Hcaps00147_SRR27481873 | India | 303 | 302 | 303 | 300 | 303 | 303 |
| Hcaps00145_SRR27481871 | India | 304 | 304 | 304 | 304 | 304 | 304 |
| Hcaps00139_SRR27481865 | India | 305 | 307 | 305 | 308 | 306 | 309 |
| Hcaps00143_SRR27481869 | India | 306 | 308 | 309 | 306 | 305 | 307 |
| Hcaps00138_SRR27481864 | India | 307 | 309 | 306 | 309 | 309 | 308 |
| Hcaps00151_SRR27481877 | India | 308 | 305 | 310 | 307 | 307 | 305 |
| Hcaps00149_SRR27481875 | India | 309 | 306 | 308 | 305 | 308 | 306 |
| Hcaps00142_SRR27481868 | India | 310 | 310 | 311 | 311 | 310 | 310 |
| Hcaps00148_SRR27481874 | India | 311 | 311 | 307 | 310 | 311 | 311 |
| Hcaps00137_SRR27481863 | India | 312 | 312 | 312 | 312 | 312 | 312 |
| Hcaps00144_SRR27481870 | India | 313 | 313 | 313 | 313 | 313 | 313 |

Position numbers are from top (#1) to bottom of trees.
wg = MLST scheme based on whole genome assemblies.
ann = MLST scheme based on annotations obtained via NCBI Genbank (Table S4).
fan = MLST scheme based on annotations generated with Funannotate (Table S4).
0300 = chewBBACA/pyrodigal setting of minimum ORF length of 300 nt.
0500 = chewBBACA/pyrodigal setting of minimum ORF length of 500 nt.

An Excel spreadsheet version of this Table is available at Zenodo (doi: 10.5281/zenodo.22865196) and Figshare (doi: 10.6084/m9.figshare.33946246).

**Table S6.** Linux command line instructions used in this study

All commands were run in Ubuntu 22.04.5 LTS, using dedicated Conda environments for sra-tools, Shovill, sra-tools, Quast, chewBBACA, grapetree and BUSCO, while seqfu was installed in all environments, and funannotate was used with a Docker image.

Download Illumina reads from NCBI SRA with sra-tools:

```
prefetch -p SRR31893254
fasterq-dump SRR31893254
pigz -p 14 *.fastq
rm -rf SRR31893254
```

Shovill genome assembly:

```
shovill --R1 SRRxxx_1.fastq.gz --R2 SRRxxx_2.fastq.gz --outdir SRRxxx_shovill --cpus 16 --minlen 500 --mincov 2.00 --ram 48.00 --depth 100 --nocorr --gsize 30.0M
```

Convert hyphens into underscores (required for ParSNP):

```
sed -i 's/\-/\_/' *.fna
```

Convert lowercase letters into uppercase letters (required for Quast):

```
sed -i 's/.*/\U&/' *.fna
```

Convert non-ACGT characters in all lines except for those starting with >:

```
sed '/^[^>]/ s/[^AGTC]/N/gi' < assembly.fna > assembly_edit.fna
```

Quast analysis:

```
quast.py --threads 10 --fast --no-check -m 0 -o quast_results/assembly_quast assembly.fna
```

Seqfu rename headers of FASTA files assemblies:

```
fu-multirelabel -b SRRxxxxxx.fna --comment-separator " " > HcapsXXXXXX_ SRRxxxxxx.fna
```

Extract gene sequences (exons+introns) from annotated genome:

```
agat_sp_extract_sequences.pl -g HcapsXXXXXX_ SRRxxxxxx.gff -f HcapsXXXXXX_SRRxxxxxx.fna -t gene -o HcapsXXXXXX_ SRRxxxxxx_genes.fasta
```

Create Pyrodigal training file with 300 and 500 nt minimum ORF length:

```
pyrodigal -g 1 -c -n -p single -t Histoplasma_capsulatum_0300.trn -i Hcapsulatum_GCA017310615.fna --min-gene 300
pyrodigal -g 1 -c -n -p single -t Histoplasma_capsulatum_0500.trn -i Hcapsulatum_GCA017310615.fna --min-gene 500
```

Create chewBBACA schemes:

```
chewBBACA.py CreateSchema -i histoplasma_genomes/ -o Hcaps_output_scheme_0300 --ptf Histoplasma_capsulatum_0300.trn --cpu 14 --t 1 --l 300
```

| |
|---|
| chewBBACA.py CreateSchema -i histoplasma_genomes/ -o Hcaps_output_scheme_0500 --ptf Histoplasma_capsulatum_0500.trn --cpu 14 --t 1 --l 500 |

Running chewBBACA allele calls:

| |
|---|
| chewBBACA.py AlleleCall -i histoplasma_genomes/ -g Hcaps_output_scheme_0300/schema_seed -o results_Histoplasma_scheme_0300 --cpu 14 --t 1 --l 300 --ptf Histoplasma_capsulatum_0300.trn |
| chewBBACA.py AlleleCall -i histoplasma_genomes/ -g Hcaps_output_scheme_0500/schema_seed -o results_Histoplasma_scheme_0500 --cpu 14 --t 1 --l 500 --ptf Histoplasma_capsulatum_0500.trn |

Make phylogenetic trees using outputs from chewBBACA allele calls:

| |
|---|
| grapetree -n 14 -p results_Histoplasma_scheme_0300/results_alleles.tsv -m RapidNJ > Histoplasma_capsulatum_scheme_0300.nwk |
| grapetree -n 14 -p results_Histoplasma_scheme_0500/results_alleles.tsv -m RapidNJ > Histoplasma_capsulatum_scheme_0500.nwk |

Funannotate (using Funannotate Docker image):

| |
|---|
| funannotate-docker clean -i Hcaps_GCA017310615.fna -o Hcapsulatum_GCA017310615_cleaned.fna --exhaustive --cpus 16 |
| funannotate-docker sort -i Hcaps_GCA017310615_cleaned.fna -o Hcaps_GCA017310615_sorted.fna -b GCA017310615 |
| funannotate-docker mask -i Hcaps_GCA017310615_sorted.fna -o Hcap_GCA017310615_masked.fna --cpus 16 |
| funannotate-docker predict -i Hcaps_GCA017310615_masked.fna -o Hcaps_GCA017310615_predict --species "Histoplasma capsulatum" --strain Hcaps_GCA017310615 --name GCA017310615 --busco_seed_species histoplasma_capsulatum --cpus 16 |
| funannotate-docker annotate -i Hcapsulatum_GCA017310615_funannotate -o Hcapsulatum_GCA017310615_funannotated -s "Histoplasma capsulatum" --cpus 16 |

BUSCO analysis:

| |
|---|
| busco -i Hcapsulatum_GCA017310615.fna -m genome -o busco_Hcapsulatum_GCA017310615 -c 16 -l ajellomycetaceae_odb12.2 |

A text version of this Table is available at Zenodo (doi: 10.5281/zenodo.22865196) and Figshare (doi: 10.6084/m9.figshare.33946246).
